# Multi-Omic, Multi-Tissue Responses to Acute Exercise in Sedentary Adults: Findings from the Molecular Transducers of Physical Activity Consortium

**DOI:** 10.64898/2026.02.27.702183

**Authors:** MoTrPAC Study Group, Daniel H. Katz, Christopher A. Jin, Gina M. Many, Gregory R. Smith, Hasmik Keshishian, Natalie M. Clark, Gayatri Iyer, Cheehoon Ahn, Malene E. Lindholm, Tyler J. Sagendorf, Nikolai G. Vetr, David Amar, Jacob L. Barber, Anna R. Brandt, Paul M. Coen, Yongchao Ge, Patrick Hart, Fang-Chi Hsu, Byron C. Jaeger, David Jimenez-Morales, Damon T. Leach, D. R. Mani, Shruti Marwaha, Samuel Montalvo, Hanna Pincas, Prashant Rao, James A. Sanford, Kevin S. Smith, Gaurav Tiwari, Anne Marie Weitzel, Joshua N. Adkins, Euan A. Ashley, Steven A. Carr, Michael E. Miller, Stephen B. Montgomery, Venugopalan D. Nair, Jeremy M. Robbins, Michael P. Snyder, Lauren M. Sparks, Russell Tracy, Martin J. Walsh, Matthew T. Wheeler, Ashley Y. Xia, Stuart C. Sealfon, Robert E. Gerszten, Scott Trappe, Charles F. Burant, Bret H. Goodpaster, Hiba Abou Assi, Mary Anne S. Amper, Brian J. Andonian, Isaac K. Attah, Alicia Belangee, Bryan C. Bergman, Daniel H. Bessesen, Sue C. Bodine, Gerard A. Boyd, Nicholas T. Broskey, Thomas W. Buford, Toby L. Chambers, Clarisa Chavez Martinez, Maria Chikina, Alex Claiborne, Zachary S. Clayton, Clary B. Clish, Katherine A. Collins-Bennett, Dan M. Cooper, Tiffany M. Cortes, Gary R. Cutter, Matthew Douglass, Sara E. Espinoza, Charles R. Evans, Facundo M. Fernandez, Johanna Y. Fleischman, Daniel E. Forman, Will A. Fountain, David A. Gaul, Catherine Gervais, Aaron H. Gouw, Kevin J. Gries, Marina A. Gritsenko, Fadia Haddad, Joshua R. Hansen, Trevor Hastie, Zhenxin Hou, Joseph A. Houmard, Kim M. Huffman, Ryan P. Hughes, Chelsea M. Hutchinson-Bunch, Olga Ilkayeva, John M. Jakicic, Catherine M. Jankowski, Pierre M. Jean-Beltran, Neil M. Johannsen, Johanna L. Johnson, Maureen T. Kachman, Erin E. Kershaw, Wendy M. Kohrt, William E. Kraus, Dillon J. Kuszmaul, Bridget Lester, Minghui Lu, Colleen E. Lynch, Nada Marjanovic, Sandra T. May, Edward L. Melanson, Nikhil Milind, Matthew E. Monroe, Cristhian Montenegro, Ronald J. Moore, Kerrie L. Moreau, Nicolas Musi, Masatoshi Naruse, Christopher B. Newgard, Bradley C Nindl, German Nudelman, Nora-Lovette Okwara, Eric A. Ortlund, Vladislav A. Petyuk, Paul D. Piehowski, David Popoli, Wei-Jun Qian, Shlomit Radom-Aizik, Tuomo Rankinen, Abraham Raskind, Alexander Raskind, Blake B. Rasmussen, Ulrika Raue, Eric Ravussin, R. Scott Rector, W. Jack Rejeski, Joseph Rigdon, Stas Rirak, Ethan Robbins, Margaret Robinson, Kaitlyn R. Rogers, Renee J. Rogers, Jessica L. Rooney, Irene E. Schauer, Robert S. Schwartz, Courtney G. Simmons, Chad M. Skiles, Tanu Soni, Maja Stefanovic-Racic, Cynthia L. Stowe, Andrew M. Stroh, Yifei Sun, Kristen J. Sutton, Anna Thalacker-Mercer, Robert Tibshirani, Todd A. Trappe, Mital Vasoya, Caroline S. Vincenty, Elena Volpi, Alexandria Vornholt, Katie L. Whytock, Angela Wiggins, Yilin Xie, Gilhyeon Yoon, Jay Yu, Xuechen Yu, Elena Zaslavsky, Zidong Zhang, Bingqing Zhao, Jimmy Zhen

## Abstract

Regular physical activity represents one of the most effective strategies for maintaining human health, yet the underlying molecular transducers of these benefits remain incompletely understood. Multi-omic assays provide new opportunities to study coordinated molecular responses of body tissues to different exercise modalities. The Molecular Transducers of Physical Activity Consortium (MoTrPAC) was established to address this need by creating a molecular map of the response to physical activity. Described here is the first human cohort of MoTrPAC: sedentary adults enrolled prior to study suspension during the COVID-19 pandemic (N=175) randomized to either endurance or resistance exercise, or non-exercise control. We detail the global acute molecular response in skeletal muscle, adipose tissue, and blood, integrated at multiple levels: tissue, exercise modality, timepoint, and omic category. These analyses characterize key molecular pathways, identify central regulators, and implicate novel candidate exerkines in mediating multi-organ exercise effects.

## Introduction

Physical activity is known to improve human health and prevent illness, with benefits extending to multiple domains, including cognitive, metabolic, cardiovascular, musculoskeletal, immune, gastrointestinal, and reproductive health.^1^ For health benefits to accrue with regular exercise, each acute bout of exercise initiates a cascade of molecular events that, when repeated over time, leads to the long-term effects of exercise training.^2,3^ However, the precise nature of the acute molecular response to exercise remains incompletely understood. Several molecular responses have been described, including flux in intracellular factors (e.g., ATP, NAD) in the contracting muscle and other tissues,^2,4^ provision of fuel for contracting muscle and protein accretion,^5^ and alterations in circulating signaling molecules (e.g., exerkines).^6^ Further, acute exercise effects are known to impact cellular behavior across molecular classes, including the methylome, the accessible chromatin landscape, the transcriptome, the proteome, the phosphoproteome, and the metabolome.^7^

While a number of studies have examined the epigenomic,^8–10^ transcriptomic,^11–16^ proteomic/phosphoproteomic,^17–21^ and/or metabolomic/lipidomic responses to acute exercise,^22,23^ the majority have focused on a single “ome” and a single tissue and are often limited to endurance-type exercise. Studies frequently examine skeletal muscle (SKM), with only a handful examining adipose tissue (AT)^9,24^ or blood.^25–29^ Thus, there is a critical need for an integrated multi-omic, cross-tissue, temporally resolved approach to elucidate the molecular responses to multiple exercise modalities. Such data can lead to the identification of novel therapeutic targets and inform more precise exercise recommendations for specific groups.^30^

The Molecular Transducers of Physical Activity Consortium (MoTrPAC), a National Institutes of Health Common Fund program, was created to build an extensive molecular “map” of exercise. MoTrPAC has previously released a multi-omic, multi-tissue dataset mapping the molecular response to endurance exercise training in rats.^31^ We now provide the first human data, derived from participants enrolled prior to study suspension during the COVID-19 pandemic. These generally healthy sedentary adult volunteers were randomized to undergo an acute bout of either endurance (EE) or resistance exercise (RE), or to remain supine as non-exercise controls (CON). Each provided multiple blood, muscle (vastus lateralis), and adipose (abdominal subcutaneous) samples for multi-omic profiling at multiple timepoints. These dense multi-tissue, multi-omic data provide the basis for an integrative map of the acute response to resistance and endurance exercise. Herein, we provide a global landscape of the multi-tissue response. A detailed account of the clinical intervention and physiological responses to exercise, companion works highlighting each individual tissue, and a detailed analysis of splicing variation are reported separately (ref Clinical Landscape in Review at Nature Medicine).^32–36^

## Results

### Study design and participant characteristics

We analyzed 175 generally healthy sedentary individuals who participated in an initial acute exercise bout (or were a resting control) and had at least one sample with biomolecular profiling.^37^ The trial components and acute exercise bout were standardized and performed across clinical centers and are described elsewhere (Methods, Figs. 1a, b) (ref Clinical Landscape in Review at Nature Medicine).^37^ In brief, participants first underwent baseline testing including anthropometrics, vitals, clinical labs, cardiopulmonary exercise testing (CPET), and strength testing. Participants were randomized in an approximate 8:8:3 ratio to EE, RE, or CON groups and simultaneously randomized to temporal biospecimen collection groups (Methods).

**Figure 1-.**
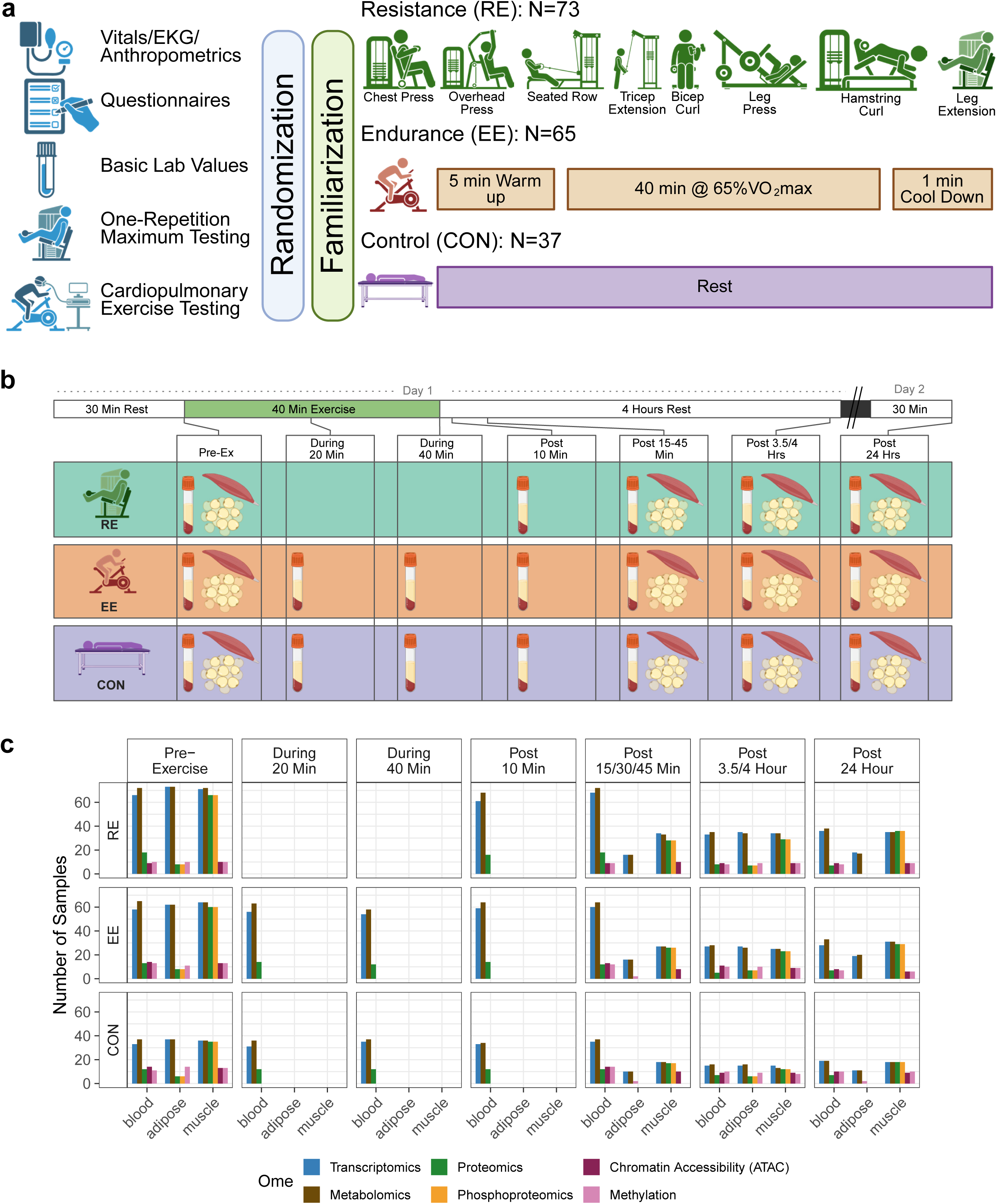
Study overview. **a**, Overview of the acute exercise bout. Baseline testing is completed before randomization. After randomization, a familiarization period for exercise is completed. Patients arrive in the morning, fasted, to complete their respective interventions. Created in https://BioRender.com. **b**, Schematic of muscle, blood, and adipose biospecimen collection. No samples are collected during resistance exercise. Created in https://BioRender.com. **c**, Number of samples measured in each omic category (on any assay) by tissue, exercise modality and timepoint. See Extended Data Fig 1 to see overlapping measurements for all participants.

Timepoint-matched non-exercising controls were deemed critical to control for the molecular effects of time of day, fasting, tissue sampling, and other non-exercise intervention stimuli. The majority of participants were female (72%), including a larger percentage (84%) of the CON group as compared to EE and RE (Table 1) (ref. Clinical Landscape in Review at Nature Medicine). Mean age was 41 ± 15 years, average body mass index (BMI) was 26.9 ± 4.0 kg/m^2^, and average waist circumference was 92 ± 12 cm. Baseline CPET testing by cycle ergometer showed an average VO_2_peak of 24 ± 7.0 ml/kg/min (Females: 22.2 ± 5.1 ml/kg/min; Males: 31.3 ± 7.1 mL/kg/min). Participant surveys and whole genome sequencing derived ancestry analysis showed that participants came from a variety of racial, ethnic, and genetic backgrounds (Table 1, Extended Data Fig. 1a).

**Table 1.** Participant Characteristics.

| Characteristic | Overall<br>N = 175 | EE<br>N = 65 | RE<br>N = 73 | CON<br>N = 37 |
| --- | --- | --- | --- | --- |
| **Demographics** |  |  |  |  |
| Sex |  |  |  |  |
| Female | 126 (72%) | 46 (71%) | 49 (67%) | 31 (84%) |
| Male | 49 (28%) | 19 (29%) | 24 (33%) | 6 (16%) |
| Age (yrs) | 41 (15) | 42 (14) | 40 (15) | 42 (15) |
| Height (cm) | 168 (9) | 168 (9) | 168 (10) | 167 (8) |
| Weight (kg) | 76 (14) | 76 (13) | 77 (14) | 73 (13) |
| BMI (kg/m<sup>2</sup>) | 26.9 (4.0) | 27.0 (4.0) | 27.2 (4.0) | 26.2 (3.9) |
| Waist Circumference (cm) | 92 (12) | 93 (11) | 93 (12) | 89 (13) |
| Systolic BP (mmHg) | 116 (12) | 117 (11) | 116 (12) | 114 (11) |
| Diastolic BP (mmHg) | 72 (8) | 73 (9) | 72 (8) | 72 (8) |
| Hemoglobin A1c (%) | 5.30 (0.30) | 5.29 (0.32) | 5.31 (0.32) | 5.29 (0.21) |
| eGFR (mL/min/1.73 m<sup>2</sup>) | 101 (20) | 103 (18) | 101 (22) | 99 (19) |
| **Cardiopulmonary Exercise Test (CPET)** |  |  |  |  |
| VO<sub>2</sub>peak (L) | 1.86 (0.65) | 1.90 (0.68) | 1.89 (0.65) | 1.73 (0.56) |
| VO<sub>2</sub>peak (mL/min/kg) | 24 (7) | 25 (7) | 25 (7) | 24 (6) |
| Workload (peak, Watts) | 152 (50) | 155 (53) | 153 (52) | 142 (40) |
| O<sub>2</sub> Pulse (peak, ml/beat) | 10.6 (3.3) | 10.9 (3.5) | 10.8 (3.4) | 10.0 (2.8) |
| RER (peak) | 1.16 (0.08) | 1.17 (0.07) | 1.15 (0.08) | 1.16 (0.08) |
| **Strength Tests** |  |  |  |  |
| Isometric Knee Extension (Nm) | 148 (57) | 145 (54) | 156 (60) | 137 (53) |
| Hand Grip (kg) | 30 (10) | 30 (10) | 31 (12) | 28 (7) |
| *Values are mean (SD) for continuous variables and n (%) for categorical variables.*<br>*Abbreviations: EE = endurance exercise; RE = resistance exercise; CON = control; BMI = body mass index; BP = blood pressure; eGFR = estimated glomerular filtration rate; CPET = cardiopulmonary exercise testing, and RER = Respiratory Exchange Ratio.* |  |  |  |  |

After randomization, sedentary participants assigned to EE or RE had two or three familiarization sessions that were completed 5–8 days before the acute exercise test (ref. Clinical Landscape in Review at Nature Medicine, Fig. 1a). On the acute exercise day, participants arrived in the morning following an overnight fast. Biospecimens for multi-omic analysis were collected from three tissues (blood, SKM, and AT) at baseline (Fig. 1b, Methods, Extended Data Fig. 1b). Biospecimen collection varied across group, tissue, and timepoint due to participant burden; application of omic assays also varied due to technical constraints (Fig. 1b-c, Extended Data Fig. 1b, Supplementary Table 1, Methods). For example, proteome and phosphoproteome profiling were performed at all timepoints available in SKM, but were limited to a single timepoint in AT. Thus, combinations of tissue-ome-timepoint have varying power to detect differences in molecular abundance, and results should be interpreted accordingly. To enhance transparency and generalizability, sample sizes for all features (Extended Data Fig. 1c) and for individual features specifically discussed in the manuscript are provided (Supplementary Results, Supplementary Table 1). For each tissue-ome combination (e.g., SKM metabolomics), principal components of variation, using all feature measurements, were correlated to technical and phenotypic variables (Supplementary Results, Extended Data Fig. 1d). Participant characteristics are further profiled in a separate manuscript (ref. Clinical Landscape in Review at Nature Medicine).

### Exercise-induced multi-omic changes

Exercise impacted thousands of molecular features, with significant variation between exercise modality, tissue, and ome (Fig. 2a, Supplementary Table 2). The inclusion of a non-exercising control group not only reduced false attribution of non-exercise effects (Methods), but also permitted identification of exercise-responsive molecules which did not change in absolute abundance, but were nonetheless affected by exercise (Extended Data Fig. 2a, b). For example, the core clock gene *PER2* was unchanged in SKM-RE relative to baseline across time points, but was significantly higher compared to CON at 3.5 hours post-exercise (Extended Data Fig. 2b), highlighting the utility of CON in interpreting the dynamic changes following exercise and alluding to circadian effects.

**Figure 2-.**
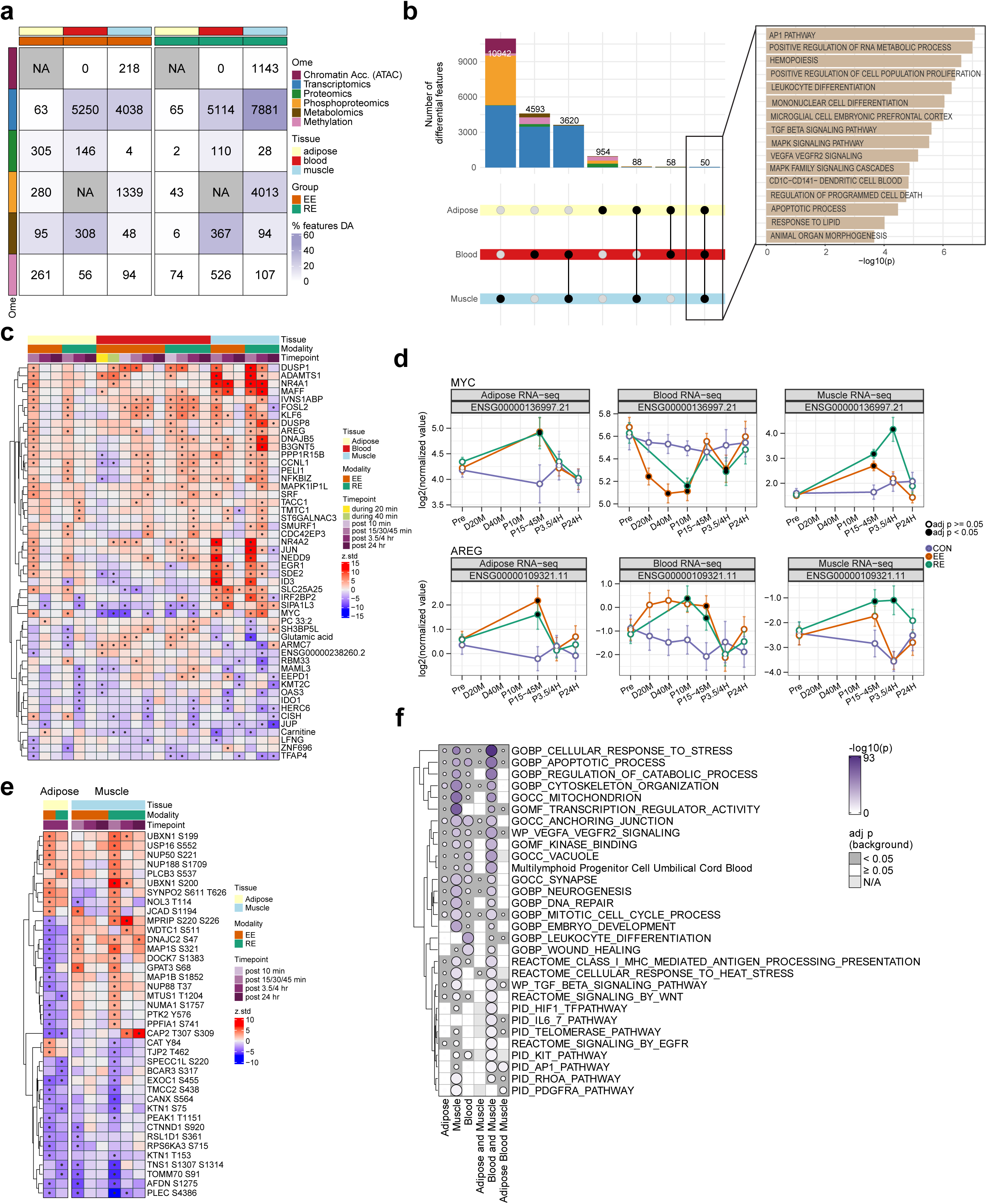
Exercise-induced multi-omic changes. **a**, Heatmap displaying the number (annotated in each cell) of differentially abundant (DA) features (adj. p-value < 0.05) by ome, tissue, and exercise modality combination at any timepoint. The color intensity of each box reflects the percentage of DA features relative to total assayed features per omic layer (see color bar), where gray indicates not analyzed. EE = Endurance Exercise; RE = Resistance Exercise. **b**, UpSet plot of DA features across all tissues, exercise modalities, and omes (left). Black dots with connected lines represent the intersection of DA omic features by tissue. Number of DA features of any ome are indicated on top of the bars. Over-representation analysis (ORA) of 47 shared DA transcripts between all three tissues (right). **c**, Heatmap of all 50 DA features (transcripts and metabolites) shared between all three tissues presented according to timepoint and exercise modality. Z-scores of data are used for the heatmap. (*) indicates adj. p-value < 0.05. **d**, Temporal trajectories of *MYC* and *AREG* transcripts that are DA in all three tissues. EE = endurance exercise, RE = resistance exercise, CON = control group, Pre = pre-exercise, D20M = during 20 minutes, D40M = during 40 minutes, P10M = 10 minutes post-exercise, P15-45M = 15, 30, or 45 minutes post-exercise (depending on tissue), P3.5-4H = 3.5 or 4 hours post-exercise (depending on tissue), P24H = 24 hours post-exercise. Data is shown as group-timepoint mean ± 95% confidence interval for the group mean with a black circle indicating model significance relative to control at adj. p-val < 0.05. Plots display Ensembl ID. **e**, Heatmap of 39 DA phosphosites shared between SKM and AT. Phosphoproteomics was only performed at the pre-exercise and 3.5 hour timepoint in the AT. Z-scores of data are used for the heatmap. (*) indicates adj. p-value < 0.05. **f**, Heatmap showing ORA of DA features in each tissue and for features with overlap between all three tissues and tissue pairs manually curated to show overlap across tissues using Molecular Signatures Database (MSigDB) C2 and C5 databases, and the CellMarker 2.0 database^126^. Pathways shown are significant in at least one of the groups (adj. p-val < 0.05, as indicated by the grey background of each cell). Notably, no pathway was unique to blood and adipose.

Across both exercise modalities, the majority of differentially abundant (DA) features were tissue-specific, with blood and SKM showing the greatest concordance (Fig. 2b). Fifty DA features were shared across all three tissues, with 47 being transcripts (Fig. 2b, Extended Data Fig. 2c). Over-representation analysis (ORA) of shared DA transcripts revealed pathway enrichments related to hemopoiesis, RNA metabolic processes, AP1, TGFβ, MAPK and VEGFR signaling, apoptosis, regulation of cell proliferation, and organ morphogenesis (Fig. 2b; Supplementary Table 2). Among these shared features, the direction and magnitude of exercise effects varied across tissues and modalities (Fig. 2c, Extended Data Fig. 2d, Supplementary Results). For example, the proto-oncogene *MYC*, which promotes VEGF-mediated angiogenesis,^38^ decreased in blood cells but increased in SKM and AT (Figs. 2c, d), whereas expression of the epidermal growth factor receptor agonist *AREG* increased across all three tissues (Figs. 2c, d, Supplementary Results).

Metabolomic effects were most prominent in blood, with substantial overlap between blood DA metabolites and those in SKM (n=70) and AT (n=47) (Extended Data Fig. 2c). All three tissues shared three DA metabolites: carnitine, glutamic acid, and PC 33:2 (Fig. 2c).

Despite limited phosphoproteomic profiling in the AT, 39 phosphosites were exercise-responsive in both SKM and AT, with 24 sharing directionality (Fig. 2e). For example, junctional protein associated with coronary artery disease (JCAD) s1194, which functions to promote angiogenesis, increased in SKM and AT (EE only),^39^ consistent with shared transcriptomic enrichments in VEGFR signaling. Conversely, GPAT3 S68, an enzyme involved in lipid synthesis,^40,41^ decreased with EE in AT and increased with both exercise modalities in SKM, supportive of altered substrate flux between AT and SKM in the early post-exercise response. Enrichment analysis by ORA against the PTMSigDB collection revealed shared enrichment of MAP2K2 in the AT and SKM (Extended Data Fig. 2e), which promotes cellular proliferation,^42^ and is further consistent with transcriptomic enrichments in cellular growth and differentiation terms (Fig. 2b).

Given the paucity of shared DA features across tissues, we further investigated tissue-concordant pathway enrichments via ORA of tissue-specific DA features. Pathways related to tissue regeneration (e.g., PDGFRA and mitotic cell cycle process), angiogenesis (e.g., VEGFR), neurogenesis, and inflammation (e.g., leukocyte differentiation) were enriched across all three tissues, while SKM and blood shared unique enrichments in wound healing (Fig. 2f, Supplementary Table 2). These shared pathway-level enrichments contrasted limited DA feature overlap across tissues (Fig. 2b and Extended Data Fig. 2c), suggesting that exercise induces conserved biological responses despite tissue-specific molecular regulation.

Given the cohort was 72% female, we performed a sensitivity analysis to ascertain the generalizability of our sex-corrected model by performing sex-specific DA analyses to determine feature-level differences by timepoint and group. Pearson’s correlation between the logFC in males and the logFC in females across the most consistently assessed omic layers (transcriptomics and metabolomics) revealed high concordance for features that were DA in the joint analysis (R = 0.929, Supplementary Table 2, Extended Data Fig. 3a). Examination of individual feature temporal trajectories also revealed sex concordance (Extended Data Figs. 3b,c). The most notable sex difference appeared in blood transcriptomics in the RE early post-exercise response, revealing a larger effect size in male participants (logFC correlation slopes: 10 minutes: 1.38; 30 minutes: 1.20; 3.5 hours: 1.31), warranting future validation and exploration. Subsequent pathway enrichment revealed more pronounced innate immune signaling enrichment in males, including neutrophil degranulation and secretory granule terms, as well as differential features involved in these inflammatory processes (e.g., *GZMB*, *CD177, MMP8)* (Extended Data Fig. 3a). As exercise is known to have substantial effects on blood cell populations (e.g., via neutrophil demargination),^43^ further sensitivity analyses were performed to assess the effect of cell-type shifts on DA results in blood transcriptomics. These analyses displayed moderate concordance with complete blood count parameters, which were available only at baseline. Accounting for estimated cell-type covariates during and post-exercise had minimal impact on the DA results (Extended Data Figs. 3d, e; Supplementary Results).

### Exercise modality-specific effects across tissues and omes

The number of molecules observed to be affected by each exercise modality differed substantially (Fig. 2a); RE led to a greater number of DA molecules in SKM and blood, with just 22% and 23% overlap with EE, respectively, although the EE group had two additional timepoints collected in the blood. In AT, EE drove changes in more molecules as compared to RE, with only 2% of EE DA molecules overlapping with RE (Fig. 3a and Extended Data Fig. 4a). EE and RE effects were then contrasted directly for all timepoints with data from both groups, such that for any given molecular feature, differences are described for EE vs. CON, RE vs. CON, as well as EE vs. RE (Fig. 3a). Overall, the majority of SKM DA features differed between modalities, while in AT and blood few molecules exhibited statistically significant modality differences when contrasted directly, despite the numerical differences in significant DA molecules from the EE or RE vs. CON contrasts in those tissues (Fig. 3a).

**Figure 3-.**
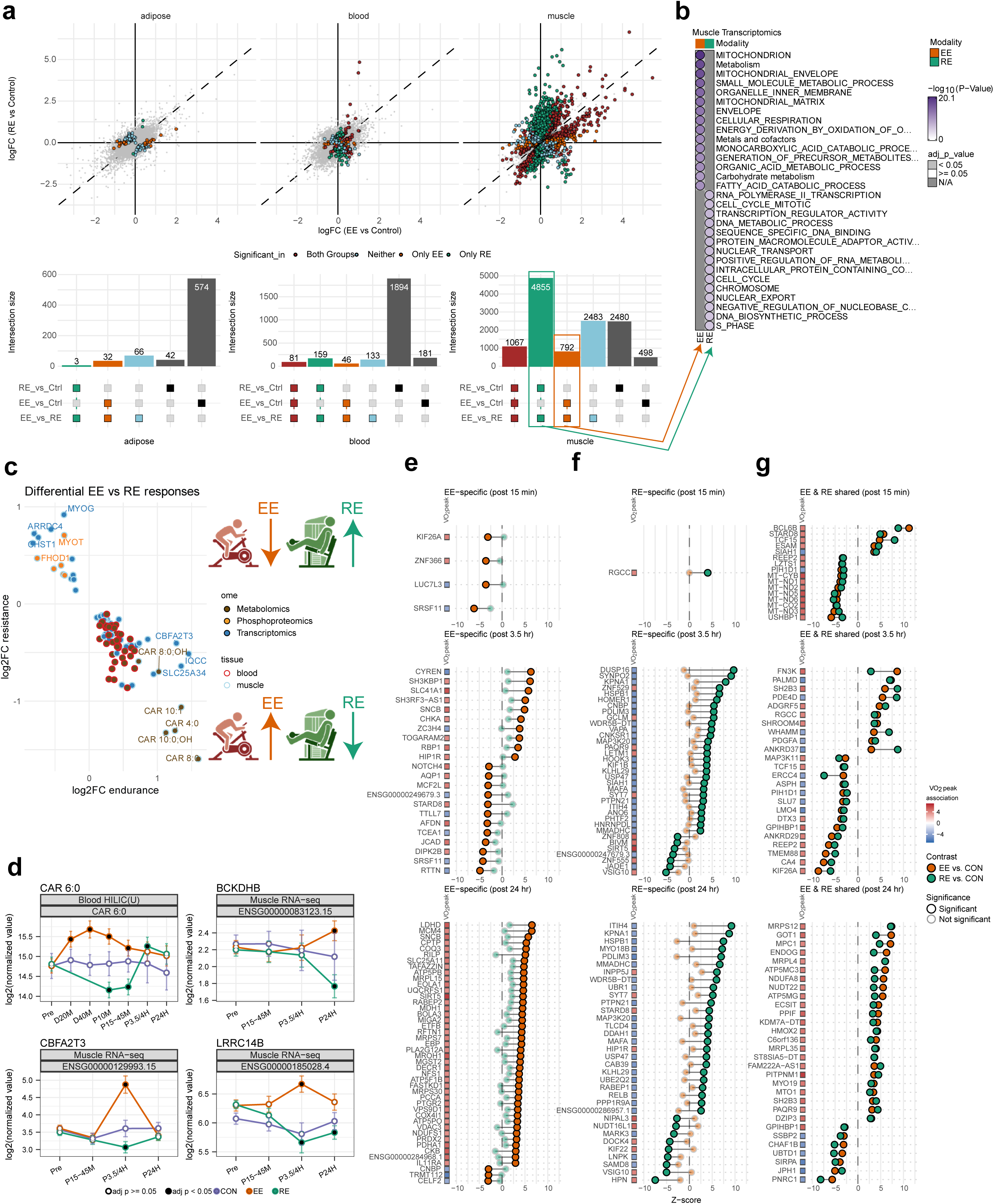
Differential responses to endurance and resistance exercise. **a**, Top panel: For all measured features, the log fold change (logFC) for EE vs. CON (x-axis) and RE vs. CON (y-axis) are plotted by points. Molecules which are significant (adj. p-value < 0.05) in the EE vs. RE comparisons and EE/RE vs. CON comparisons are indicated by colored points, colored by contrasts (EE-specific = orange, RE-specific = green), whereas features only significant in the EE vs. RE contrast are colored in light blue. All features which are non-significant in the EE vs. RE comparison are indicated in light grey. For each feature, only the timepoint with the most significant difference between EE and RE is included to improve visualization. Bottom panel: UpSet plots display the intersection of features included in the top panel significant in EE vs. RE, EE vs. CON, and RE vs. CON comparisons. Colored intersection labels match the colors described in the top panel. **b**, Overrepresentation analysis of muscle transcriptomic features from panel A specific to each exercise modality (green and orange points) using C2 and C5 MSigDB database. No enrichments were shared between the EE-specific (orange) features and RE-specific (green) features. Top 15 terms ranked by p-value are shown. **c**, Oppositely regulated features in the two exercise modalities (relative to control, adj. p-val < 0.05). Inclusion requires features are also significant in a direct comparison between EE and RE (adj. p-val<0.05). Shown are blood and SKM (no features met these criteria in AT). **d**, Timewise abundance changes of 4 specific features showing modality-divergent regulation in the blood or SKM. Pre = pre-exercise, D20M = during 20 minutes, D40M = during 40 minutes, P10M = 10 minutes post-exercise, P15-45M = 15, 30, 45 minutes post-exercise (depending on tissue), P3.5-4H = 3.5 or 4 hours post-exercise (depending on tissue), P24H = 24 hours post-exercise. Data is shown as group-timepoint mean ± 95% confidence interval with a black circle indicating significance relative to control at adj. p-val<0.05. CAR 6:0 = C6 Carnitine, BCKDHB = Branched Chain Keto Acid Dehydrogenase E1 Subunit Beta, CBFA2T3 = CBFA Partner Transcriptional Co-Repressor 3, LRRC14B = Leucine Rich Repeat Containing 14B. Plots indicate Ensembl for RNA or Uniprot IDs for proteins where appropriate. HILIC(U) = Hydrophilic Interaction Liquid Chromatography (Untargeted) metabolomics platform. **e-g**, Dumbbell plots showing baseline muscle transcriptomic-VO_2_peak associations and response to acute exercise. The top 70 genes, ranked by t-statistic for association with VO_2_peak, are shown, Genes are grouped as EE-specific (**e**), RE-specific (**f**), or EE-RE shared (**g**) responses at 15 min, 3.5 hours, and 24 hours post-exercise. Colored tiles adjacent to gene names indicate strength and direction of association of each gene with VO_2_peak, with red indicating positive and blue indicating negative associations. Points show post-exercise Z-scores for EE (orange) and RE (green). Statistically significant points are indicated by a black border.

In SKM, 4855 features were RE-specific, whereas 792 were EE-specific (Fig. 3a). ORA of SKM modality-specific features demonstrated EE enrichments for mitochondrial, carbohydrate, and fat metabolism pathways, versus RE enrichments for cell cycle, nuclear transport and DNA biosynthesis (Fig. 3b). Detailed enrichment analysis by timepoint showed the HSF1 pathway, a primary cellular response to increase expression of heat shock proteins, to have modality-specific activation pattern with the *HSPH1* transcript showing particularly strong temporal divergence between modalities, rising early post-exercise in EE but late after RE, consistent with overall pathway activation patterns (Supplementary Results, Extended Data Fig. 4b - 4e).

To explore the greatest modality differences, we identified molecules with divergent regulation (e.g., upregulated in EE vs. CON and downregulated in RE vs. CON), which revealed 38 features in blood, 71 in SKM, and 0 in AT (Fig. 3c; Supplementary Table 3). In blood, only metabolites were divergent (n=38); all increased in EE and decreased in RE (e.g., CAR 6:0, Fig. 3d), while SKM displayed divergent features across the transcriptome, metabolome, and phosphoproteome (Extended Data Fig. 4f). In blood and SKM, two acylcarnitines (CAR 10:1 and CAR 11:0) increased in response to EE and decreased with RE early post-exercise, indicating accelerated fatty acid metabolism following EE (Fig. 3c).^44,45^ SKM modality-divergent features further reflected differences in acute metabolic responses to EE and RE. For example, expression of the *HIF1A* transcriptional corepressor *CBFA2T3* (Fig. 3d) increased in EE and decreased in RE, and several key *HIF1A* glycolytic response genes (e.g., *LDHA, PDK4,* and *PFKP*) were upregulated with RE (data not shown). This, paired with acylcarnitine changes and modality differences in the peak lactate response (Clinical Landscape in Review at Nature Medicine), supports modality-divergent mechanisms for substrate utilization. Amino acid metabolism was also oppositely regulated in SKM, with *BCKDHB*, a subunit of the rate-limiting branched chain amino acid metabolism BCKDH complex, increasing in response to EE and decreasing in response to RE (Fig. 3d). Finally, *LRRC14B* – a gene with minimal experimental data, but which was recently identified as a regulator of striated muscle function and genetically associated with cardiac dysfunction – ^46,47^ increased in EE and decreased in RE (Fig. 3d). Taken together, these results highlight molecules that define the divergent phenotypic responses to EE and RE, making them critical targets for future investigation.

### Association between molecular features and clinical traits

Fitness, cardiometabolic, and clinical traits adapt differently to specific exercise modalities (e.g., VO_2_peak to EE, or strength to RE; ref Clinical Landscape in Review at Nature Medicine). To gain insight into the molecular underpinnings, we identified molecules that were both associated with such traits at baseline and also responsive to acute exercise.

At baseline, the bulk of associations were between insulin resistance (as measured by Homeostatic Model Assessment for Insulin Resistance, HOMA-IR) and both AT transcripts (n=2293) and blood metabolites (n=141). VO_2_peak was associated with numerous SKM transcripts (n=946) and SKM proteins (n=493, Extended Data Fig. 4g, Extended Data Table 1, Supplementary Results, Supplementary Table 3). Few features showed baseline associations with strength metrics, consistent with prior work,^48,49^ though given sample size, power in these analyses was lower (Methods).

In the subset of SKM transcripts associated with baseline VO_2_peak, ORA showed that transcripts with a positive association with baseline VO_2_peak were strongly enriched for mitochondrial oxidative phosphorylation (OXPHOS) terms, whereas transcripts with a negative association with VO_2_peak were enriched for sarcomeric organization, calcium handling, and translational and post-transcriptional regulation terms (Supplementary Table 3).

We next analyzed how these VO_2_peak associated SKM transcripts responded to acute exercise. We hypothesized that if a gene shows concordant expression trends with both baseline VO_2_peak and the response to acute exercise, it may contribute to long-term cardiorespiratory fitness adaptation. Transcripts were divided according to their acute exercise response in each modality – EE-specific, RE-specific, and EE-RE shared – and timepoint (Fig. 3e-g).

At the early and middle timepoints, the observed patterns were variable. At 15 minutes post-exercise, several mitochondrial-encoded OXPHOS genes positively associated with baseline VO_2_peak were downregulated after both exercise modalities, while upregulated genes were related to reparative processes. At 3.5 hours post-exercise, the EE-specific response showed no consistent relationship to the baseline VO_2_peak association (15 of 30 genes concordant), and the most coherent signal after either modality was transient suppression of endothelial genes positively associated with VO_2_peak, accompanied by induction of the hypoxia-responsive *ANKRD37*.^50^

A much sharper pattern emerged at 24 hours post-exercise. The effect of acute exercise on EE-specific transcripts and EE-RE shared transcripts was predominantly concordant with the baseline VO_2_peak association – acute exercise upregulated transcripts with a positive baseline association and downregulated transcripts with a negative baseline association (Fig. 3e,g). Most of the concordant genes were related to metabolism and included OXPHOS subunits (*NDUFS1, COX4I1, ATP5PB, ATP5F1B, ATP5MG, ATP5MC3, NDUFA8*), regulators and components of mitochondrial gene expression (*FASTKD1, MRPS7, MRPS12, MRPL4, MRPL15, MTO1*), components of the malate-aspartate shuttle (*GOT1*, *MDH1*, *SLC25A11*), and genes involved in mitochondrial dynamics and membrane remodeling (*MIGA2, TAFAZZIN, PPIF*), together suggesting a concordant mitochondrial remodeling response to acute exercise. EE additionally induced the expression of genes related to mitochondrial biogenesis and substrate transport.

Conversely, RE-specific transcripts showed predominantly discordant responses at both 3.5 and 24 hours post-exercise (Fig. 3f). At 24 hours, RE-upregulated transcripts included genes involved in myofibrillar remodeling (*SYNPO2, PDLIM3, MYO18B, HOMER1*), proteostasis (*SIAH1, KLHL29, USP47, UBE2Q2*), inflammation (*RELB*), and mechanical-stress signaling (*MAP3K20/*ZAK, whose muscle-enriched isoform is activated at the Z-disc by mechanical load).^51^

Taken together, while these findings alone do not directly establish molecular mediation, they do nominate a set of transcripts which may transduce acute exercise effects to long-term cardiorespiratory adaptation, while identifying RE-specific transcripts linked to muscle remodeling and less likely to support cardiorespiratory adaptation.

### Temporal molecular responses with acute exercise

Acute exercise initiates a coordinated, temporal cascade across multiple tissues.^27^ We aimed to capture these behaviors across tissues at multiple levels: for each timepoint in each tissue (simple tissue-ome-timepoint enrichment), for molecular trajectories in each tissue (c-means clustering), and for coordinated molecular trajectories across tissues (pathway-level information extractor [PLIER], Methods).

Of the hundreds of pathways identified by enrichment analysis by timepoint (Supplementary Table 4), shared enrichments across all three tissues emerged for the transcriptome (Fig. 4a), proteome (Fig. 4b), and metabolome (Extended Data Fig. 5a). Consistent with our previous timepoint-agnostic ORA analysis of DA features (Fig. 2b), VEGF, IL-6, AP1, and EGFR (e.g., ERBB1) signaling, OXPHOS, and circadian clock genes emerged as top enriched transcriptomic pathways across all three tissues and revealed greater clarity on modality and tissue-specific temporal trajectories. For example, in SKM and blood, RE induced prolonged transcriptomic enrichments of the angiogenic PDGFRB and VEGF pathways (Fig. 4a). At 24 hours, OXPHOS transcriptomic enrichments were greater with EE than RE, whereas at the protein level RE demonstrated a stronger suppression in OXPHOS proteins as compared to EE (Fig. 4b). Differences in protein turnover and translation may contribute to this discrepancy. Both modalities increased SKM proteasome protein enrichments at 24 hours, whereas RE induced sustained suppression of mitochondrial ribosome proteins throughout recovery. The greater EE-induced transcription of OXPHOS genes, coupled with suppression of autophagy genes at 3.5 hours, may offset proteasome-based clearance (Fig. 4a-b) and enable endurance training-mediated gains in OXPHOS.

**Figure 4.**
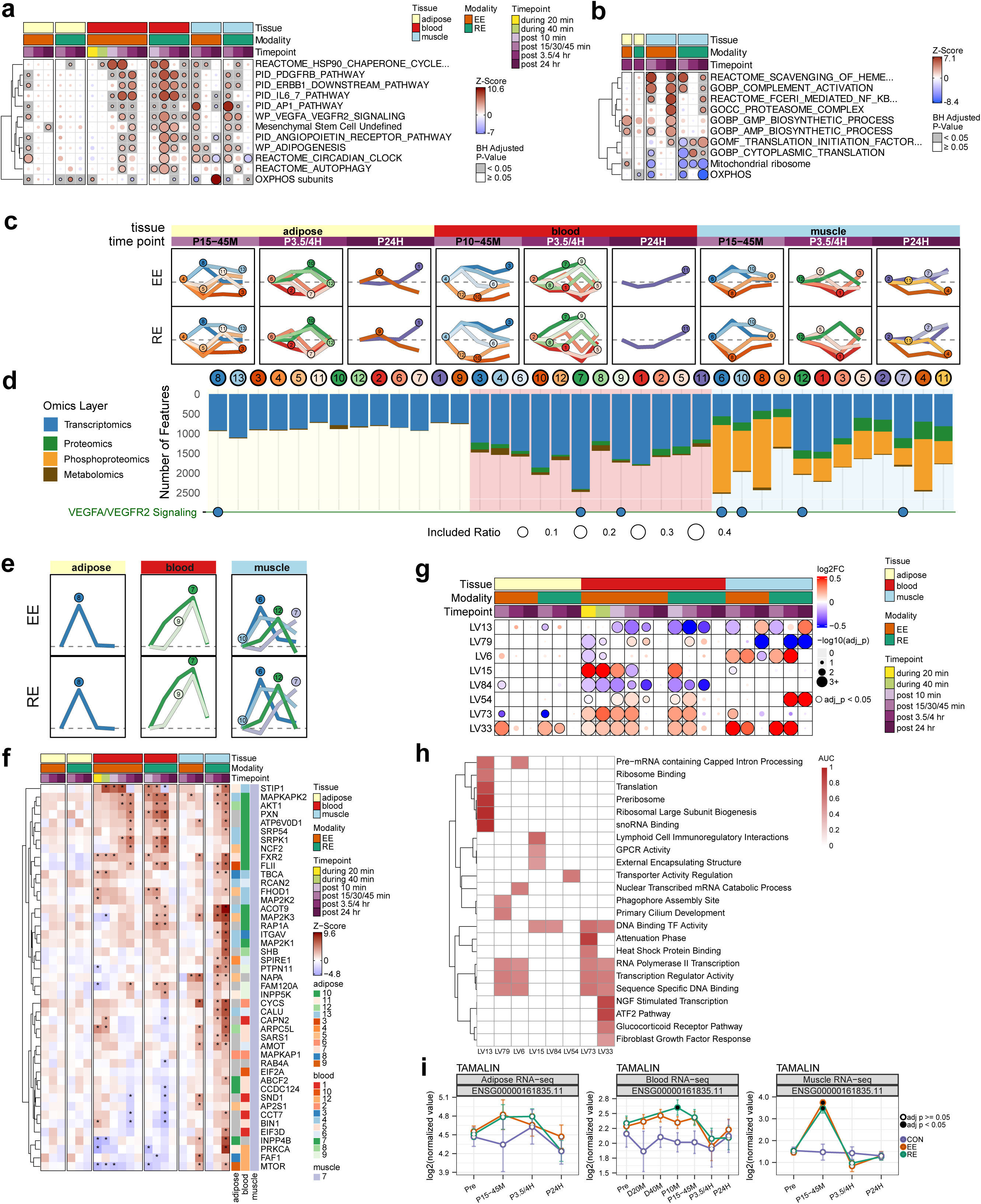
Temporal mapping of the molecular response to exercise across tissues. **a,b**, Pathway Enrichment for each tissue-timepoint in transcriptome (a) and proteome (b) utilizing Pre-ranked Correlation-Adjusted MEan-RAnk (CAMERA-PR). Bubbles indicating pathway enrichment are colored according to z-score and a black circle with gray background indicates significance (adj. p-value < 0.05). **c**, Temporal trajectories (sparklines) of all cluster centroids identified by fuzzy c-means clustering. Each line represents the average Z-score profile of a single cluster. Line color indicates the cluster, to be matched to the colored circles below. Clusters are organized by tissue and then by timepoint of maximal response (maximal absolute deviation [MAD] from zero); light purple indicates Early (15-45 minutes [and/or 10 minutes for blood]), medium purple indicates Mid (3.5-4 hours), and dark purple indicates Late (24 hours). Plots are split top and bottom by exercise modality (Endurance, EE; Resistance, RE). **d**, Omics composition of hard-clustered features. Each column represents a distinct cluster, ordered by tissue and temporal profile. The bar chart shows the total number of features assigned to each cluster (with a membership probability > 0.3), with bar segments colored by their omics layer of origin. Circles at the top display the cluster ID, with fill color corresponding to the temporal trajectory profile shown in (c). Background shading indicates the tissue of origin (adipose, blood, or muscle). **e**, Sparklines for c-means cluster trajectories which are enriched for VEGF pathway features. These are identical to the trajectories in (c) and are reprinted for clarity. **f**, Multi-tissue heatmap of Z-scores for transcriptomic features belonging to the VEGFA-VEGFR2 signaling pathway and cluster muscle-7. Columns represent timepoints grouped by tissue and exercise modality. The right-hand annotation indicates the c-means cluster assignment for each gene in adipose, blood, and muscle, with colors corresponding to the cluster profiles in (c-f). **g**, Bubble heatmap from the PLIER analysis showing the significance (size = adjusted p-value) and directionality (color = log2 fold-change) of the EE or RE response in the transcriptomic data in each tissue described by a subset of LVs with the most significant identified transcriptomic exercise responses. **h**, Top enriched pathways for the RNAseq LVs described in (g). Scale is the AUC of a statistical test of association between pathway and LV associated genes. The top five significant pathways are included for each LV, if they exist. **i**, Feature plots of the *TAMALIN* response to acute exercise in adipose, blood, and muscle.

To better understand multi-omic trajectories, we analyzed temporal molecular signatures across the post-exercise timepoints using fuzzy c-means clustering. To balance dimension reduction with specificity, we identified 13 clusters in AT and 12 in blood and SKM (Fig. 4c, see Methods, Supplementary Results) representing exercise response features across omes (Fig. 4d). ORA of features assigned to each cluster revealed more nuanced common pathway enrichments (Extended Data Fig. 5b, Supplementary Table 4, Supplementary Results). The VEGF signaling pathway, critical to the regenerative properties of exercise,^52–54^ exemplifies this temporal diversity, as seven temporal clusters were enriched with VEGF signaling features, including four within SKM, spanning the temporal spectrum (Fig. 4d-e, Extended Data Fig. 5b). Notably, three SKM clusters (10, 12, and 7) were elevated at 3.5 hours but followed distinct trajectories, highlighting the importance of trajectory-level analysis over single timepoint comparisons. Within each tissue, cluster membership is non-overlapping, but between tissues we observed modest overlap (Extended Data Fig. 5c), consistent with global analyses (Fig. 2b, Extended Data Fig. 2c).

Exploring known VEGF pathway genes within these trajectories, several novel exercise-responsive genes were identified including *SDCBP* and *HLX* (muscle cluster 12) which demonstrated peak expression in blood and muscle at 3.5 hours, as well as *INPP4B* and *AMOT* (muscle cluster 7) which were elevated at 24 hours (Fig. 4f, Extended Data Fig. 5d-e, Supplementary Results). Muscle cluster 12 and blood clusters 7 and 9 also contained enrichments for neurogenesis and autophagy, suggesting an overall tissue-regenerative exercise response. *NAPA* and *AP2S1*, also novel exercise genes, were upregulated at 24 hours. Their role in vesicular trafficking and endocytosis suggests this may be a key mechanism for translating acute exercise into sustained adaptation.^55–57^

To characterize coordinated cross-tissue molecular responses to exercise using latent variables (LVs), we applied the PLIER method (Extended Data Fig. 6a) to both the transcriptomic and metabolomic data (Fig. 4g, Supplementary Results, Extended Data Fig. 6a-i, Supplementary Table 4). At 15 to 45 minutes after exercise, we observed transcriptomic upregulation of several key transcription factors and response regulators in all three tissues within one of the identified LVs, LV33 (Extended Data Fig. 6d). LV33 included *NR4A1 and NR4A2*, known regulators of SKM adaptation,^58^ including immune cell activation–a key process in muscle regeneration.^59,60^ LV33 also included *ADAMTS1,* an anti-angiogenic regulator with well-described actions in muscle,^61^ here found to also be induced in AT and blood, suggesting multi-tissue control of angiogenesis (Extended Data Fig. 5d, Extended Data Fig. 6d, Supplementary Table 4). LV33 was further enriched for the neurotrophin nerve growth factor (NGF), which promotes neurogenesis and activates immune responses.^62,63^ While NGF and other neurotrophin family genes are responsive to exercise, *TAMALIN* emerged as a novel exercise-responsive neuroregulatory gene in LV33 (Fig. 4i). Functioning as a neuronal scaffold protein, *TAMALIN* promotes postsynaptic signaling and neuron survival.^64,65^ Altogether, findings from PLIER suggest a coordinated angiogenic and neurogenic response to acute exercise in the blood and SKM (Extended Data Fig. 5b), and warrant further investigations into the potentially novel role of *TAMALIN*.

PLIER applied to the cross-tissue metabolomic data highlighted differences in substrate flux in response to EE or RE (Supplementary Results, Extended Data Fig. 6e-i). Represented by LV17, acylcarnitines—intermediates of fatty acid oxidation—increased after EE but decreased after RE in both blood and SKM (Extended Data Fig. 6f,i).^66^ Consistent with this observation, EE induced specific upregulation of key fatty acid oxidation genes *HADHA*, *MLYCD*, and *ACAA2* (Supplementary Table 2), a finding explored in more detail in a muscle-specific companion paper,^34^ and consistent with clinical measures of fat oxidation (ref Clinical Landscape in Review at Nature Medicine).

### Transcription factor response in acute exercise

We next sought to identify putative transcription factors (TFs) underlying tissue-specific transcriptomic responses to exercise. We performed TF binding motif enrichment analysis and identified motifs enriched among exercise-responsive genes, which point to increased TF function and downstream effects. Application of Hypergeometric Optimization of Motif EnRichment (HOMER)^67^ to the differentially expressed genes (DEGs) revealed the majority of motif enrichments were identified within the larger DEG sets observed in SKM and blood (Fig. 5a and 5b; Supplementary Table 5). Motifs in the specificity protein (SP), Kruppel-like factor (KLF), and interferon regulatory family (IRF) TF families were the most consistently enriched across tissues, with SKM and blood displaying KLF and SP enrichment at a majority of timepoints measured (Fig. 5c). These TFs have various roles in tissue physiology, including regulation of metabolism, adipogenesis, and angiogenesis.^68–70^ In the AT, both pro- and anti-adipogenic KLF family members were enriched immediately following EE,^71^ followed by enrichment of the binding motif for the adipogenic TF RXR,^72^ suggesting acute effects of EE on adipogenic potential. At 3.5-4 hours, interferon-stimulated response elements (ISRE) and IRF TFs were enriched with both modalities in the AT and blood, as well as in SKM with EE. The circadian CLOCK TF motif was enriched in SKM and blood with RE only. Unique to blood were RUNX1 and ETV2, which are both involved in hematopoiesis, among other functions.^73^

**Figure 5-.**
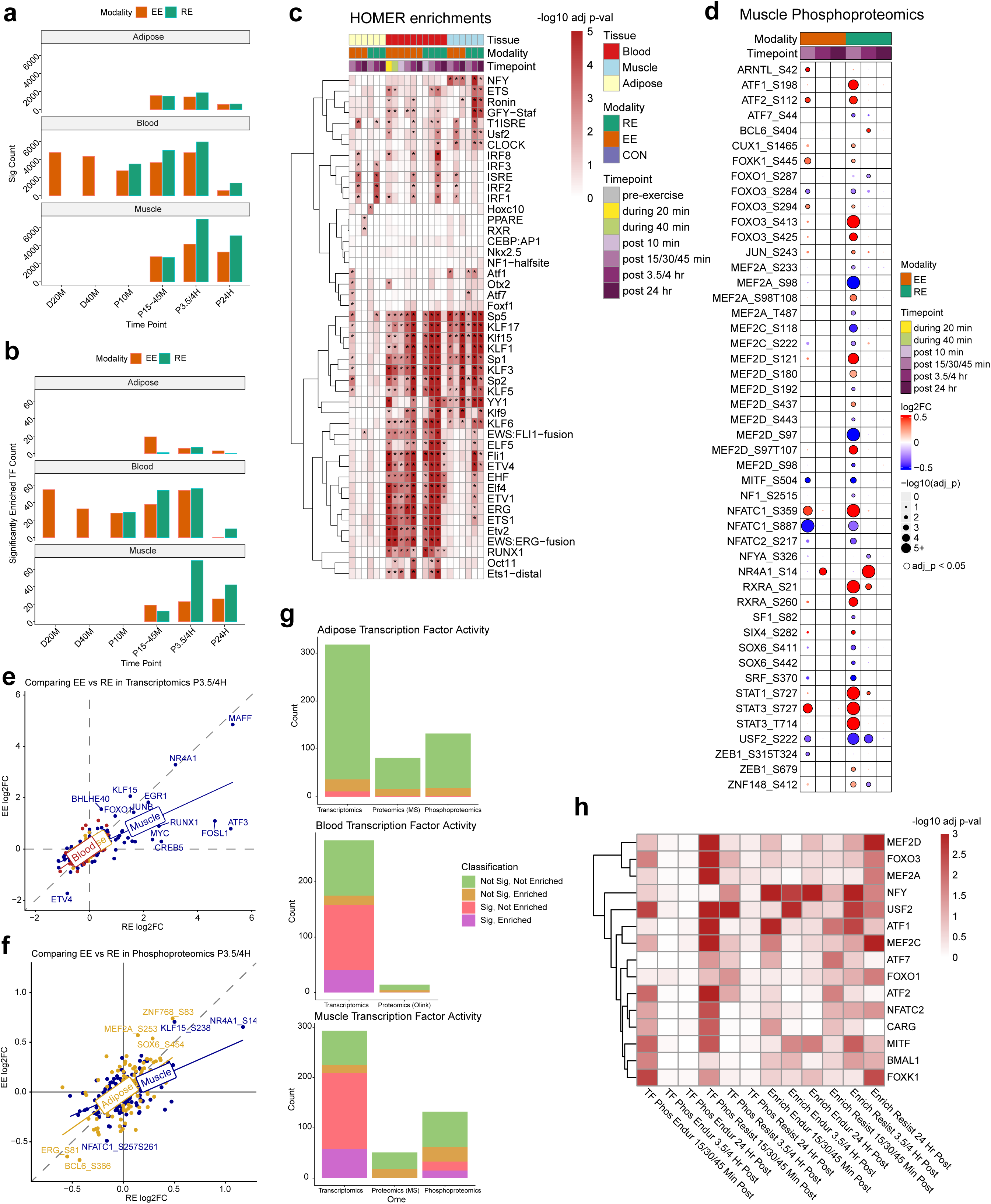
Transcription factor analysis. **a**, Barplot of counts of significant (nominal p-val < 0.05) genes for each tissue-modality-timepoint combination used as input for transcription factor (TF) enrichment analysis. **b**, Barplot of counts of significantly (adjusted p-val < 0.05) enriched TFs for the inputted significant genes for each tissue, modality, timepoint combination. **c**, Heatmap of selected top enriched TFs for each tissue, modality and timepoint combination. Heatmap entries show −log10 adjusted p-values of TF motif enrichment and TFs are hierarchically clustered by similar enrichment significance across conditions. *’s reflect motif enrichments with adj p-val < 0.05. **d**, Bubble heatmap of the most significant, adjusted p-val < 0.05, TF phosphoproteomic responses to acute exercise in muscle. **e-f**, Scatter plots of the L2FC (log2 fold change) of transcriptomic (**e**) and phosphoproteomic (**f**) responses to EE vs RE for all three tissues, a linear model used to generate the fit line for each tissue. **g**, Stacked barplots for each tissue and ome reflecting the proportion of TFs with data in that tissue/ome combination that can be classified as significantly responding to acute exercise and/or significantly enriched for target genes significantly responding to acute exercise. **h**, Heatmap of select TFs which exhibited significant muscle phosphoproteomic responses to exercise and were enriched for target significant genes in muscle. The first six columns from the left represent the significance of the TF phosphoproteomic response to either EE or RE at each post-exercise timepoint. The last six columns represent the significance of enrichment for the TF among the significant genes responding to either EE or RE at each post-exercise timepoint. Endur = Endurance, Resist = Resistance.

As a corollary to increased TF function via motif enrichment, TFs were also likely to be differentially expressed during acute exercise; 72% of identified TFs were themselves DEGs in SKM, 57% in blood, and 3% in AT. Nine TFs were among the 47 transcripts that exhibited significant expression increases at 15-45 minutes post-exercise in all three tissues (Fig. 2c, Extended Data Fig. 7): these included *NR4A1*, *JUN*, *MAFF*, *FOSL2, KLF6*, and *SRF* (also identified in PLIER analysis, Supplementary Table 4 [LV33]). *TFAP4* decreased post-exercise in all tissues, while previously discussed *MYC*, alongside *EGR1*, increased in both AT and SKM, but decreased in blood (Extended Data Fig. 7). MYC and EGR1 have been shown to functionally interact during cancer treatment to promote tumor cell death,^74^ suggesting their interaction may extend to exercise response and have pleiotropic effects. Phosphorylation also regulated TF activity, with 48 DA phosphosites identified across 29 distinct TFs in skeletal muscle—predominantly at 15 minutes post-resistance exercise (Fig. 5d). Transcriptomic and phosphoproteomic changes were more skewed towards RE in SKM relative to other tissues (Fig. 5e,f).

Combining all three levels of TF regulation (Fig. 5g), we identified 15 TFs in SKM that were differentially expressed, differentially phosphorylated, and had downstream motif-enrichment in response to exercise. In examining those 15 TFs, we identified timescale patterns in TF and target activity. Phosphorylation changes tended to occur early, at 15 minutes post-exercise in SKM, whereas motif enrichment occurred at later post-exercise timepoints (Fig. 5h), as demonstrated by the MEF2 and FOXO TFs with motif enrichment peaking 24 hours post-exercise. This pattern was also found with USF2, MITF, NFATC2, and circadian-regulatory BMAL1. NFY motif enrichment occurred broadly across all post-exercise timepoints, with a higher significance following EE than RE. Conversely, both the phosphorylation patterns and motif enrichment of the ATF family of TFs and the CArG motif occurred at 15 minutes post-exercise, suggesting a more rapid acting TF response. Taken together, these analyses reveal multi-tissue and tissue-specific TF responses to acute exercise, with SKM showing direct connections between phosphorylation changes and motif enrichment, providing insight into the temporal scales of TF activity.

### Multi-omic regulatory networks reveal TFEB as an exercise response hub

While transcription factors are often key regulators of biological responses, metabolites and other molecules can play central roles in coordinating signaling events and downstream physiological responses.^75,76^ Using Spatiotemporal Clustering and Inference of Omics Networks (SC-ION)^77^ to define regulatory networks (Fig. 6a), we identified multiple metabolites and TFs as putative central regulators. Findings included NFYA, a subunit of the NFY TF also identified by HOMER (Supplementary Table 6). Others not identified by HOMER included PITX3, a protein expressed in SKM and important in myogenic regulation,^78^ and STAT5B, a key regulator of myogenic IL-15 and growth hormone signaling.^79^

**Figure 6-.**
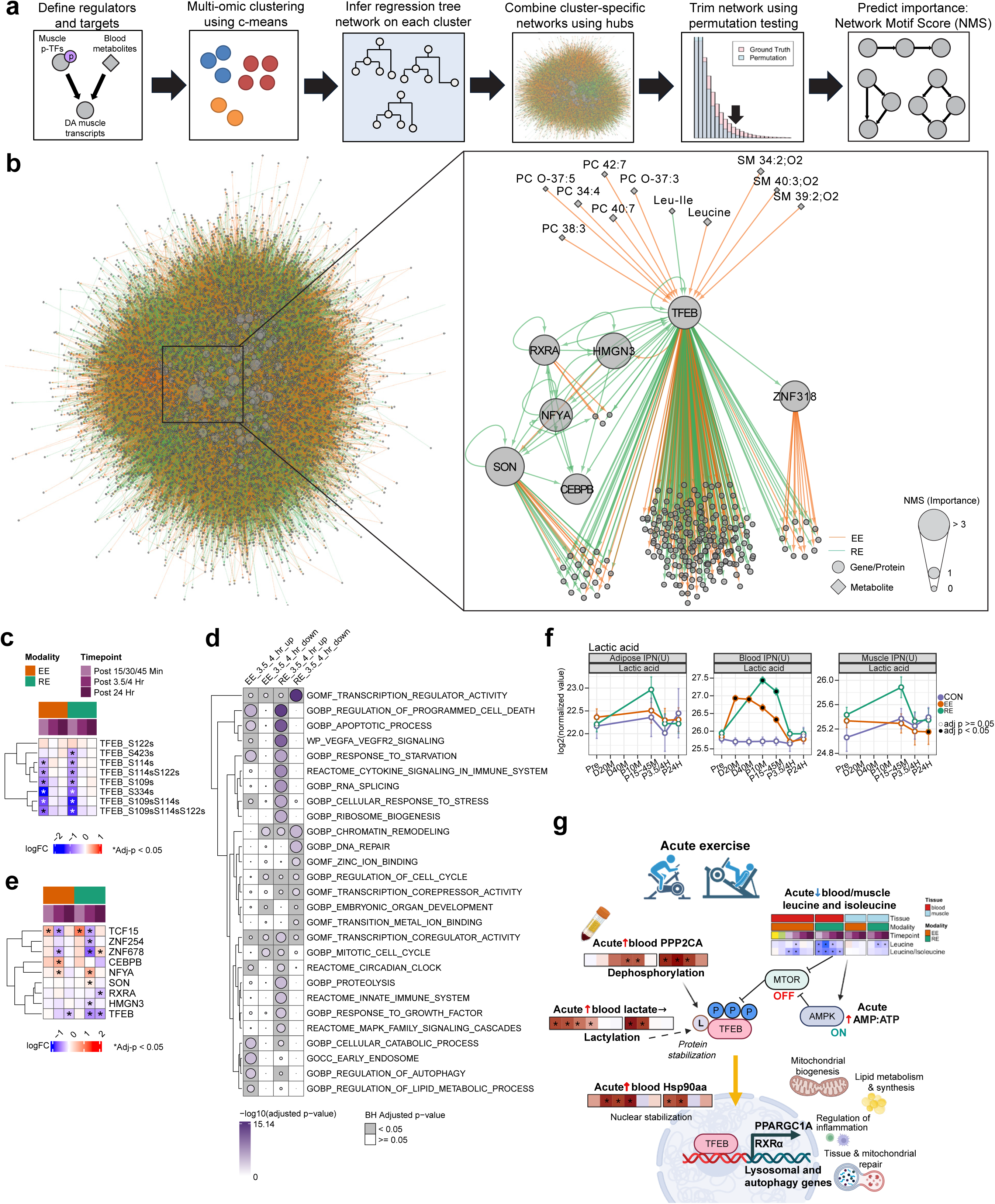
Multi-omic regulatory network analysis. **a**, SC-ION workflow. **b**, (left) Network merged across exercise modalities. (right) First-neighbor network of *TFEB*, filtered for upstream metabolites, ChIP-seq validated TFEB targets, and shared targets of TFEB and RXRA (not necessarily ChIP-seq validated). All metabolites and high importance targets are labeled. Size represents importance (NMS). **c**, Heatmap of TFEB phosphosites log fold change in SKM. **d**, ORA on TFEB-bound target genes in EE or RE using the C5 database. Genes were separated by fold change (up: positive, down: negative) prior to ORA. Dot plot shows selected significant terms with adjusted p-value < 0.05 in any group. **e**, Heatmap of RNA-seq log2 fold-change of the shared targets of *TFEB* and *RXRA*. **f**, Temporal expression of lactate (quantified as lactic acid) in response to exercise across all three tissues. Data are shown as group-timepoint mean ± 95% confidence interval with a black circle indicating significance relative to control at FDR<0.05. **g**, Schematic illustrating the predicted multi-omics, multi-tissue TFEB signaling module in response to exercise.

In particular, we focused on TFEB (Fig. 6b), which ranked highly in network importance and is a key regulator of lysosomal and autophagy genes, with recent murine data identifying roles for TFEB in mitochondrial biogenesis and metabolic flexibility.^80,81^ TFEB has been shown to be exercise-responsive in rodents and humans.^17,82,83^ Notably, TFEB is in the same TF family as MITF, identified by HOMER as containing an enriched binding motif among SKM DEGs (Fig. 5h). We filtered SC-ION predicted TFEB targets to those showing prior evidence of TFEB binding via Chromatin Immunoprecipitation (ChIP) (Fig. 6b; Supplementary Table 6); SC-ION predicted TFEB targets were DA at various timepoints: a subset showed strong upregulation either early post-exercise or at 24 hours post-exercise, while most decreased at 3.5 hours post-exercise, with a stronger response in RE compared to EE (Extended Data Fig. 8a). One of these targets was TFEB itself, which decreased significantly by 24 hours in SKM following both RE and EE (Extended Data Fig. 8b).

Motivated by the network centrality and functional plausibility of TFEB activation with acute exercise, we further explored functional regulation of TFEB. Shifts in TFEB phosphorylation, specifically a modality-consistent dephosphorylation at S109, S114, and S122 at 15 minutes post-exercise (Fig. 6c), were used in SC-ION to predict its putative target genes. Further, it is known that TFEB S122 is an mTOR phosphorylation site which restricts TFEB to the cytosol,^84^ while this site is commonly dephosphorylated by protein phosphatase 2A (PP2A).^85^ In response to acute exercise, we observed an upregulation of multiple genes encoding PP2A subunits in the blood and SKM following exercise (Extended Data Fig. 8c), suggesting that exercise-induced PP2A may dephosphorylate TFEB and promote its nuclear translocation and transcriptional activity.^86^ Further, we observed that HSP90AA1, an upstream regulator of TFEB nuclear stabilization,^87^ displayed acute transcript induction in response to both EE and RE in the blood and muscle (Extended Data Fig. 8d).

To build a more comprehensive picture of downstream TFEB activity, we performed an ORA on all ChIP-bound targets of TFEB (not just those predicted by SC-ION) that increased or decreased 3.5 hours post-exercise (Fig. 6d). TFEB-regulated transcriptional targets that were upregulated by exercise were related to the apoptotic process, cellular responses to starvation and stress, VEGF signaling, and clock regulation, consistent with known roles of TFEB.^88,89^ Additional TFEB-linked pathways induced at 3.5 hours included upregulation of autophagy following EE and RE (Extended Data Fig. 8e), further suggesting a SKM remodeling role for TFEB in the acute exercise response. Further evaluation of other highly-ranked SC-ION-predicted targets included RXRA, which regulates lipid metabolism and is linked to the TFEB regulatory network.^90,91^ Combined with network identification of NFYA, a clock regulatory TF, and CEBPB, which regulates circadian autophagy,^92^ this supports a role for TFEB in orchestrating circadian variation of reparative processes in the acute exercise response (Fig. 6e).^93^

We next identified putative metabolite regulators of TFEB. The branched chain amino acids leucine and isoleucine decreased acutely in plasma with EE and RE (Extended Data Fig. 8f), consistent with attenuated downstream mTOR signaling,^94^ and permissive TFEB activation.^95^ Lactate increased acutely with exercise (Fig. 6f), and lysine lactylation has recently been shown to promote TFEB stabilization, nuclear retention, and transcriptional activity.^96^ Together, these findings support a model where exercise-induced reductions in mTOR activation and increases in PP2A activity dephosphorylate TFEB, while acute metabolite changes promote TFEB stabilization, collectively facilitating nuclear localization and a targeted downstream transcriptional network (Fig. 6g).

### Cellular communication network factor 1 is a novel candidate exerkine

To identify putative exercise-induced secreted factors that may, in turn, lead to salutary health effects (i.e., “exerkines”), we compared DA transcriptomic, proteomic, and phosphoproteomic features from SKM and AT, as well as blood transcriptomic data, to plasma proteomic data in order to identify matched molecular changes in tissues and plasma. This analysis revealed 115 molecular features gene-matched to DA plasma proteins. These 115 features were primarily transcripts in blood and SKM and demonstrated directionally concordant changes with plasma proteins following both EE and RE (Fig. 7a, Extended Data Fig. 9a,b; Supplementary Table 7). Of these, 67% (77) were directionally concordant (i.e. both up or both down), and 28% (n=32) demonstrated tissue changes concurrently or prior to plasma changes.

**Figure 7-.**
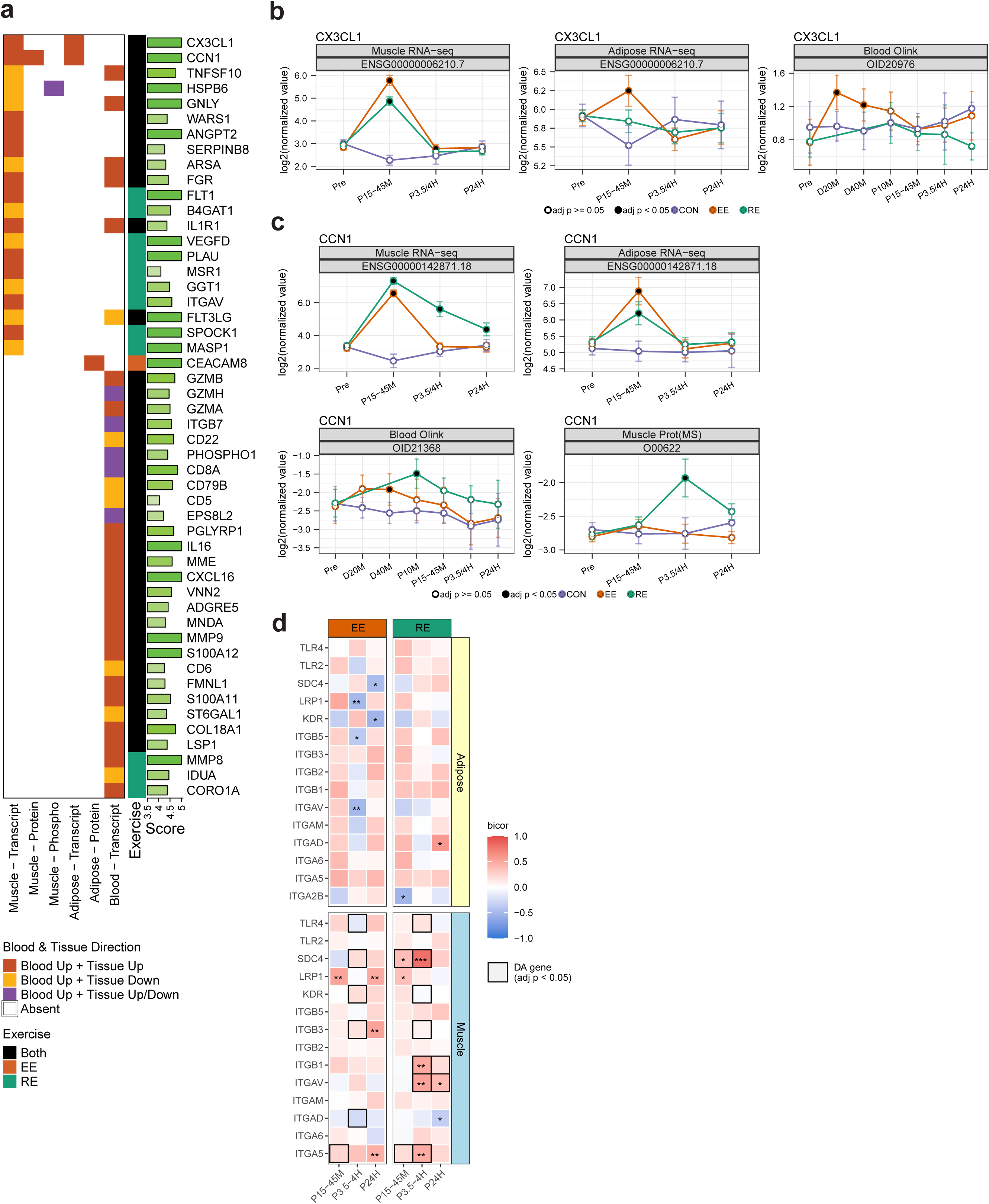
Cellular communication network factor 1 is a novel candidate exerkine. **a**, Heatmap showing differentially regulated features across SKM, AT, and blood transcriptomics, proteomics, and phosphoproteomics (adjusted p < 0.05 at any time points) that also exhibit significant upregulation in plasma proteomics (adjusted p < 0.05 at any timepoint, including during and post-exercise). Gene symbols from plasma Olink proteomics were used to map to corresponding tissue-specific features. “Blood up” indicates significant upregulation of a given plasma protein at one or more timepoints; “Tissue up” indicates significant upregulation of tissue-specific features at one or more timepoints. Horizontal green bars denote the extracellular localization score (range 0–5) using COMPARTMENTS (see Methods); only features with a score > 4 are shown. The exercise column indicates features differentially regulated by EE, RE or “Both”. **b,c**, Temporal trajectories of Fractalkine (*CX3CL1*, B) and cellular communication network factor 1 (*CCN1*, C) transcript expression in AT and SKM, and plasma protein abundance, and SKM protein (CCN1 only). EE = Endurance Exercise, RE = Resistance Exercise, CON = control group, Pre = pre-exercise, D20M = during 20 minutes, D40M = during 40 minutes, P10M = 10 minutes post-exercise, P15-45M = 15, 30, or 45 minutes post-exercise (depending on tissue), P3.5-4H = 3.5 or 4 hours post-exercise (depending on tissue), P24H = 24 hours post-exercise. Data is shown as group-timepoint mean ± 95% confidence interval for the group mean with a black circle indicating model significance relative to control at adj. p-val<0.05. Plots indicate Ensembl ID for transcripts and Olink ID or Uniprot accession number for proteins. **d**, Heatmap showing intra-tissue correlation (biweight midcorrelation, bicor value) between *CCN1* transcript expression and known CCN1 receptor expression. Box outlines indicate timepoint-tissue-modality combinations with DA expression of the given receptor (adjusted p < 0.05). Heatmap color indicates biweight midcorrelation with asterisks indicating nominal significance of correlation: * < 0.05, ** < 0.01, *** <0.001.

Several of these putative protein exerkines are related to angiogenesis, including PXN, NOS3, and ANGPT2 (Fig. 7a, Extended Data Fig. 9b),^97–99^ and inflammatory responses such as TNF superfamily proteins.^100^ We focused on features most likely to be localized to the extracellular compartment given their likelihood to be secreted (Fig. 7a), resulting in 50 primary candidates.^101^ All other candidates are shown in Extended Data Fig. 9b and Supplementary Table 7.

While most genes/proteins showed changes in just one tissue-ome combination, several demonstrated multi-tissue and multi-omic changes (Fig. 7a). For example, fractalkine (*CX3CL1*), a previously described exerkine,^102^ demonstrated increased expression in SKM (EE and RE) and AT (EE only, Fig. 7a,b). Expression of cellular communication network factor 1 (CCN1), an angiogenic factor previously reported to be secreted by AT and SKM,^103–105^ demonstrated pronounced exercise effects across tissues. It increased with both modalities in the AT and SKM, while plasma protein levels increased during EE and immediately following RE (Fig. 7c). Expression levels of *CCN1* in AT and SKM were not correlated (Extended Data Fig. 9c) at any timepoint, suggesting tissue-specific expression.

To explore CCN1 signaling further, we identified a list of known receptors for CCN1,^106–110^ and each was assessed at the transcript level for responsiveness to acute exercise and correlation at each timepoint with CCN1 expression levels in the same tissue (Fig. 7d). While transcripts in AT did not significantly change with exercise or consistently correlate with CCN1 levels, in SKM multiple receptors met both criteria. Most notable was *SDC4*, which encodes the coreceptor syndecan-4. In RE, *SDC4* levels both responded to acute exercise and were strongly correlated with *CCN1* transcript levels, particularly notable considering this correlation was observed at 3.5 hours post-RE – the only modality-tissue-timepoint with significant increases in CCN1 at the protein level. Taken together, these data suggest CCN1 is a novel exerkine candidate, with CCN1 signaling positioned as an important exercise response pathway, particularly in RE, given the concordant upregulation of known CCN1 receptors.

## Discussion

The Molecular Transducers of Physical Activity Consortium is simultaneously focused on building a molecular map of exercise response while also using that map to answer pivotal questions about molecular responses to exercise. While the present work represents a subset of the total data anticipated in MoTrPAC (pre-Covid suspension, acute exercise bout only),^111^ we: (1) identify thousands of molecular features that change with acute bouts of EE or RE, (2) detail novel patterns of human exercise responses including unique and sometimes divergent patterns of response to EE or RE, and (3) make available–both through R packages and an online browser–the full resulting dataset. This represents the largest multi-tissue, multi-omic, randomized controlled trial of acute exercise available. Further, we highlight the crucial role of the non-exercising group in our data. While many smaller multi-omic studies utilize pre-exercise samples as the only comparator, many molecular features shift over time due to fasting, circadian rhythms, and stress response from sampling; our data identify and discern these effects from true exercise effects.

By collecting multiple tissue specimens (blood, SKM, and AT) from MoTrPAC participants, we were able to compare and contrast tissue-based molecular signatures of exercise. We observed that tissue responses to acute exercise, for either EE or RE, were more different than they were similar. Even when restricted to transcriptomics and metabolomics, which had balanced sample sizes across tissues, just 50 features among thousands were affected by exercise in all three tissues. Further, when temporal patterns of molecular responses were analyzed using pathway enrichment, several pathways were engaged in all three tissues, but the genes that were DA within these pathways depended much more on tissue type than on exercise modality. Thus, tissues utilize different genes within key pathways to respond to exercise on varied timescales.

Still, some pathways were affected in multiple tissues and in both modalities, including mitochondrial pathways, VEGF and other angiogenic signaling pathways, and tissue regenerative and remodeling pathways. Given the known elements of the biological response to exercise, these pathways are expected to be engaged throughout the body. What emerged in the present data were details of the specific features and temporal responses in each tissue: for example, elements of the VEGF signaling pathway were engaged at early and intermediate timepoints post-exercise in AT and blood, respectively, and across the entire post-exercise period in SKM.

The present data provide a valuable opportunity to compare and contrast EE and RE. While there were many shared molecular responses between EE and RE, there were also a large number of features, particularly in the blood and SKM, that demonstrated modality-specific responses. A small but important subset of these modality-specific features were actually divergent in their response, with opposing effects between modalities. These are likely to be fundamental to the functional responses in each modality. As an example, *LRRC14B* not only diverged between modalities, but its association with exercise is novel. Indeed, very little was previously known about this gene at all,^46^ and the present work positions it as a potential mechanistic switch in EE versus RE.

Contrasting EE and RE also provided a natural window into the connection between molecular response and cardiorespiratory fitness. For the subgroup of SKM transcripts associated with VO_2_peak at the pre-exercise baseline, those with acute bout responses that were EE-specific and upregulated at 24 hours post-exercise had a positive association with VO_2_peak at baseline. RE-specific transcripts upregulated at 24 hours had just the opposite relationship–a negative association with VO_2_peak at baseline. These findings raise important questions and direct future investigations. First, concordant results (those which are upregulated by acute exercise and associated with higher VO_2_peak at baseline) are prime candidates as transducers of exercise-induced cardiorespiratory fitness adaptation, hypotheses readily testable in forthcoming MoTrPAC analyses involving both highly active participants and exercise training of sedentary participants.^111^ Further, the discordant relationship between RE-specific transcripts and baseline VO_2_peak confounds evidence that RE can lead to improvements in VO_2_peak,^112^ but likely reflect the acute effects of RE, including induction of SKM transcriptional pathways related to inflammation and sarcomere remodeling.^113^ Whether these transient changes are sustained during training and whether these transcripts play important roles in transducing strength adaptation, relative to aerobic adaptation, can also be explored in forthcoming data.

Emerging across multiple facets of the data, the importance of autophagy and protein turnover was highlighted, an effect more prominent in RE. While TFEB’s role in autophagy is well described,^84,114–116^ our data extend this observation by highlighting its central role in activating autophagy and linked circadian and metabolic transcriptional networks in the acute exercise response. Probing of additional phosphoproteomic datasets in rodents and humans reveal acute dephosphorylation of TFEB as a conserved acute exercise response.^17,82^ TFEB is a master regulator of autophagy,^117^ and autophagy has been posited as a key element of the benefits conferred by exercise, including cognitive and skeletal muscle benefits.^2,118,119^ We additionally identified putative metabolic regulators of TFEB dephosphorylation to include an acute decrease in plasma leucine and isoleucine and propose a model where acute increases in plasma lactate stabilize TFEB transcriptional activity.^96^ Using the present TFEB data as a map for future studies will help to mechanistically link exercise-induced TFEB activation with SKM and non-SKM organ health.

Understanding the communicative link between exercising SKM and other tissues is a key area of interest. Here, we posit many plausible new exerkines, though *CCN1* provides a particularly striking example. We show here for the first time that CCN1 levels at the transcript and/or protein level shift in AT, SKM, and plasma. Pathways downstream of CCN1 signaling, such as ECM remodeling and angiogenesis, which are themselves known to be critical to SKM and exercise adaptation,^120,121^ may thus extend to other tissues via CCN1 signaling. Whether CCN1 is signaling in an autocrine, paracrine, or endocrine fashion is yet unclear; forthcoming data from acute exercise in rats are likely to be informative.^111^ Interestingly, CCN1 response is described in cardiac muscle and has been shown for myocardial infarction, both in scar formation^122^ and angiogenesis.^123^ Further it has been described as a biomarker of adverse events in heart failure,^124^ presumably due to induction in response to injury. Given the present findings, it will be instructive to determine the effects of exercise on CCN1 signaling in cardiovascular disease.

Our data have several important limitations. Across tissues, timepoints, modalities and omes, the number of samples is lower than the full cohort of N = 175 due to limitations in biopsy numbers per participant, omic downsampling due to feasibility, assay limitations, and trial design. Nonetheless, the paired nature of sampling (pre- and post-exercise) and inclusion of time-matched controls fortifies resolution of independent exercise effects. Further, while this sample size was among the largest in the field of acute exercise, it was still too limited to assess exercise interactions with sex, age, or other baseline characteristics reliably. Nonetheless, SKM transcriptomic responses to EE and RE across multiple timepoints aligned with findings from a smaller study of 24 male subjects,^125^ as detailed in a SKM-specific companion paper.^34^ Finally, while control samples are a highlight of the study, we did observe greater technical variation in ATAC-seq control samples, potentially limiting the number of DA peaks, though many more peaks were identified in a within-group analysis (Supplementary Results).

In sum, we provide the results of the largest multi-tissue, multi-omic trial of acute exercise, identifying a wide range of new insights to drive forward the field of exercise physiology at the molecular level. The results of the full randomized controlled trial (N = 1,541), the pediatric cohort (N = 457), and the highly active study (N = 295) are forthcoming.^37,111^ Given the large scope of this study, a number of companion papers explore the molecular responses in greater detail by tissue and ome.^32–35^ Moreover, we anticipate the full MoTrPAC cohort will validate and expand the current findings to determine the roles of sex, age, genetics, and other demographic variables on acute exercise responses and on training adaptations.

## Supporting information

Supplementary Results

## Extended Data Figure Legends

**Extended Data Figure 1.**
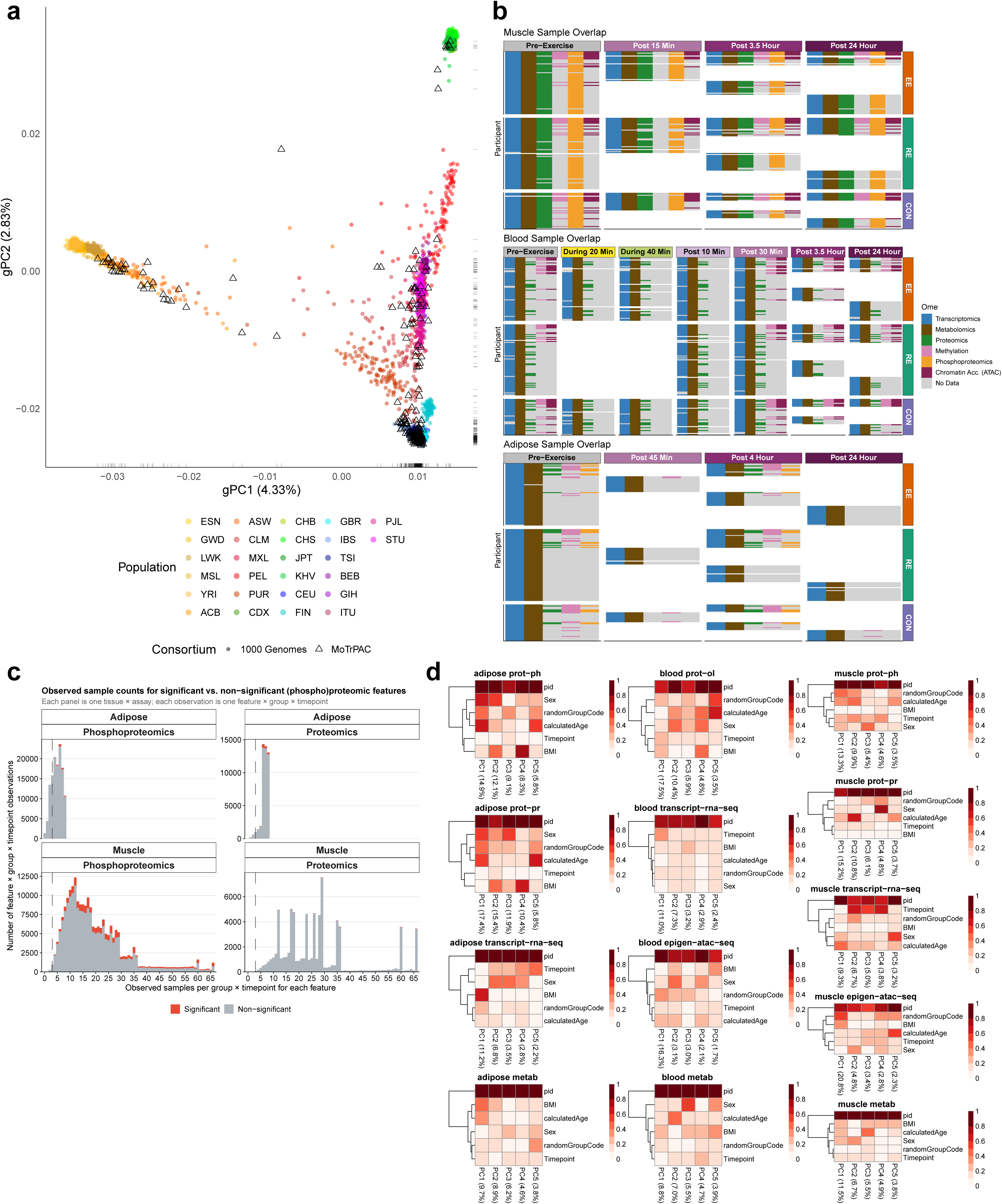
Participant genotypes, sample-ome overlap distribution and tissue-ome covariate structure, related to Figure 1. **a**, First two principal components of genetic ancestry. Participants from MoTrPAC are plotted using triangles, while participants from 1000 Genomes are plotted using points. Categorical ancestry according to 1000 Genomes is indicated by color. MoTrPAC participants are not categorized by ancestry. **b**, Matrix of omic data availability. Each row of the heatmap represents a participant, with each participant represented once in each of the three tissues. If data is available for a particular participant for a given tissue-ome-time point combination, this is indicated by a colored cell in the matrix. A grey cell indicates no omic data available for the sample collected. A white cell indicates that no sample was collected in that participant for that tissue-time point. **c**,Number of samples observed in each exercise group-timepoint combination in mass-spectrometry proteomic and phosphoproteomic platforms. Since there is missingness inherent in the measurement of mass-spectrometry, the number of samples observed for the same group-timepoint combination can vary by feature. Any feature that was deemed statistically significant in the differential analysis is highlighted in red. A line at n=3 indicates that the cutoff used for input to differential analysis; any feature with any group-timepoint with less than n=3 was removed from downstream analysis. **d**,Canonical correlation analysis of the first five principal components of ome-tissue combinations to key MoTrPAC experimental covariates. Colors in each block represent canonical correlation between each principal component and each experimental parameter. Metabolite platforms were combined into one matrix and subset to participant-timepoint combinations present in all metabolomic platforms. Principal component labels indicate percentage of total omic variance explained by each principal component.

**Extended Data Figure 2.**
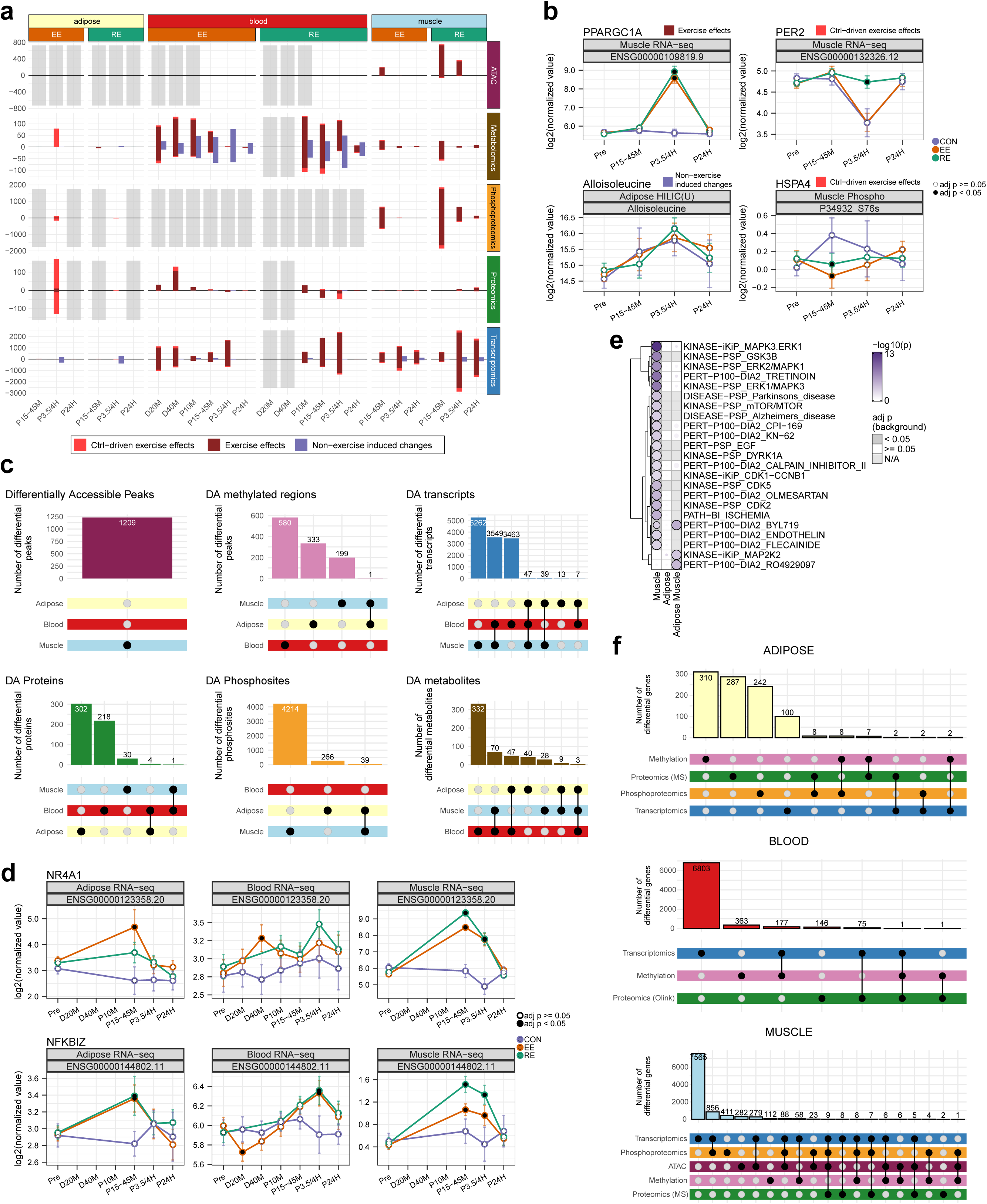
Distribution and overlap of differential multi-omic features across tissues, related to Figure 2. **a**,Barplot summarizing the classification of molecular features by type of differential response across tissues and omes. Features were grouped into three categories based on significance patterns in exercise versus control comparisons: (1) Control-driven exercise effects; (2) Exercise effects; and (3) Non-exercise induced changes. This categorization is intended to distinguish biologically meaningful exercise responses from potential false positives from otherwise uncontrolled experimental designs. Each feature upregulated relative to control is plotted above the axis, and each downregulated relative to control is plotted below the axis. **b**,Representative examples of molecular features classified into categories based on their differential response patterns across timepoints and conditions. *PPARGC1A* (Muscle RNA-seq) represents a robust exercise effect. HSPA4 (Muscle Phosphoproteomics) and *PER2* (Muscle RNA-seq) exemplifies a control-driven effect. Alloisoleucine (Adipose metabolomics) illustrates non-exercise induced changes. Pre = pre-exercise, P15-45M = 15, 30, 45 minutes post-exercise (depending on tissue), P3.5-4H = 3.5 or 4 hours post-exercise (depending on tissue, P24H = 24 hours post-exercise. Data is shown as group-time point mean ± approximated 95% confidence interval with a black circle indicating significance relative to control at adj. p-val<0.05. Plots indicate Ensembl ID for RNA or Uniprot IDs for proteins where appropriate. **c**,UpSet plots by ome across tissues: ATAC-seq, MethylCap-seq, transcriptomics, proteomics, phosphoproteomics, and metabolomics. Each plot displays the overlap of significant features (adjusted p-value < 0.05) across three tissues regardless of exercise modality or timepoint. Vertical bars represent the number of features unique to or shared among the tissues, as shown in the matrix below each plot. Numbers on top of the bars indicate the total number of features in that set. **d**,Temporal trajectories of select DA features shared between all three tissues. Pre = pre-exercise, D20M = during 20 minutes, D40M = during 40 minutes, P10M = 10 minutes post-exercise, P15-45M = 15, 30, 45 minutes post-exercise (depending on tissue), P3.5-4H = 3.5 or 4 hours post-exercise (depending on tissue, P24H = 24 hours post-exercise. Points represent group means ± approximated 95% confidence interval; black points circles denote comparisons to control with adjusted p-values < 0.05. Plots indicate Ensembl ID for RNA. **e**,Over-representation analysis of DA phosphosites unique or shared between the AT and SKM against the PTMsigDB collection. **f**,Panel of three UpSet plots, one for each tissue (AT, SKM, and blood), illustrating the intersection of all significant features (adjusted p-value < 0.05) across epigenome (ATAC-seq, MethyCap-seq), transcriptome, proteome, and phosphoproteome (metabolomics data was not included). Features were mapped to gene symbols to enable direct comparison across omics layers. For each tissue, vertical bars represent the number of features unique to, or shared among, the indicated omes as shown by the matrix below each plot. Features that were DA at any time point and in either modality were considered for matching across omic layers.

**Extended Data Figure 3.**
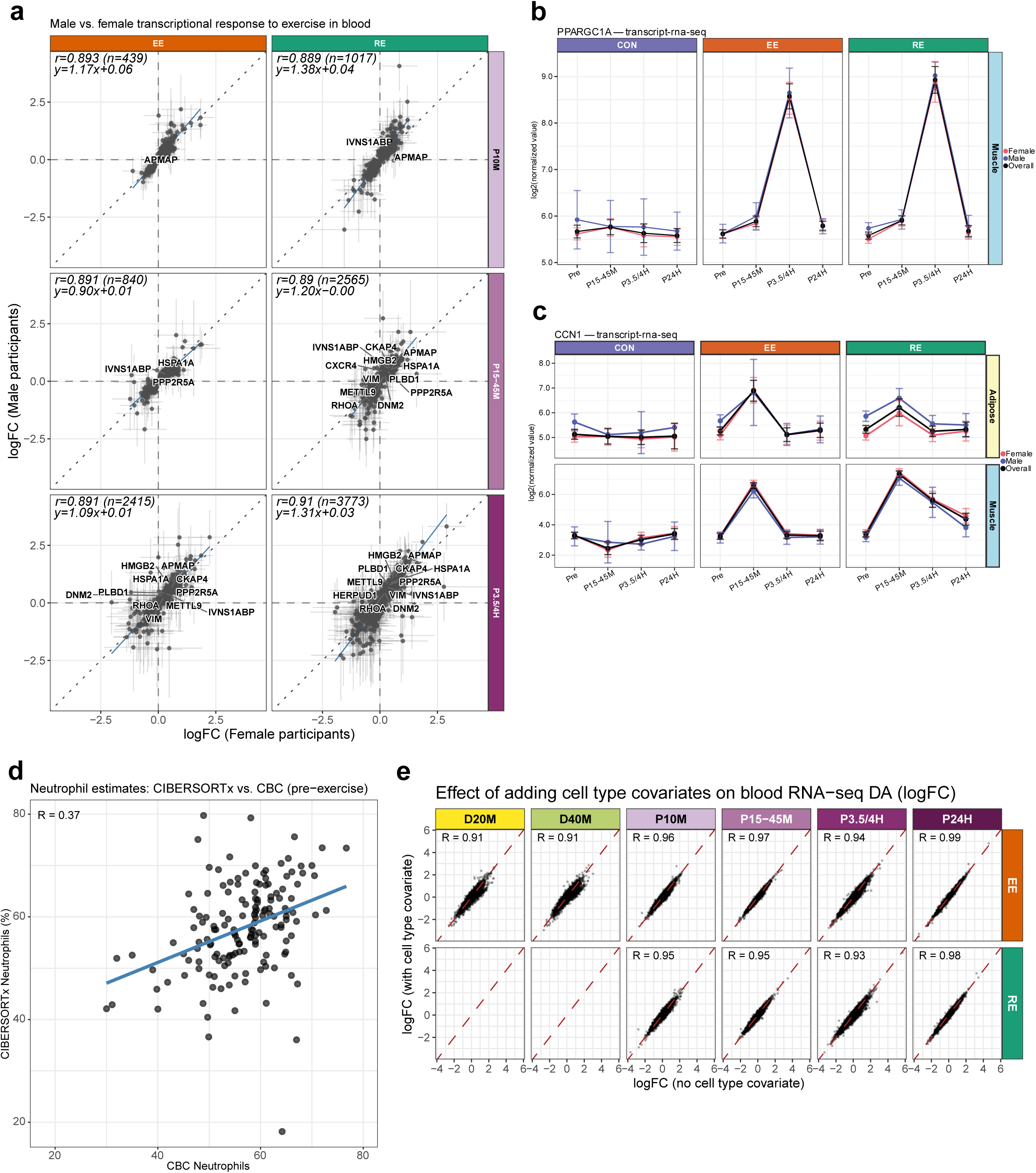
Sex and cell type-specific sensitivity analysis from blood transcriptomics data, related to Figure 2. **a**, Exercise-induced log fold-changes (logFC, post-exercise relative to pre-exercise) were estimated separately in male and female participants and are plotted against one another for blood transcripts: female logFC on the x-axis and male logFC on the y-axis. Panels are arranged by post-exercise timepoint (rows: P10M, 10 minutes post; P15-45M, 15/30/45 minutes post; P3.5/4H, 3.5/4 h post) and exercise modality (columns: EE, endurance exercise; RE, resistance exercise). Each point is a transcript that was significant in the primary, sex-combined PreCAWG differential analysis (FDR < 0.05); horizontal and vertical bars show the 95% confidence intervals of the female and male logFC estimates, respectively. The dotted diagonal is the line of identity (equal response in both sexes) and the blue line is the ordinary least-squares fit; the Pearson correlation (r), the number of features (n), and the regression equation are annotated in each panel. Labeled transcripts are up to 10 per timepoint with the smallest male-versus-female adjusted p-value (FDR < 0.05) in RE; the same transcripts are labeled in both columns. Dashed grey lines mark zero logFC. **b,c**, For PPARGC1A (B) and CCN1(C), the mean normalized abundance (± 95% confidence interval) is shown at each timepoint for female (red), male (blue), and all (overall, black) participants, faceted by tissue–assay (rows) and exercise group (columns: CON, control; EE, endurance; RE, resistance). Timepoints: Pre, pre-exercise; D20M/D40M, 20/40 minutes during exercise; P10M, 10 minutes post; P15-45M, 15/30/45 minutes post; P3.5/4H, 3.5/4 h post; P24H, 24 h post. Concordant male and female trajectories indicate that the reported response for these features is not driven by the female-majority composition of the cohort. Per-sex sample sizes are reduced relative to the full-cohort analysis, so confidence intervals are wider and these panels should be interpreted as descriptive. Points represent normalized abundance and no indication of statistical significance is present in this panel. **d**, Concordance between CIBERSORTx-estimated and clinically measured neutrophil proportions. Each point is a participant, comparing the clinical complete blood count (CBC) neutrophil value (x-axis) at the screening visit with the CIBERSORTx-deconvolved neutrophil percentage (y-axis) from the pre-exercise timepoint. The blue line is the ordinary least-squares fit and the Pearson correlation (r) is annotated. Two samples with <5% measured neutrophils from CBC, assumed to be erroneous, were removed from this analysis. **e**, Effect of adding deconvolution-derived cell-type covariates on the blood RNA-seq differential analysis. Per-transcript exercise effect estimates (logFC) from the main model (x-axis) are plotted against estimates from the model that additionally includes the z-scored coarse cell-type fractions (T cells, B cells, NK cells, monocytes, and neutrophils) as numeric covariates (y-axis). Each point is a transcript; panels are faceted by exercise group (rows: ADUEndur, endurance; ADUResist, resistance) and post-exercise timepoint (columns). The dashed red line is the identity line and the Pearson correlation (R) is annotated in each panel.

**Extended Data Figure 4.**
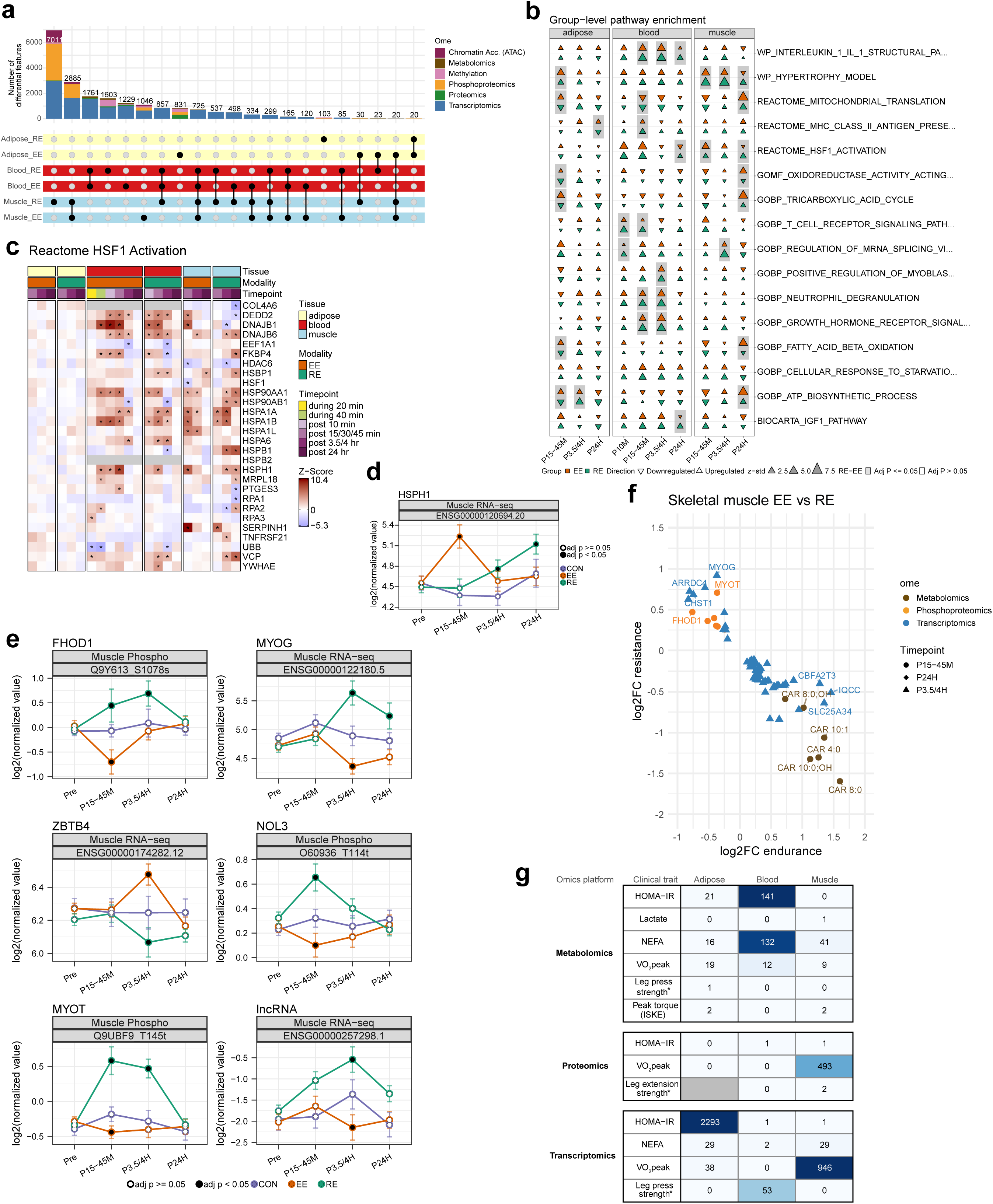
Differential responses to endurance and resistance exercise, related to Figure 3. **a**, UpSet plot depicting the overlap of significant molecular features across tissue-modality combinations, at any timepoint. Vertical bars indicate the number of features that are unique or shared, while colors represent the omic contribution. Numbers on top of the bars indicate the total number of features in that set. Black dots in the lower matrix indicate the tissue-modality membership for each intersection set. **b**, CAMERA-PR (Correlation Adjusted MEan RAnk gene set test—PreRanked) pathway enrichment heatmaps highlighting pathways that are different between EE and RE. The CAMERA-PR results for EE vs. CON and RE vs. CON are described by orange and green triangles, respectively. The direction of the triangle indicates upregulation in EE or RE relative to CON. Any pathways that are significant when directly compared between EE and RE are highlighted with a light grey background. The top 5 pathways by lowest p-value in each tissue are visualized. **c**, Heatmap of Z-scores for transcriptomic features belonging to the HSF1 pathway. Columns represent timepoints grouped by tissue and exercise modality. **d**, Trajectories for *HSPH1* transcript. Pre = pre-exercise, P15-45M = 15, 30, 45 minutes post-exercise (depending on tissue), P3.5-4H = 3.5 or 4 hours post-exercise (depending on tissue, P24H = 24 hours post-exercise. Data is shown as group-time point mean ± 95% confidence interval with a black circle indicating significance relative to control at adj. p-value<0.05. Plots indicate Ensembl ID. **e**, Trajectories of selected features in SKM included in figures 3c and ED4f. Pre = pre-exercise, P15-45M = 15, 30, 45 minutes post-exercise (depending on tissue), P3.5-4H = 3.5 or 4 hours post-exercise (depending on tissue, P24H = 24 hours post-exercise. Data is shown as group-time point mean ± 95% confidence interval with a black circle indicating significance relative to control at adj. p-value<0.05. Plots indicate Ensembl for RNA or Uniprot IDs for proteins where appropriate. NOL3 = Nucleolar Protein 3, MYOT = myotilin. **f**, Oppositely regulated features in the two exercise modalities (relative to CON, adj. p-value<0.05), that were also significant in a direct comparison between EE and RE (adj. p-value<0.05) in SKM. Point color indicates assay and symbol indicates time point. **g**, Heatmap of significant omics features associated with clinical traits across tissues. Counts of significant (FDR < 0.05) features are shown by assay, clinical trait, and tissue type. Clinical traits include: peak blood lactate,HOMA-IR, leg press strength, non-esterified fatty acids (NEFA), peak torque (isometric knee extension), and VO_2_peak. Grey cells indicate the combination did not have sufficient samples to estimate associations in any features (methods). Strength metrics (leg press strength) were measured in only resistance exercise participants, representing 56-73 samples per model, compared to up to 172 for non-strength metrics.

**Extended Data Figure 5.**
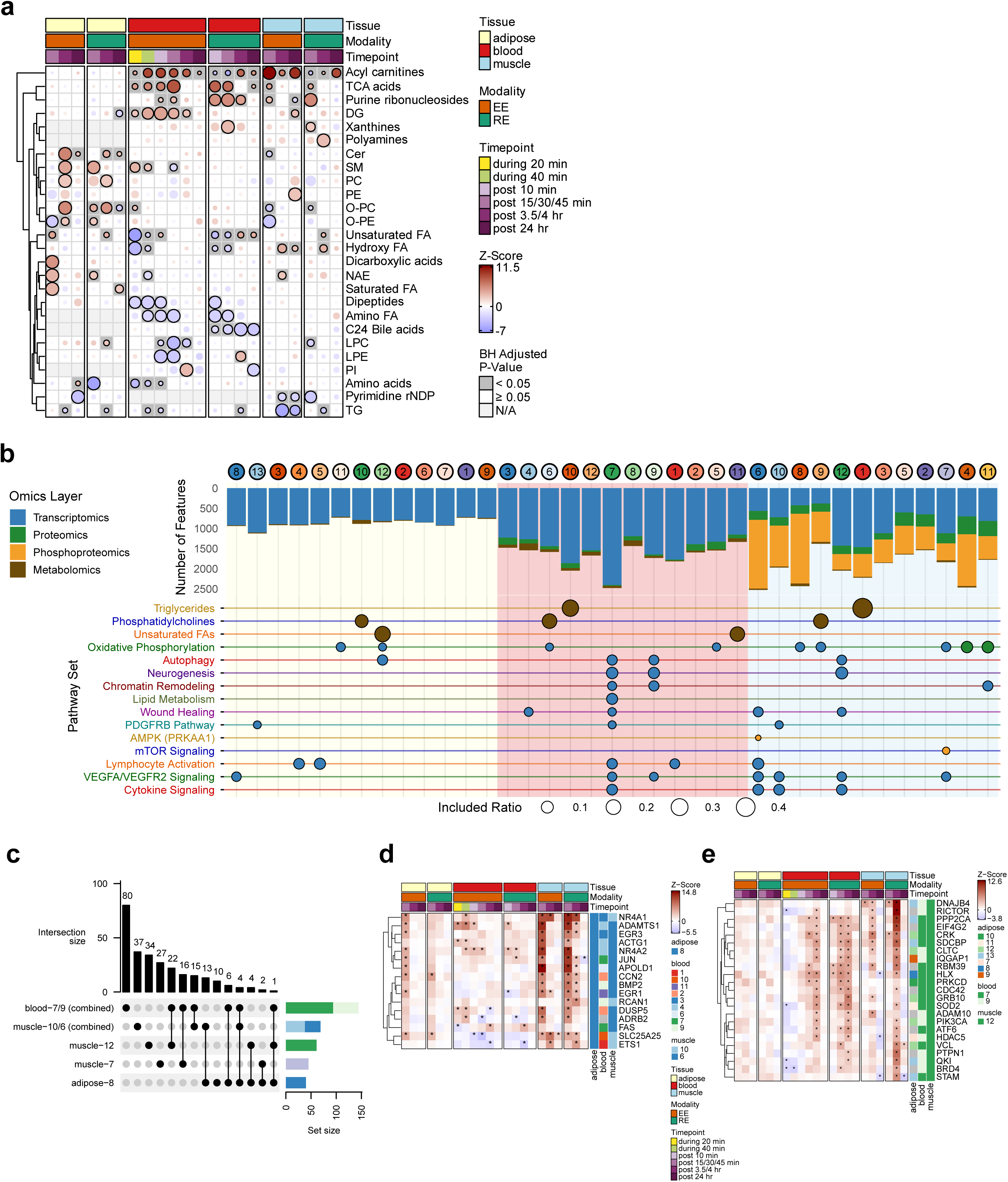
Multi-tissue metabolomic enrichments, C-means trajectories and associated pathways and multi-tissue feature heatmaps, related to Figure 4. **a**, Pathway Enrichment for each tissue-timepoint in metabolomic RefMet classes via Pre-ranked Correlation-Adjusted MEan-RAnk (CAMERA-PR). **b**, Temporal cluster enrichments, extended. Circles represent cluster numbers and fill colors corresponding to the temporal trajectories shown in Fig. 4c-d. Clusters are organized by tissue and then by timepoint of maximal response (maximal absolute deviation [MAD] from zero).The bar chart shows the total number of features assigned to each cluster (with a membership probability > 0.3), with bar segments colored by their omics layer of origin. Background shading indicates the tissue of origin (adipose, blood, or muscle). The dot plot displays significant pathway enrichments (FDR < 0.05) for each cluster. The size of each point is proportional to the “included ratio” (the proportion of features within the cluster that belong to the pathway set). The fill color of each point indicates the omics layer of the enriched features. AMPK (PRKAA1) = PSP_PRKAA1, mTOR Signaling = PSP_MTOR, Lipid Metabolism = GOBP_LIPID_METABOLIC_PROCESS, PDGFRB Pathway = PID_PDGFRB_PATHWAY, Chromatin Remodeling = GOBP_CHROMATIN_REMODELING, Triglycerides = REFMET_TG, Unsaturated FAs = REFMET_Unsaturated FA, Neurogenesis = GOBP_NEUROGENESIS, Phosphatidylcholines = REFMET_PC, Oxidative Phosphorylation = MITOCARTA_OXPHOS, Wound Healing = GOBP_WOUND_HEALING, Lymphocyte Activation = GOBP_LYMPHOCYTE_ACTIVATION, VEGFA/VEGFR2 Signaling = WP_VEGFA_VEGFR2_SIGNALING, Cytokine Signaling = REACTOME_CYTOKINE_SIGNALING_IN_IMMUNE_SYSTEM, **c**, UpSet plot comparing feature membership in each cluster shown in Fig. 4e. Clusters with a matching timepoint of maximal absolute deviation are paired for brevity. **d**,**e**, Multi-tissue heatmaps of Z-scores for transcriptomic features belonging to the VEGFA-VEGFR2 signaling pathway. Columns represent timepoints grouped by tissue and exercise modality. The right-hand annotation indicates the c-means cluster assignment for each gene in adipose, blood, and muscle, with colors corresponding to the cluster profiles in Fig. 4c-d and Extended Data Fig. 5b. Features assigned to adipose cluster 8 and muscle clusters 6 or 10 are shown in (d) and features assigned to blood clusters 7 and 9 and muscle cluster 12 in (e).

**Extended Data Figure 6.**
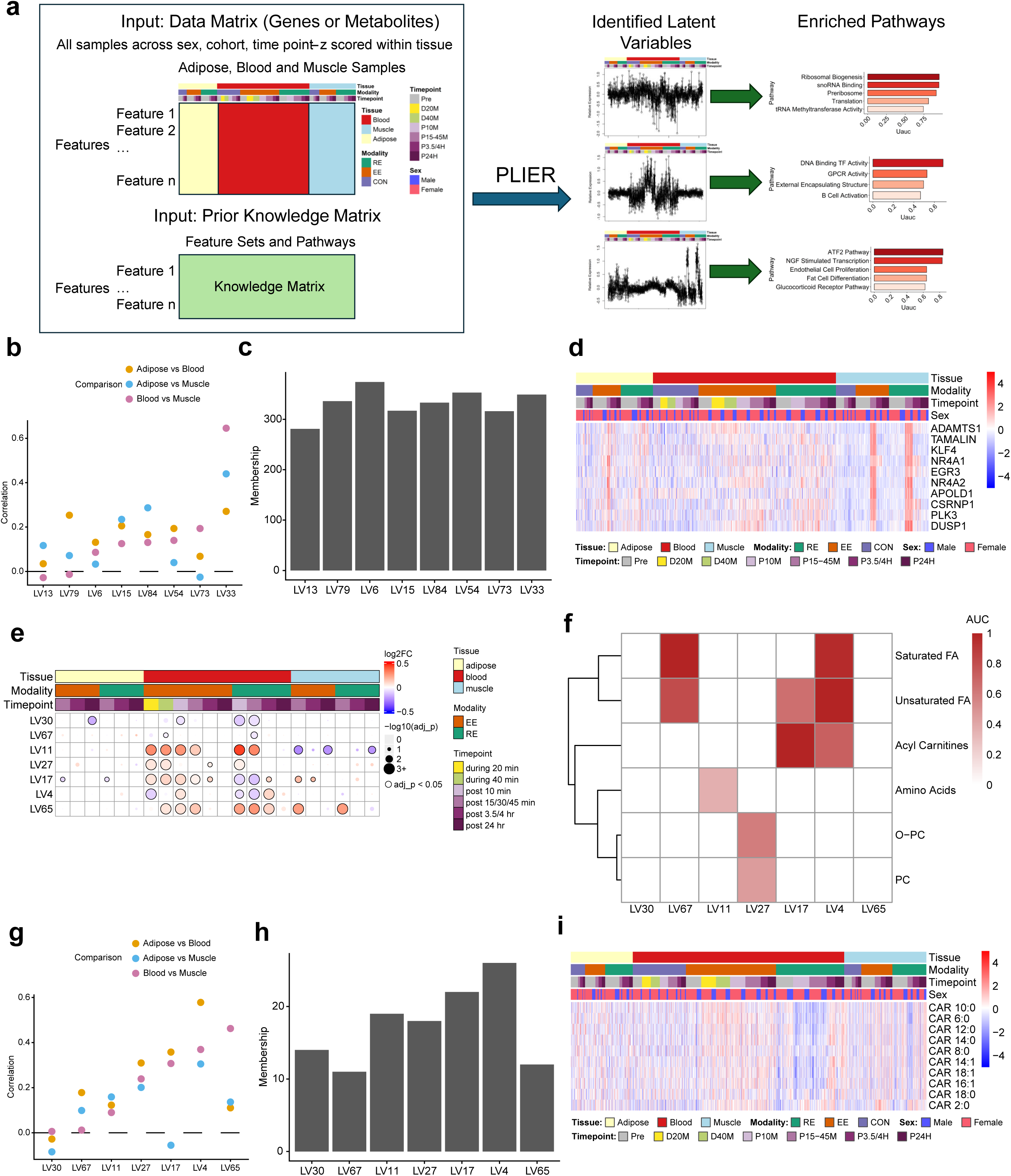
PLIER schematic, and expanded PLIER results, related to Figure 4. **a**, Schematic of PLIER method design as applied. PLIER used as input Z-scored normalized RNAseq data from all three tissues concatenated into a single input matrix where rows were the overlapping feature set and columns were the samples in each of the three tissues, as well as a matrix of prior knowledge for the inputted feature set. PLIER outputs a series of Latent Variables (LVs) reflecting distinct patterns of behavior across the measured samples with a set of associated features that align with the pattern and specific pathway enrichments based upon the prior knowledge and associated feature set. PLIER was run separately on the cross-tissue RNAseq and cross-tissue metabolomics data. **b**,**c**, The cross-tissue Pearson correlation (b) and count of highly associated transcripts (c) for each of the transcriptomic LVs described in Fig. 4g. **d**, Heatmap of the top ten genes associated with LV33 from the cross-tissue RNAseq PLIER analysis representing the within-tissue z-scores per gene for each participant at every tissue-time point combination. **e**, Bubble heatmap of the significance (size = adjusted p-value) and directionality (color = log2 fold-change) of the EE or RE response in each tissue described by a subset of metabolomic-data derived PLIER LVs with the most significant metabolomic exercise responses. **f**, Significantly enriched metabolite classifications for the metabolomics LVs described in (e). Scale is the AUC of a statistical test of association between classification and LV-associated genes. **g**,**h**, The cross-tissue Pearson correlation (g) and count of highly associated metabolites (h) for each of the metabolomics LVs described in (e). **i**, Heatmap of the top ten metabolites associated with LV17 from the cross-tissue metabolomics PLIER analysis representing the within-tissue z-scores per metabolite for each participant at every tissue-time point.

**Extended Data Figure 7.**
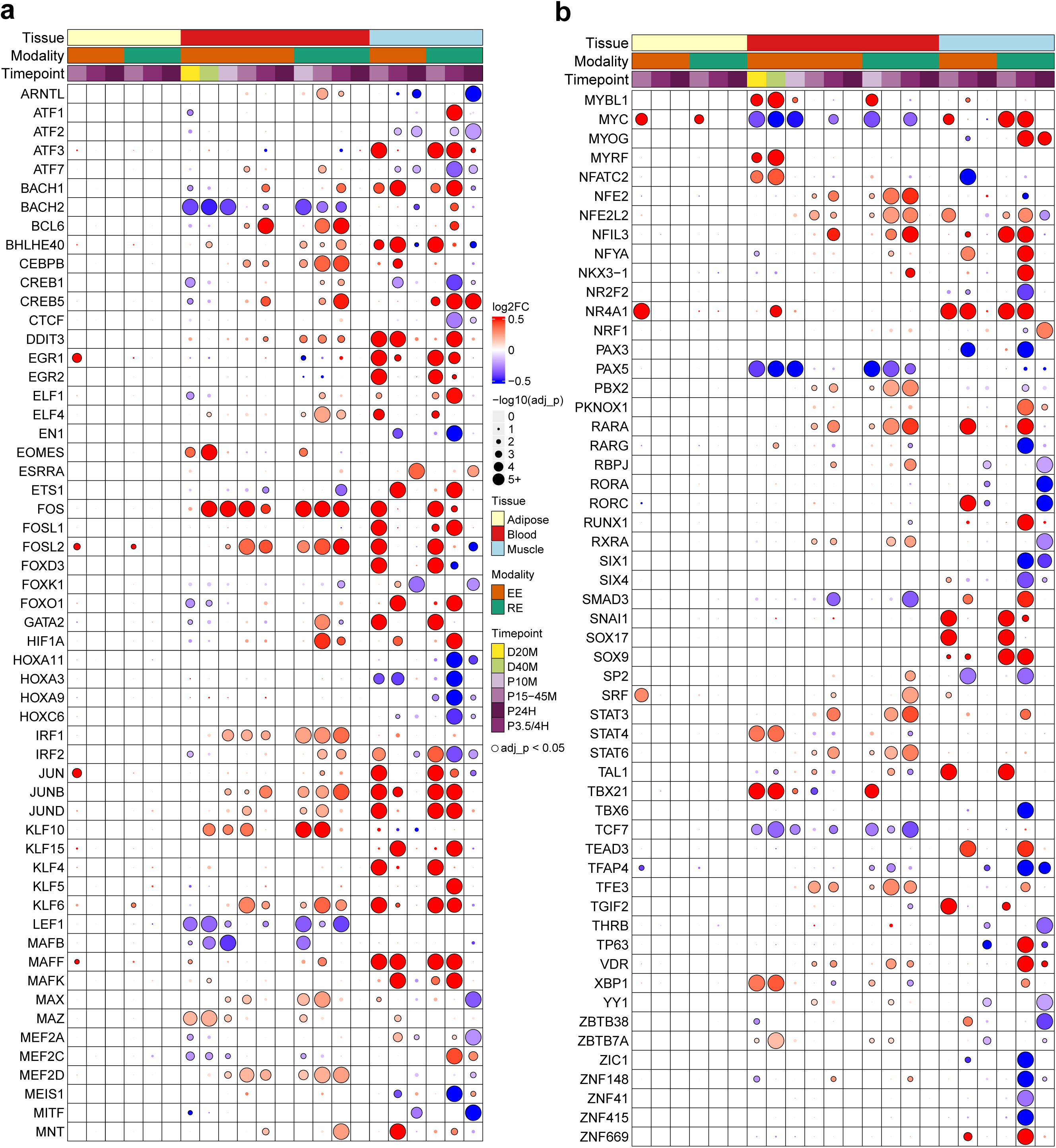
Differential transcription factor expression, related to Figure 5. **a**,**b**, Bubble heatmap of TFs with significant responses satisfying a threshold of adj p-val < 1e-05 at the RNAseq level to EE or RE in at least one time point/modality. Color reflects L2FC of response for each time point/modality comparison and size of dot reflects adjusted p-value. The plot is divided in half to more easily view the large number of TFs with significant responses with the first 56 in (a) and the final 55 in (b).

**Extended Data Figure 8.**
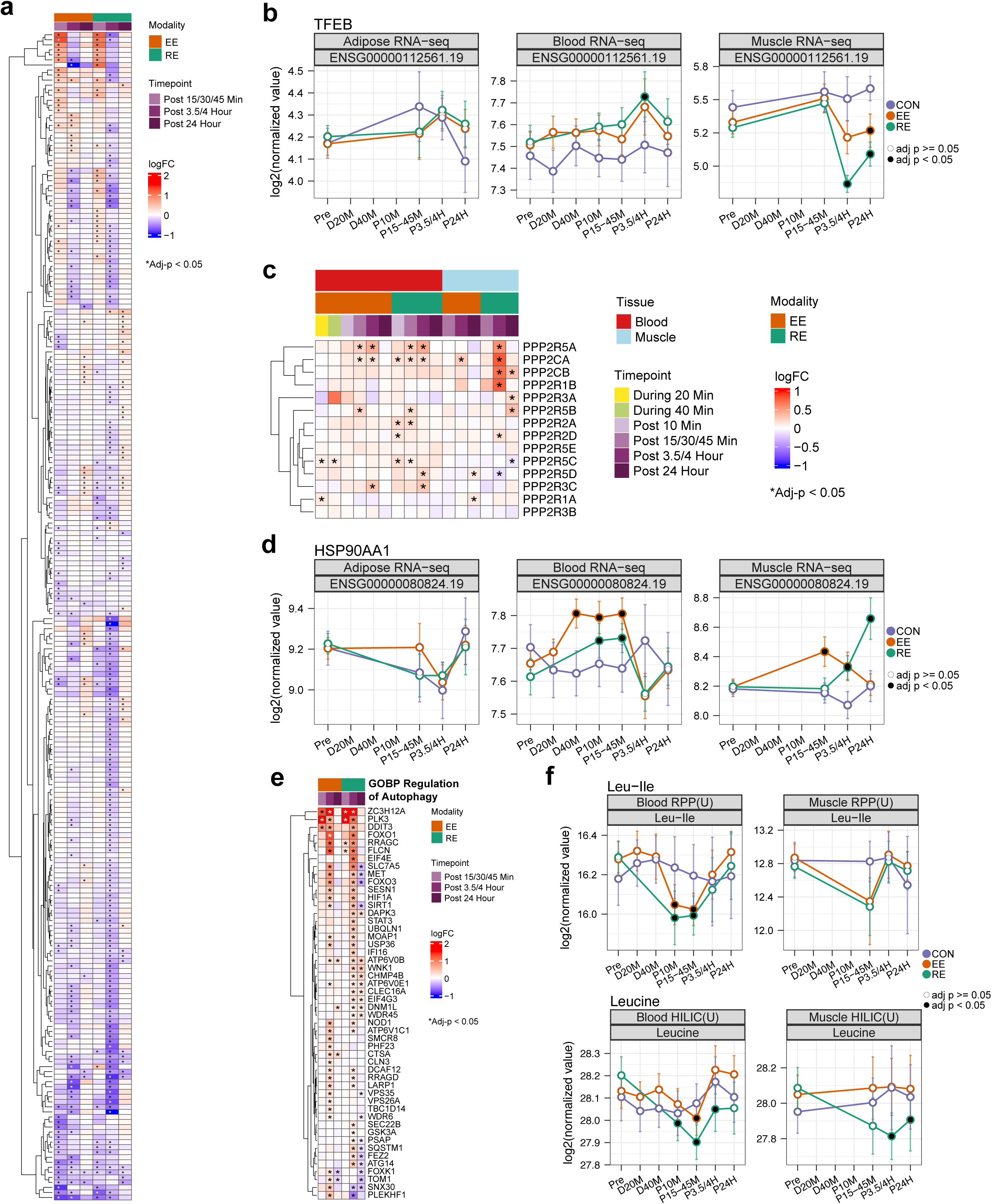
TFEB pathway regulation in response to EE and RE, related to Figure 6. **a**,Heatmap of exercise-induced RNA-seq log2 fold-changes for TFEB target genes identified by SC-ION and supported by literature ChIP-binding evidence. **b**,Temporal expression of TFEB transcript in response to exercise across the three tissues. Data are shown as group-timepoint mean ± 95% confidence interval with a black circle indicating significance relative to control at FDR<0.05. **c**, Heatmap of RNA-seq expression of PP2A subunits quantified in both blood and SKM. **d**, Temporal expression of HSP90AA1 transcript in response to exercise across the three tissues. Data are shown as group-timepoint mean ± 95% confidence interval with a black circle indicating significance relative to control at FDR<0.05. **e**, Transcriptome expression of ChIP-supported TFEB targets which are in the GO-BP Regulation of Autophagy gene set. **f**, Temporal expression of Leu-Ile and Leucine metabolites in response to exercise in blood. Data are shown as group-timepoint mean ± 95% confidence interval with a black circle indicating significance relative to control at FDR<0.05.

**Extended Data Figure 9.**
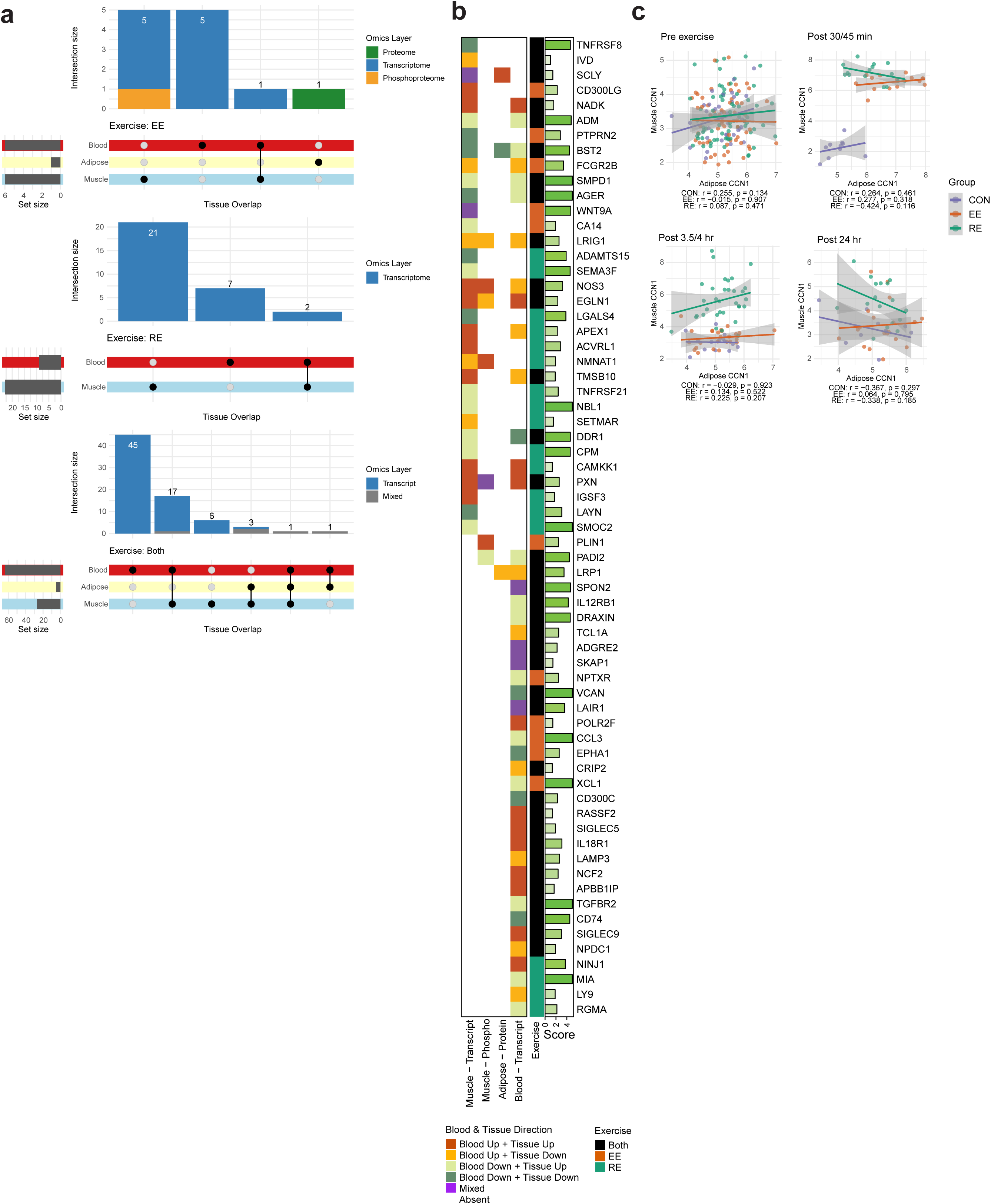
Extended secreted factor analysis and cross-tissue CCN1 correlation, related to Figure 7. **a**, UpSet plots showing global and modality-specific features differentially regulated in tissues (adjusted p < 0.05) that are also differentially regulated in plasma proteomes (adjusted p < 0.05) for EE only (top panel), RE only (middle panel) and both EE and RE (bottom panel). For AT and SKM, gene symbols from transcriptomic, proteomic, and phosphoproteomic layers were concatenated; for blood, transcriptomic data were used. Post-exercise time points were collapsed for each upset plot. Edges indicate the number of features contributing to each set, and intersecting lines represent shared differential abundance across multiple tissues or layers. ‘Mixed’ indicates a feature whose contributing omics layer differed across the tissues in that intersection (e.g., DA at the transcript level in SKM but the protein level in AT). **b**, Heatmap showing differentially regulated features across SKM, AT, and blood transcriptomes, proteomes, and phosphoproteomes (adjusted p < 0.05 at post 15-45 min, 3.5-4 hour, and 24 hour post-exercise) that also exhibit significant upregulation in plasma proteomes (adjusted p < 0.05 at any time point, including during and post-exercise) that were not included in the Figure 7a. Gene symbols from plasma proteome were used to map to corresponding features in tissue transcriptome and phosphoproteome. “Blood up” and “Blood down” indicate significant upregulation and downregulation in blood proteome at one or more time points, respectively; “Tissue up” and “Tissue down” indicate significant upregulation or downregulation in tissue omics at one or more time points, respectively. Horizontal green bars denote the extracellular score (range 0–5) from COMPARTMENTS (see Methods); “Both” in the Exercise column indicates features differentially regulated in both EE and RE. **c**, Correlation plots of *CCN1* transcript expression in each group (CON, EE, RE) between adipose and muscle at each time point (Pre-exercise, post 30/45 min, post 3.5/4 hour, and post 24 hour). Pearson correlation and p-values are assessed for each comparison.

**Extended Data Table 1.** Selected molecular features associated with baseline clinical traits and acute exercise.

| Clinical trait | Tissue | Ome | Molecule name | Baseline trait ass. (t_stat) | Adj p-value (t_stat) | Baseline trait ass. (beta) | EE response (logFC) | Adj p-value (EE) | RE response (logFC) | Adj p-value (RE) | Time point | EE vs RE Diff | Molecule Info |
| --- | --- | --- | --- | --- | --- | --- | --- | --- | --- | --- | --- | --- | --- |
| HOMA-IR | Adipose | RNA-seq | CCN2 | 3.49 | 0.013 | 0.111 | 1.29 | <0.001 | 1.34 | <0.001 | P15-45M | NO | Matricellular protein regulating cell adhesion |
| Legpress | Blood | RNA-seq | URB1 | -5.25 | 0.003 | -0.001 | -0.35 | 0.004 | -0.23 | 0.046 | P3.5/4H | NO | Nucleolar protein involved in ribosome production and rRNA processing |
| Legpress | Blood | RNA-seq | CEP68 | -4.4 | 0.025 | -0.001 | -0.23 | 0.024 | NA | NA | D20M | NA | Protein involved in centrosome organization and cilia formation |
| Legpress | Blood | RNA-seq | CSKMT | -4.33 | 0.03 | -0.003 | 0.64 | 0.025 | NA | NA | D40M | NA | Mitochondrial enzyme; methylates citrate synthase |
| NEFA | Muscle | RNA-seq | ANGPTL4 | 6.78 | <0.001 | 3.074 | 1.56 | 0.001 | 1.11 | 0.011 | P3.5/4H | NO | Secreted glycoprotein regulating metabolism |
| NEFA | Blood | RNA-seq | CPT1A | 6.01 | <0.001 | 0.63 | 0.21 | 0.001 | NA | NA | D40M | NA | Fatty acid transporter in mitochondrial membrane |
| NEFA | Muscle | RNA-seq | DBP | -4.28 | 0.033 | -0.59 | 0.56 | <0.001 | -0.27 | 0.108 | P3.5/4H | YES | Transcription factor involved in circadian clock regulation |
| NEFA | Muscle | RNA-seq | PDK4 | 5.43 | 0.001 | 2.155 | 1.51 | <0.001 | 2.55 | <0.001 | P3.5/4H | YES | Kinase that inhibits the pyruvate dehydrogenase complex |
| NEFA | Muscle | RNA-seq | PFKFB4 | -4.15 | 0.034 | -0.521 | -0.012 | 0.973 | -0.35 | 0.035 | P3.5/4H | YES | Bifunctional kinase/phosphatase regulating fructose-2,6-bisphosphate |
| NEFA | Muscle | RNA-seq | PREB | 5.24 | 0.002 | 0.491 | 0.47 | <0.001 | 0.27 | 0.003 | P24H | YES | Transcriptional and protein transport regulator |
| NEFA | Muscle | RNA-seq | TXNIP | 5.98 | <0.001 | 1.05 | -0.93 | <0.001 | -0.71 | <0.001 | P3.5/4H | NO | Regulator of cellular redox signaling |
| VO<sub>2</sub>peak | Muscle | RNA-seq | CKB | 5.37 | <0.001 | 0.78 | 0.505 | 0.039 | -0.25 | 0.318 | P24H | YES | Creatine kinase |
| VO<sub>2</sub>peak | Muscle | RNA-seq | FUT1 | 3.47 | 0.022 | 0.28 | 1.78 | <0.001 | 1.46 | <0.001 | P15-45M | NO | Fucosyltransferase |
| VO<sub>2</sub>peak | Muscle | RNA-seq | HSPB1 | -4.69 | 0.001 | -0.32 | -0.34 | 0.066 | 0.97 | <0.001 | P24H | YES | Heat shock protein affecting protein folding |
| VO<sub>2</sub>peak | Muscle | RNA-seq | NCL | -3.69 | 0.014 | -0.0714 | -0.07 | 0.347 | -0.136 | 0.006 | P24H | NO | Nucleolin involved in synthesis and maturation of ribosomes |
| VO<sub>2</sub>peak | Muscle | RNA-seq | PDLIM3 | -5.39 | <0.001 | -0.28 | -0.23 | 0.062 | 0.59 | <0.001 | P24H | YES | Cytoskeletal protein |
| VO<sub>2</sub>peak | Muscle | RNA-seq | PITPNM1 | 6.18 | <0.001 | 0.59 | 0.38 | 0.01 | 0.48 | <0.001 | P24H | NO | Phosphatidylinositol transfer protein |
| VO<sub>2</sub>peak | Muscle | RNA-seq | PPARGC1A | 3.39 | 0.026 | 0.21 | 3.02 | <0.001 | 3.38 | <0.001 | P3.5/4H | YES | Transcriptional coactivator regulating metabolic gene expression |
| VO<sub>2</sub>peak | Muscle | RNA-seq | PRRT4 | -3.16 | 0.039 | -0.38 | 0.22 | 0.538 | 1.41 | <0.001 | P24H | YES | Membrane protein |
| VO<sub>2</sub>peak | Muscle | RNA-seq | SRSF11 | -3.72 | 0.013 | -0.072 | -0.23 | <0.001 | -0.09 | 0.128 | P15-45M | YES | Splicing factor |

## Supplementary Information

Supplementary Table 1. Participant numbers, related to Figure 1

Supplementary Table 2. Detailed differential analysis results, related to Figure 2

Supplementary Table 3. Direct comparison of EE vs. RE, association of molecular changes to clinical phenotypes, related to Figure 3

Supplementary Table 4. Temporal molecular response analysis, related to Figure 4

Supplementary Table 5. Homer analysis results, related to Figure 5

Supplementary Table 6. SC-ION results, related to Figure 6

Supplementary Table 7. Exerkine candidate list, related to Figure 7

## Methods

### Experimental model and study participant details

#### IRB

The Molecular Transducers of Physical Activity Consortium (MoTrPAC) (NCT03960827) is a multicenter study designed to isolate the effects of structured exercise training on the molecular mechanisms underlying the health benefits of exercise and physical activity. Methods are described here in sufficient detail to interpret results, with references to supplementary material and prior publications for detailed descriptions. The present work includes data from prior to the suspension of the study in March 2020 due to the Covid-19 pandemic. The study was conducted in accordance with the Declaration of Helsinki and approved by the Institutional Review Board of Johns Hopkins University School of Medicine (IRB protocol # JHMIRB5; approval date: 05/06/2019). All MoTrPAC participants provided written informed consent for the MoTrPAC study indicating whether they agreed to open data sharing and at what level of sharing they wanted. Participants could choose to openly share all de-identified data, with the knowledge that they could be reidentified, and they could also choose to openly share limited de-identified individual level data, which is lower risk of re-identification. All analyses and resulting data and results are shared in compliance with the NIH Genomic Data Sharing (GDS) policy and DSMB requirements for the randomized study. The analysis of data from the present subset of participants, enrolled prior to the Covid-19 suspension, was approved by the DSMB on July 19, 2021.

#### Participant Characteristics

Volunteers were screened to: (a) ensure they met all eligibility criteria; (b) acquire phenotypic assessments of the study population; and (c) verify they were healthy and medically cleared to be formally randomized into the study (cite Clinical Landscape).^37^ Participants were then randomized into intervention groups stratified by clinical site and completed a pre-intervention baseline acute test. A total of 176 participants completed the baseline acute test (EE=66, RE=73, and CON=37), and among those, 175 (99%) had at least one biospecimen sample collected. Participants then began 12 weeks of exercise training or control conditions. Upon completion of the respective interventions, participants repeated phenotypic testing and the acute test including biospecimen sampling. Due to the COVID-19 suspension, only 45 participants (26%) completed the post-intervention follow-up acute exercise test (some with less than 12 weeks of training), with 44 completing the post-intervention biospecimen collections. See *Clinical Landscape* for further detail.

#### Participant Assessments

As described in *Clinical Landscape and protocol paper*, prior to randomization, participant screening was completed via questionnaires, and measurements of anthropometrics, resting heart rate and blood pressure, fasted blood panel, cardiorespiratory fitness, muscular strength, and free-living activity and sedentary behavior.(ref Clinical landscape, in preparation/submission)^37^ Grip strength of the dominant hand was obtained using the Jamar Handheld Hydraulic Dynamometer (JLW Instruments, Chicago, IL). See *Clinical Landscape* for participant assessment results.

#### Acute Exercise Intervention

The pre-intervention baseline EE acute bouts (Fig. 1a) were composed of three parts: (1) a 5 minute warm up at 50% of the estimated power output to elicit 65% VO_2_peak, (2) 40 minute cycling at ∼65% VO_2_peak, and (3) a 1 minute recovery at ∼25 W. The pre-intervention baseline RE acute bouts were composed of a 5 minute warm-up at 50-60% heart rate reserve (HRR) followed by completion of three sets each of five upper (chest press, overhead press, seated row, triceps extension, biceps curl) and three lower body (leg press, leg curl, leg extension) exercises to volitional fatigue (∼10RM) with 90sec rest between each set. Participants randomized to CON did not complete exercise during their acute tests. Participants rested supine for 40 minutes to mirror the EE and RE acute test schedule. See *Clinical Landscape* for acute bout results.

#### Biospecimen Collection

To standardize conditions prior to the acute test, participants were instructed to comply with a variety of controls related to COX-inhibitors, biotin, caffeine, alcohol, exercise, and final nutrition consumption.^37^ Blood, muscle, and adipose samples were collected for the acute test at specific timepoints before, during, and after exercise (Fig. 1b).

Participants arrived fasted in the morning, and rested supine for at least 30 minutes prior to pre-exercise biospecimen collections. All participants had pre-exercise sampling from all three tissues. After the pre-exercise biospecimen collections, the subsequent collection timepoints varied for each training group and their randomized temporal profile (early, middle, late, all). Temporal profiles were used to reduce the burden of repeat sampling while still maintaining biochemical coverage of the full post-exercise period (Fig. 1b, Supplementary Table 1). During and after exercise, biospecimen collection varied by randomized group, tissue, and temporal group. At 20 and 40 minutes of exercise, blood was collected from only EE and CON groups due to logistical limitations in obtaining blood during RE. Immediately post-exercise, at 10 minutes, a blood sample was collected in all three groups. Finally, samples from all three tissues were collected at early (SKM at 15, blood at 30, and AT at 45 minutes post-exercise), intermediate (blood and SKM at 3.5, and AT at 4 hours post-exercise), and late (24 hours post-exercise in all tissues) timepoints. The post-exercise timepoint began at the completion of the 40 minutes of cycling, third set of leg extension, or 40 minutes of rest, depending on the randomized group. Except for the EE blood collection timepoints during exercise, participants rested supine or seated for biospecimen collections. The collection and processing protocols for each tissue were previously described in the human clinical protocol design paper.^37^ See *Clinical Landscape* for biospecimen collection success results.

The number of biospecimen samples that were profiled on any given assay varied by tissue, timepoint, and randomized intervention group for feasibility reasons. Multi-omic coverage is summarized in Fig. 1c, Extended Data Fig. 1b, and Supplementary Table 1.

#### Post training intervention data

As mentioned, 44 of the 175 individuals who began the trial and had biospecimens collected did complete at least a portion of the training intervention period and the post-training acute bout session. These biospecimens were collected, and molecular data were generated for them. While these data are to be shared with the scientific community as detailed in the MoTrPAC Data Sharing Plan, the data are not analyzed here due to the low sample size and potential confounding between participant characteristics and groups of interest. Rather, the focus is on the baseline acute bout data. The full cohort dataset will allow exploration of multiple hypotheses related to training effects and is forthcoming.

### Molecular Quantification Methods

#### Sample identification quality control

To detect sample mislabeling and sex discordance across multi-omic datasets, we employed Somalier (v0.3.1)^127^ on BAM files from WGS, RNA-seq, and ATAC-seq samples using the RNA-seq sites VCF (sites.hg38.rna.vcf.gz). Somalier computes pairwise relatedness (ranging from 0 to 1) and identity-by-state (IBS) metrics to verify sample identity across tissues, time points, and assay types. Specifically, IBS0 represents the number of sites where one sample is homozygous-reference and another is homozygous-alternative, while IBS2 represents sites where samples share identical genotypes. Samples originating from the same individual were expected to exhibit high relatedness (>0.9), low IBS0, and high IBS2 values.

Sex concordance was verified using complementary approaches tailored to each data type. For WGS and ATAC-seq, genetic sex was inferred directly using Somalier. For RNA-seq data, sex was determined using a custom expression-based script leveraging well-established sex-linked markers (e.g., high XIST expression with low or absent DDX3Y, RPS4Y1, KDM5D, and UTY expression in females).^128,129^ We developed an integrated quality control pipeline combining Somalier metrics with sex concordance checks to systematically identify mislabeled samples. Samples were flagged as potentially mislabeled if they demonstrated very low relatedness (<0.2) and elevated IBS0 values relative to all other samples from the same participant across time points, tissues, and assays. This pipeline identified six potentially mislabeled ATAC-seq samples, which were excluded from downstream analyses. No mislabeled samples or sex discordances were detected in the WGS or RNA-seq datasets.

#### Whole Genome Sequencing (WGS)

In the present cohort, 174 of 175 participants had Illumina whole genome sequencing (WGS). DNA was extracted from cell pellets obtained from the whole blood collected in EDTA tubes using QIAamp DNA Mini Kit (QIAamp #51104) by following the manufacturer’s recommendations (Qiagen). WGS libraries were prepared using the Truseq DNA PCR-free Library Preparation Kit (Illumina # FC-121-3003) in accordance with the manufacturer’s instructions. Briefly, 1 μg of DNA was sheared using a Covaris LE220 sonicator (adaptive focused acoustics). DNA fragments underwent bead-based size selection and were subsequently end-repaired, adenylated, and ligated to Illumina sequencing adapters. Final libraries were quantified using the Qubit Fluorometer (Life Technologies) or Spectromax M2 (Molecular Devices) and Fragment Analyzer (Advanced Analytical) or Agilent 2100 BioAnalyzer. Libraries were sequenced to 30x coverage using 2×100bp paired-end run on a NovaSeq 6000 sequencer on an S4 flow cell according to the manufacturer’s instructions (Illumina). Sequenced reads were aligned to the GRCh38 reference genome (hs38DH, which includes alternative sequences, HLA regions, and decoy sequences) using the Burrows-Wheeler Aligner (BWA-MEM v0.7.15).^130^ Post-alignment processing included fixing mate pairs, base quality recalibration, and variant calling and filtration, all performed according to GATK best practices workflow that includes marking of duplicate reads by the use of Picard tools (v2.4.1),^131^ and base quality score recalibration (BQSR) via Genome Analysis Toolkit (GATK v3.5).^132,133^ Variant annotation was conducted using the Variant Effect Predictor (VEP),^134^ which provides information on variant allele frequencies from gnomAD,^135^ predicts the impact of nucleotide changes on protein sequences, and identifies variant-disease associations. Quality control of the data and variant calls were performed using Picard (CollectVariantCallingMetrics) and samtools.^131,136^ Joint genotyping was carried out on individual sample GVCFs to produce highly sensitive variant calls for the entire cohort.

#### ATAC-sequencing

ATAC-seq (Assay for Transposase Accessible Chromatin using sequencing) assays were divided by tissue, with SKM tissue assayed at Icahn School of Medicine at Mount Sinai and blood PBMCs assayed at Stanford University. Samples for ATAC PBMC processing were randomized as described in https://github.com/MoTrPAC/clinical-sample-batching. SKM samples were processed for ATAC-seq by study group (CON, EE, and RE). Eight to 12 samples were processed at a time for nuclear extraction and library preparation, and all the samples in the study group were sequenced together. For muscle, nuclei from aliquoted tissue samples (10 mg for muscle tissues) were extracted using the Omni-ATAC protocol with modifications. Two tissue-specific consortium reference standards were included for sample processing QC. 500-600 ul 1X homogenization buffer with protease inhibitor was added to the sample micronic tubes plus three 1.8 mm ceramic bead media. Tubes were vortexed for 5-10 seconds 3 times and kept on ice. The mixture was homogenized using Bead Ruptor Elite (Bead Mill Homogenizer) and Omni BR Cryo Cooling Unit at the settings: Tube 1.2 ml, Speed: 1.0, Cycles: 2, Time: 20 Sec, Dwell Time: 20 Sec, Temperature: 4-8⁰C. The homogenized mixture was filtered with Mini 20 um Pluristrainer and waited for 20-30 seconds. If the homogenate did not pass through the filter easily, the homogenate tubes were briefly centrifuged with filters attached. Nuclei were stained with Trypan blue dye 0.4% and counted using Countess II Automated Cell Counter. Depending upon the counts, the number of nuclei to use for transposition was calculated. For human pre-COVID muscle samples, 250K-300K nuclei were used. Calculated nuclei were added to 1 ml RSB in 2.0 ml Eppendorf tubes, mixed well and centrifuged at 510 RCF for 10 minutes on fixed angle centrifuge at 4⁰C. 70% supernatant was removed without disturbing pellet at the bottom and nuclei were again centrifuged at 510 RCF for 5 minutes. Almost all of the supernatant was removed. A 50 μl transposition mix was added to the pellet and mixed by pipetting 6 times, then incubated at 37⁰C on a thermomixer with 1000 rpm for 30 minutes and immediately transferred to ice after incubation. The transposed DNA was purified using Zymo DNA Clean and Concentrator kit (Zymo Research, D4014). We amplified the DNA product using NEBnext High-Fidelity 2x PCR Master Mix (NEB, M0541L) and custom-indexed primers. Purified PCR libraries were amplified using Ampure XP beads, double-sided bead purification (sample to beads ratios 0.5X and 1.3X).

Whole blood was collected in a BD Vacutainer Cell Preparation Tube and processed for PBMC extraction and purification. Isolated PBMCs were frozen at −80°C in CoolCell Containers (Corning) and stored in liquid nitrogen. Frozen PBMCs were thawed in a 37°C water bath for 2-3 min. An aliquot of PBMC were stained with DAPI and counted using Countess III Automated Cell Counter. An aliquot of 50,000 cells was added to 1 ml of ATAC-RSB and spun at 1000 g for 10 minutes at 4⁰C. The supernatant was removed without disturbing the pellet. The nuclei pellet was resuspended by pipetting in 50 μL of transposition mixture (2.5μL Transposase, 25 μL 2x tagmentation buffer, 16.5 μL PBS, 0.5 μL 1% digitonin, 0.5 μL 10% Tween-20, 5 μL water) and incubated at 37°C on a thermomixer for 30 minutes with 1000 rpm shaking. The transposed DNA was purified using Qiagen MinElute Purification kits (Qiagen, 28006), and amplified using NEBnext High-Fidelity 2x PCR Master Mix (NEB, M0541L) and custom indexed primers. 90 μL SPRIselect beads (beads : DNA = 1.8) were used to purify the PCR reaction and remove excess primers and primer dimers.

#### ATAC-seq library sequencing and data processing

Pooled libraries were sequenced on an Illumina NovaSeq 6000 platform (Illumina, San Diego, CA, USA) using a paired-end 100 base-pair run configuration to a target depth of 25 million read pairs (50 million paired-end reads) per sample for muscle and to a target depth of 35 million read pairs for PBMCs. We demultiplexed reads with bcl2fastq2 (v2.20.0) (Illumina, San Diego, CA, USA). Data was processed with the ENCODE ATAC-seq pipeline (v1.7.0) (https://github.com/ENCODE-DCC/atac-seq-pipeline). Samples from a single sex, group and exercise timepoint, e.g., endurance exercising males 30 minutes post-exercise bout, were analyzed together as biological replicates in a single workflow. Briefly, adapters were trimmed with cutadapt (v2.5) and aligned to human genome hg38 with Bowtie 2 (v2.3.4.3).^137^ Duplicate reads and reads mapping to the mitochondrial chromosome were removed. Signal files and peak calls were generated using MACS2 v2.2.4,^138^ both from reads from each sample and pooled reads from all biological replicates. We compared pooled peaks with peaks called for each replicate individually using the Irreproducibility Discovery Rate and thresholded them to generate an optimal set of peaks. The cloud implementation of the ENCODE ATAC-seq pipeline and source code for the post-processing steps are available at https://github.com/MoTrPAC/motrpac-atac-seq-pipeline. Optimal peaks (overlap.optimal_peak.narrowPeak.bed.gz) from all workflows were concatenated, trimmed to 200 base pairs around the summit, and sorted and merged with bedtools v2.29.0 to generate a master peak list.^139^ This peak list was intersected with the filtered alignments from each sample using bedtools coverage with options -nonamecheck and -counts to generate a peak by sample matrix of raw counts. The remaining steps were applied separately on raw counts from each tissue. Peaks from non-autosomal chromosomes were removed, as well as peaks that did not have at least 10 read counts in six samples, which aided in increasing significant peak output in differential analysis. Principal component analysis (PCA) was conducted to find sample outliers to be excluded from downstream analysis using the *call_pca_outliers* command (found in the https://github.com/MoTrPAC/MotrpacRatTraining6mo/ R package).^31^ No PCA outliers were identified. Filtered raw counts were then quantile-normalized with variancePartition::voomWithDreamWeights,^140^ which is akin to limma::voom,^141^ with the added ability to include random effects in the formula. Four samples in the blood (Peripheral Blood Mononuclear Cell) ATAC-seq and 2 samples in SKM ATAC-seq were flagged as potential sample mislabels based on Somalier analysis,^127^ and were removed from downstream analysis. See ‘Sample identification quality control’ above for further details. See Supplementary Results for an overview of experimental limitations in the muscle ATAC-seq data.

#### Methylation Capture sequencing

##### Data production

Plates for methylation capture were batched for methylation capture processing by exercise group. Samples were processed in subject-level batches of 4 to make 1 library pool, and all timepoints for a subject were pooled together when possible. Library pools (4 samples) were randomized into sequencing pools (40 total samples) to distribute the study groups.

Genomic DNA from muscle, adipose, and blood collected in EDTA tubes was extracted using the GenFind v3 kit (Beckman Coulter) in an automated workstation (Biomek Fxp) according to the manufacturer’s instructions. Two tissue-specific control samples (provided by the Consortium) were included to monitor the sample processing QC. The DNA was quantified by Qubit assay (dsDNA BR assay, Thermo Fisher Scientific) and the quality was determined by the Nanodrop A260/280 and A260/230 ratios. A260/280 and A260/230 ratios were used to determine the quality of the DNA. DNA samples with a A260/A230 ratio above 1.8 were used for the assay.

The Methyl-Cap sequencing libraries were prepared using the Illumina TruSeq Methyl Capture EPIC Library Prep Kit (Cat# FC-151-1003) according to the manufacturer’s instructions (https://www.illumina.com/products/by-type/sequencing-kits/library-prep-kits/truseq-methyl-capture-epic.html). Briefly,1 μg of genomic DNA in 50 μL was fragmented using a Covaris E220 ultrasonicator (Covaris, USA) and the fragment size was assessed by running 1.0 μL of the samples on a Bioanalyzer High Sensitivity DNA chip (Agilent Technologies). Samples with DNA fragments between 100-300 bp with a peak between 155–175 bp were used for library preparation. Fragmented DNA was end repaired into dA-tailed fragments and ligated with custom xGen™ Methyl UDI-UMI Adapters (Integrated DNA Technologies). Adapter-ligated DNA was purified by magnetic bead separation. The DNA samples were then pooled in groups of 4 in equal volume (10 μL/samples, 40 μL final volume) and hybridized with Illumina EPIC probe sets (covering >3.3 million targeted CpG sites). The DNA fragments hybridized with probes were captured by streptavidin-magnetic beads and purified. The enriched DNA underwent a second round of hybridization and purification to ensure high specificity of the captured regions. Bisulfite conversion was performed on the captured DNA libraries and PCR amplified. The purified libraries were analyzed with the 2100 Bioanalyzer High Sensitivity DNA assay (Agilent Technologies) and quantified by Qubit assay (ThermoFisher). The libraries contained fragment size between 230 - 450 bp with a peak around 250 - 300 bp.

The libraries were pooled and 15% PhiX DNA (Illumina) was added to each library pool to boost diversity. DNA sequencing of paired-end reads (2 x 100 bp reads) were performed on an Illumina Novaseq 6000 system (Illumina Inc.) at a depth of approximately 60 million reads per library. In order to capture the 9-base UMIs, libraries were sequenced using 17 cycles for i7 index read and 8 cycles for the i5 index read.

##### Methylation data processing, quantification and QC

The 17 cycles for the i7 index read in the libraries consist of 8 cycles of sample i7 barcode index and 9 cycles of UMI. Using the mask options --use-bases-mask Y*,I8Y*,I*,Y* --mask-short-adapter-reads 0 --minimum-trimmed-read-length 0 in bcl2fastq command to extract the two read (R1,R2) FASTQ files, and index FASTQ files that contain the UMI information. The three FASTQ files are then fed into the methylcap processing pipeline as described in https://github.com/MoTrPAC/motrpac-methyl-capture-pipeline. Briefly, the pipeline first attaches the 9 cycles of the UMI index FASTQ file as part of the read names in both R1 and R2 FASTQ files and then uses trimgalore (via cutadapt v1.18)^142^ to remove the adapters. The FASTQ files are aligned with bismark (v0.20.0)^143^ and the resulting bam files are deduplicated using the bismark command deduplicate_bismark with option “--barcode”. The bismark coverage from the adjacent neighboring positions of the two opposite strands are merged with the bismark command coverage2cytosine with the option “--merge_CpG”.

NGScheckmate (v1.01) was applied to the deduplicated bam files from the Bismark pipeline to check the within-subject genotype consistency.^144^ The percentage of reads mapped to chrX and chrY are also checked against with the sex annotation of the sample. The inconsistent samples from genotype and sex checking and bismark QC metrics (two low read depth, two high %CHH) outliers were removed from further analysis.

For each tissue (EDTA, Muscle, Adipose) separately, the bismark coverage from all samples are combined into a single R data file after removing CpG sites with 5X coverage in less than 50% of the samples. To merge highly correlated nearby sites into a region, Markov Cluster Algorithm (MCL) is then applied on the bismark coverage R data file for each tissue separately,^31,145,146^ with the following decision algorithm: a) For each sample, compute nCpG10, the number of CpG sites with at least 10X coverage. b) Filter out samples with less than 20 percentile of the nCpG10. c) apply MCL on the CpG sites with at least 10X coverage on all of the remaining samples. d) also apply the CpG merging results from c) to the filtered out samples in b).

Principal component analysis (PCA) was conducted to find sample outliers to be excluded from downstream analysis using the *call_pca_outliers* command (found in the https://github.com/MoTrPAC/MotrpacRatTraining6mo/ R package). No additional outliers were detected from PCA in this ome.

##### Methylation Statistical analysis

To accommodate the within subject mixed effects model for the differential methylation analysis, the coverage of all of the CpG sites (or CpG regions with merged CpG sites obtained from the MCL) is applied to Malax to obtain the p-value.^147^ The statistical comparison of interest compares the change from pre-exercise to a during or post-exercise timepoint in one of the two exercise groups (RE and EE) to the same change in control. Due to the limited sample sizes in the methylation data, only sex and age are the additional covariates in the statistical model. In order to reduce the computational time, the CpG sites/regions are divided into 100 small data sets for the Malax analysis using cluster computers for parallel computing and the adjusted p-values are computed on the final merged dataset. After hypothesis testing using Malax, the change in absolute methylation percentage is calculated using edgeR.^148^

#### Transcriptomics

RNA Sequencing (RNA-Seq) was performed at Stanford University and the Icahn School of Medicine at Mount Sinai. Processing randomization for blood, muscle was done according to https://github.com/MoTrPAC/clinical-sample-batching. See below for adipose randomization considerations.

##### Extraction of total RNA

Tissues (∼10 mg for muscle, ∼50 mgs for adipose) were disrupted in Agencourt RNAdvance tissue lysis buffer (Beckman Coulter, Brea, CA) using a tissue ruptor (Omni International, Kennesaw, GA, #19-040E). Total RNA was extracted in a BiomekFX automation workstation according to the manufacturer’s instructions for tissue-specific extraction. Total RNA from 400 μL of blood collected in PAXgene tubes (BD Biosciences, Franklin Lakes, NJ, # 762165) was extracted using the Agencourt RNAdvance blood specific kit (Beckman Coulter). Two tissue-specific consortium reference standards were included to monitor the sample processing QC. The RNA was quantified by NanoDrop (ThermoFisher Scientific, # ND-ONE-W) and Qubit assay (ThermoFisher Scientific), and the quality was determined by either Bioanalyzer or Fragment Analyzer analysis.

##### mRNA Sequencing Library Preparation

Universal Plus mRNA-Seq kit from NuGEN/Tecan (# 9133) were used for generation of RNA-Seq libraries derived from poly(A)-selected RNA according to the manufacturer’s instructions. Universal Plus mRNA-Seq libraries contain dual (i7 and i5) 8 bp barcodes and an 8 bp unique molecular identifier (UMI), which enable deep multiplexing of NGS sequencing samples and accurate quantification of PCR duplication levels. Approximately 500ng of total RNA was used to generate the libraries for muscle, 300ng for adipose, and 250ng of total RNA was used for blood. The Universal Plus mRNA-Seq workflow consists of poly(A) RNA selection, RNA fragmentation and double-stranded cDNA generation using a mixture of random and oligo(dT) priming, end repair to generate blunt ends, adaptor ligation, strand selection, AnyDeplete workflow to remove unwanted ribosomal and globin transcripts, and PCR amplification to enrich final library species. All library preparations were performed using a Biomek i7 laboratory automation system (Beckman Coulter). Tissue-specific reference standards provided by the consortium were included with all RNA isolations to QC the RNA.

##### RNA Sequencing, quantification, and normalization

RNA sequencing, quantification, and normalization Pooled libraries were sequenced on an Illumina NovaSeq 6000 platform (Illumina, San Diego, CA, USA) to a target depth of 40 million read pairs (80 million paired-end reads) per sample using a paired-end 100 base pair run configuration. In order to capture the 8-base UMIs, libraries were sequenced using 16 cycles for the i7 index read and 8 cycles for the i5 index read. Reads were demultiplexed with bcl2fastq2 (v2.20.0) using options --use-bases-mask Y*,I8Y*,I*,Y* --mask-short-adapter-reads 0 --minimum-trimmed-read-length 0 (Illumina, San Diego, CA, USA), and UMIs in the index FASTQ files were attached to the read FASTQ files. Adapters were trimmed with cutadapt (v1.18),^137^ and trimmed reads shorter than 20 base pairs were removed. FastQC (v0.11.8) was used to generate pre-alignment QC metrics. STAR (v2.7.0d)^149^ was used to index and align reads to release 38 of the Ensembl Homo sapiens (hg38) genome and Gencode (Version 29). Default parameters were used for STAR’s genomeGenerate run mode; in STAR’s alignReads run mode, SAM attributes were specified as NH HI AS NM MD nM, and reads were removed if they did not contain high-confidence collapsed splice junctions (--outFilterType BySJout). RSEM (v1.3.1)^150^ was used to quantify transcriptome-coordinate-sorted alignments using a forward probability of 0.5 to indicate a non-strand-specific protocol. Bowtie 2 (v2.3.4.3)^151^ was used to index and align reads to globin, rRNA, and phix sequences in order to quantify the percent of reads that mapped to these contaminants and spike-ins. UCSC’s gtfToGenePred was used to convert the hg38 gene annotation (GTF) to a refFlat file in order to run Picard CollectRnaSeqMetrics (v2.18.16) with options MINIMUM_LENGTH=50 and RRNA_FRAGMENT_PERCENTAGE=0.3. UMIs were used to accurately quantify PCR duplicates with NuGEN’s “nodup.py” script (https://github.com/tecangenomics/nudup). QC metrics from every stage of the quantification pipeline were compiled, in part with multiQC (v1.6). The openWDL-based implementation of the RNA-Seq pipeline on Google Cloud Platform is available on Github (https://github.com/MoTrPAC/motrpac-rna-seq-pipeline). Filtering of lowly expressed genes and normalization were performed separately in each tissue. RSEM gene counts were used to remove lowly expressed genes, defined as having 0.5 or fewer counts per million in at least 10% of samples. These filtered raw counts were used as input for differential analysis with the variancePartition::dream,^140^ as described in the statistical analysis methods section. To generate normalized sample-level data for downstream visualization, filtered gene counts were TMM-normalized using edgeR::calcNormFactors, followed by conversion to log2 counts per million with edgeR::cpm.

Principal Component Analysis and calculation of the variance explained by variables of interest were used to identify and quantify potential batch effects. Based on this analysis, processing batch (muscle and blood, see below for adipose), Clinical Site (all tissues), percentage of UMI duplication, and RNA Integrity Number (RIN) technical effects were regressed out of the TMM-normalized counts via linear regression using limma::RemoveBatchEffect function in R.^141^ A design matrix including age, sex, and a combination of group and timepoint was used during batch effect removal to avoid removing variance attributable to biological effects.

##### Adipose tissue processing considerations

In the adipose tissue, RNA extraction was performed in separate batches according to exercise modality, so the extraction batch and exercise modality were nearly collinear. This collinearity was identified at the RNA extraction step and samples were randomized prior to construction of cDNA libraries.

##### RNA Quality Inclusion Criteria

In the blood transcriptomic data, 79/1032 processed adult-sedentary samples had RIN values under 5. Based on established guidelines and internal quality control assessments, any blood samples with a RIN score below 5 were excluded from further analysis due to concerns about potential degradation artifacts.

#### Mass Spectrometry-based Proteomics and Phosphoproteomics

##### Study design

LC-MS/MS analysis of 379 muscle samples encompassing baseline and 3 post-intervention timepoints from all three groups (control, endurance and resistance) was performed at the Broad Institute of MIT and Harvard (BI) and Pacific Northwest National Laboratories (PNNL). Muscle samples were split evenly across the two sites, and a total of 14 samples were processed and analyzed at both sites to serve as cross-site replicates for evaluation of reproducibility. 46 adipose tissue samples representing baseline and 4-hours post-intervention from all three groups were analyzed at PNNL.

##### Generation of common reference

For both tissue types, a tissue-specific common reference material was generated from bulk human samples. The common reference sample for muscle consisted of bulk tissue digest from 5 individuals at 2/3 ratio of female/male. Samples were split equally between BI and PNNL, digested at each site following sample processing protocol described below, then mixed all digests from both sites and centrally aliquoted at PNNL. Common reference for adipose tissue was generated at PNNL using bulk tissue from 6 individuals representing a 4:2 ratio of female:male. 250 μg aliquots of both tissue specific common reference samples were made to be included in each multiplex (described below) and additional aliquots are stored for inclusion in future MoTrPAC phases to facilitate data integration.

##### Sample processing

Proteomics analyses were performed using clinical proteomics protocols described previously.^152,153^ Muscle and adipose samples were lysed in ice-cold, freshly-prepared lysis buffer (8 M urea (Sigma-Aldrich, St. Louis, Missouri), 50 mM Tris pH 8.0, 75 mM sodium chloride, 1 mM EDTA, 2 μg/ml Aprotinin (Sigma-Aldrich, St. Louis, Missouri), 10 μg/ml Leupeptin (Roche CustomBiotech, Indianapolis, Indiana), 1 mM PMSF in EtOH, 10 mM sodium fluoride, 1% phosphatase inhibitor cocktail 2 and 3 (Sigma-Aldrich, St. Louis, Missouri), 10 mM Sodium Butyrate, 2 μM SAHA, and 10 mM nicotinamide and protein concentration was determined by BCA assay. Protein lysate concentrations were normalized within samples of the same tissue type, and protein was reduced with 5 mM dithiothreitol (DTT, Sigma-Aldrich) for 1 hour at 37°C with shaking at 1000 rpm on a thermomixer, alkylated with iodoacetamide (IAA, Sigma-Aldrich) in the dark for 45 minutes at 25°C with shaking at 1000 rpm, followed by dilution of 1:4 with Tris-HCl, pH 8.0 prior to adding digestion enzymes. Proteins were first digested with LysC endopeptidase (Wako Chemicals) at a 1:50 enzyme:substrate ratio (2 hours, 25 °C, 850 rpm), followed by digestion with trypsin (Promega) at a 1:50 enzyme:substrate ratio (or 1:10 ratio for adipose tissue; 14 hours, 25 °C, 850 rpm). The next day formic acid was added to a final concentration of 1% to quench the reaction. Digested peptides were desalted using Sep-Pac C18 columns (Waters), concentrated in a vacuum centrifuge, and a BCA assay was used to determine final peptide concentrations. 250μg aliquots of each sample were prepared, dried down by vacuum centrifugation and stored at −80°C.

Tandem mass tag (TMT) 16-plex isobaric labeling reagent (ThermoFisher Scientific) was used for this study. Samples were randomized across the first 15 channels of TMT 16-plexes, and the last channel (134N) of each multiplex was used for a common reference that was prepared prior to starting the study (see above). Randomization of samples across the plexes within each site was done using https://github.com/MoTrPAC/clinical-sample-batching, with the goal to have all timepoints per participant in the same plex, and uniform distribution of groups (endurance, resistance, control), sex and sample collection clinical site across the plexes.

Peptide aliquots (250 μg per sample) were resuspended to a final concentration of 5 μg/μL in 200 mM HEPES, pH 8.5 for isobaric labeling. TMT reagent was added to each sample at a 1:2 peptide: TMT ratio, and labeling proceeded for 1 hour at 25°C with shaking at 400 rpm. The labeling reaction was diluted to a peptide concentration of 2 µg/µL using 62.5 μL of 200 mM HEPES and 20% ACN. 3 μL was removed from each sample to quantify labeling efficiency and mixing ratio. After labeling QC analysis, reactions were quenched with 5% hydroxylamine and samples within each multiplex were combined and desalted with Sep-Pac C18 columns (Waters).

Combined TMT multiplexed samples were then fractionated using high pH reversed phase chromatography on a 4.6mm ID x 250mm length Zorbax 300 Extend-C18 column (Agilent) with 5% ammonium formate/2% Acetonitrile as solvent A and 5% ammonium formate/90% acetonitrile as solvent B. Samples were fractionated with 96 minute separation gradient at flow rate of 1mL/minute and fractions were collected at each minute onto a 96-well plate. Fractions are then concatenated into 24 fractions with the following scheme: fraction 1 = A1+C1+E1+G1, fraction 2 = A2+C2+E2+G2, fraction 3 = A3+C3+E3+G3, all the way to fraction 24 = B12+D12+F12+H12 following the same scheme. 5% of each fraction was removed for global proteome analysis, and the remaining 95% was further concatenated to 12 fractions for phosphopeptide enrichment using immobilized metal affinity chromatography (IMAC).

Phosphopeptide enrichment was performed through immobilized metal affinity chromatography (IMAC) using Fe 3+ -NTA-agarose beads, freshly prepared from Ni-NTA-agarose beads (Qiagen, Hilden, Germany) by sequential incubation in 100 mM EDTA to strip nickel, washing with HPLC water, and incubation in 10 mM iron (III) chloride). Peptide fractions were resuspended to 0.5 μg/uL in 80% ACN + 0.1% TFA and incubated with beads for 30 minutes in a thermomixer set to 1000 rpm at room temperature. After 30 minutes, beads were spun down (1 minute, 1000 rcf) and supernatant was removed and saved as flow-through for subsequent enrichments. Phosphopeptides were eluted off IMAC beads in 3x 75 μL of agarose bead elution buffer (500 mM K2HPO4, pH 7.0), desalted using C18 stage tips, eluted with 50% ACN, and lyophilized. Samples were then reconstituted in 3% ACN / 0.1% FA for LC-MS/MS analysis (9 μL reconstitution / 4 μL injection at the BI; 12 μL reconstitution / 5 μL injection at PNNL).

##### Data acquisition

Broad Institute: Both proteome and phosphoproteome samples were analyzed on 75um ID Picofrit columns packed in-house with ReproSil-Pur 120 Å, C18-AQ, 1.9 µm beads to the length of 20-24cm. Online separation was performed on Easy nLC 1200 systems (ThermoFisher Scientific) with solvent A of 0.1% formic acid/3% acetonitrile and solvent B of 0.1% formic acid/90% acetonitrile, flow rate of 200nL/minute and the following gradient: 2-6% B in 1min, 6-20% B in 52min, 20-35% B in 32min, 35-60% B in 9min, 60-90% B in 1min, followed by a 5 minute hold at 90%, and 9 minute hold at 50%. Proteome fractions were analyzed on a Q Exactive Plus mass spectrometer (ThermoFisher Scientific) with MS1 scan across the 300-1800 m/z range at 70,000 resolution, AGC target of 3×10^6^ and maximum injection time of 5ms. MS2 scans of most abundant 12 ions were performed at 35,000 resolution with AGC target of 1×10^5^ and maximum injection time of 120ms, isolation window of 0.7m/z and normalized collision energy of 27. Phosphoproteome fractions were analyzed on Q Exactive HFX MS system (ThermoFisher Scientific) with the following parameters: MS1 scan across 350-1800 m/z mass range at 60,000 resolution, AGC target of 3×10^6^ and maximum injection time of 10ms. 20 most abundant ions are fragmented in MS2 scans with AGC of 1×10^5^, maximum injection time of 105ms, isolation width of 0.7 m/z and NCE of 29. In both methods dynamic exclusion was set to 45sec.

PNNL: For mass spectrometry analysis of the global proteome of muscle samples, online separation was performed using a nanoAcquity M-Class UHPLC system (Waters) and a 25 cm x 75 μm i.d. picofrit column packed in-house with C18 silica (1.7 μm UPLC BEH particles, Waters Acquity) with solvent A of 0.1% formic acid/3% acetonitrile and solvent B of 0.1% formic acid/90% acetonitrile, flow rate of 200nL/minute and the following gradient: 1% B for 8min, 8-20% B in 90min, 20-35% B in 13min, 35-75% B in 5min, 75-95% B in 3min, followed by a 6 minute hold at 90%, and 9 minute hold at 50%. Proteome fractions were analyzed on a Q Exactive Plus mass spectrometer (ThermoFisher Scientific) with MS1 scan across the 300-1800 m/z range at 60,000 resolution, AGC target of 3×10^6^ and maximum injection time of 20ms. MS2 scans of most abundant 12 ions were performed at 30,000 resolution with AGC target of 1×10^5^ and maximum injection time of 100ms, isolation window of 0.7m/z and normalized collision energy of 30.

For mass spectrometry analysis of the global proteome of adipose samples, online separation was performed using a Dionex Ultimate 3000 UHPLC system (ThermoFisher) and a 25 cm x 75 μm i.d. picofrit column packed in-house with C18 silica (1.7 μm UPLC BEH particles, Waters Acquity) with solvent A of 0.1% formic acid/3% acetonitrile and solvent B of 0.1% formic acid/90% acetonitrile, flow rate of 200nL/minute and the following gradient: 1-8% B in 10min, 8-25% B in 90min, 25-35% B in 10min, 35-75% B in 5min, and 75-5% in 3min. Proteome fractions were analyzed on a Q Exactive HF-X Plus mass spectrometer (ThermoFisher Scientific) with MS1 scan across the 300-1800 m/z range at 60,000 resolution, AGC target of 3×10^6^ and maximum injection time of 20ms. MS2 scans of most abundant 12 ions were performed at 30,000 resolution with AGC target of 1×10^5^ and maximum injection time of 100ms, isolation window of 0.7m/z and normalized collision energy of 30.

For mass spectrometry analysis of the phosphoproteome of both sample types, online separation was performed using a Dionex Ultimate 3000 UHPLC system (ThermoFisher) and a 25 cm x 75 μm i.d. picofrit column packed in-house with C18 silica (1.7 μm UPLC BEH particles, Waters Acquity) with solvent A of 0.1% formic acid/3% acetonitrile and solvent B of 0.1% formic acid/90% acetonitrile, flow rate of 200nL/minute and the following gradient: 1-8% B in 10min, 8-25% B in 90min, 25-35% B in 10min, 35-75% B in 5min, and 75-5% in 3min. Phosphoproteome fractions were analyzed on a Q Exactive HF-X Plus mass spectrometer (ThermoFisher Scientific) with MS1 scan across the 300-1800 m/z range at 60,000 resolution, AGC target of 3×10^6^ and maximum injection time of 20ms. MS2 scans of most abundant 12 ions were performed at 45,000 resolution with AGC target of 1×10^5^ and maximum injection time of 100ms, isolation window of 0.7m/z and normalized collision energy of 30.

##### Data searching

Proteome and phosphoproteome data from both BI and PNNL were searched against a composite protein database at the Bioinformatics Center (BIC) using the MSGF+ cloud-based pipeline previously described.^154^ This database comprised UniProt canonical sequences (downloaded 2022-09-13; 20383 sequences), UniProt human protein isoforms (downloaded 2022-09-13; 21982 sequences), and common contaminants (261 sequences), resulting in 42,626 sequences.

##### QC/Filtering/Normalization

The log2 Reporter ion intensity (RII) ratios to the common reference were used as quantitative values for all proteomics features (proteins and phosphosites). Datasets were filtered to remove features identified from contaminant proteins and decoy sequences. Datasets were visually evaluated for sample outliers by looking at top principal components, examining median feature abundance and distributions of RII ratio values across samples, and by quantifying the number of feature identifications within each sample. No outliers were detected in the muscle tissue dataset. In the adipose datasets (proteome and phosphoproteome), four samples were flagged as outliers based on inspection of the top principal components and evaluation of the raw quantitative results (median feature abundance <-1.0). These outlier samples were removed from the dataset, and the list of outlier samples can be found in Supplementary Table 1. The Log_2_ RII ratio values were normalized within each sample by median centering to zero. Principal Component Analysis and calculation of the variance explained by variables of interest were used to identify and quantify potential batch effects. Based on this analysis, TMT plex (muscle and adipose), Clinical Site (muscle and adipose), and Chemical Analysis Site (muscle only, where samples were analyzed at both BI and PNNL) batch effects were removed using Linear Models Implemented in the *limma::RemoveBatchEffect()* function in R.^141^ A design matrix including age, sex, and group_timepoint was used during batch effect removal in order to preserve the effect of all variables included in later statistical analysis. Correlations between technical replicates analyzed within and across CAS (where applicable) were calculated to evaluate intra- and inter-site reproducibility; the data from technical replicates were then averaged for downstream analysis. Finally, features with quantification in less of 30% of all samples were removed. For specific details of the process, see available code.

#### Affinity Proteomics

Plasma proteomics profiling was performed using the Olink (Uppsala, Sweden) assay. The Proximity Extension Assay (PEA) technology has been described.^155^ Briefly, a ∼20μL plasma sample is first incubated with two distinct antibodies that bind proximal epitopes on the protein target. The two antibodies are conjugated with complementary DNA oligonucleotide sequences, which come in close proximity upon target binding and subsequently hybridize (immunoreaction). The sequence is then extended by DNA polymerase, creating a unique amplicon (i.e., barcode) for each individual protein, which is then amplified by polymerase chain reaction. Every sample is spiked with three internal quality controls: (1) “Immuno controls” are non-human antigens measured by Olink that assess technical variation in all three steps of the reaction; (2) “extension controls” are composed of an antibody coupled to a unique pair of DNA-tags that are always in proximity, to produce a constant signal independent of the immunoreaction. This control is used to adjust the signal from each sample with respect to extension and amplification; (3) “amplification/detection” control is a complete double-stranded DNA amplicon that does not require either proximity binding or extension to generate a signal, specifically monitoring only the amplification/detection step. Additionally, eight external controls are added to separate wells on each sample plate. Two pooled plasma or “external sample” controls, generated by pooling several μL of plasma from samples from healthy volunteers, are used to assess potential variation between runs and plates (i.e. to calculate inter-assay and intra-assay CVs). Three negative controls (buffer only) per plate are used to monitor background noise generated when DNA-tags come in close proximity without prior binding to the appropriate protein. These negative controls set the background level for each protein assay in order to calculate the limit of detection (LOD). Finally, each plate includes a Plate Control (PC) of a separate pooled EDTA plasma sample plated in triplicate; the median of the PC triplicates is used to normalize each assay and to compensate for potential variation between runs and plates. Data values for measurements below LOD are reported for all samples.

In total, 378 MoTrPAC sedentary adult samples were assayed across 9 plates in the same batch. No samples failed Olink standard quality control and median intra-assay and inter-assay CVs were 14.5% and 30.2%, respectively. Principal component analysis was performed and demonstrated minimal effects by plate and clinical site. Sample outliers were identified as those >3x the interquartile range for at least one of the first three principal components. Four samples (1.1%) from 3 different individuals were flagged as potential outliers and excluded from analyses. Final data are presented as NPX (normalized protein expression) values, Olink’s relative protein quantification unit on a log2 scale.

#### Metabolomics and Lipidomics

The metabolomic analysis was performed by investigators across multiple *Chemical Analysis Sites* that employed different technical platforms for data acquisition. At the highest level, these platforms were divided into two classes: *Targeted* and *Untargeted*. The data generated by the untargeted platforms were further divided into Named (confidently identified chemical entities) and Unnamed (confidently detected, but no chemical name annotation) compounds.

Targeted metabolomics data were generated at 3 analysis Sites: Duke University, the Mayo Clinic, and Emory University. Duke quantified metabolites belonging to the metabolite classes Acetyl-CoA, Keto Acids, and Nucleic Acids (“acoa”, “ka”, “nuc”), Mayo quantified amines and TCA intermediates (“amines” and “tca”), while Emory quantified Oxylipins (“oxylipneg”).

The untargeted metabolomics data were generated at 3 analysis Sites: the Broad Institute, the University of Michigan, and the Georgia Institute of Technology (Georgia Tech). The Broad Institute applied Hydrophilic interaction liquid chromatography (HILIC) in the positive ion mode (hilicpos), Michigan applied reverse phase liquid chromatography in both positive and negative ion modes (“rppos” and “rpneg”) and ion-pairing chromatography in the negative mode (“ionpneg”), and Georgia Tech performed lipidomics assays using reverse phase chromatography in both positive and negative ion modes (“lrppos” and “lrpneg”).

##### LC-MS/MS analysis of branched-chain keto acids

Duke University conducted targeted profiling of branched-chain keto acid metabolites. Plasma samples containing isotopically labeled ketoleucine (KIC)-d3, ketoisovalerate (KIV)-^13^C_5_ (from Cambridge Isotope Laboratories), and 3-methyl-2-oxovalerate (KMV)-d8 (from Toronto Research Chemicals, Canada) internal standards, were subjected to deproteinization with 3M perchloric acid. Tissue homogenates were prepared at 100 mg/ml in 3M perchloric acid and 200 μL was centrifuged.

Next, 200 μL of 25 M o-phenylenediamine (OPD) in 3M HCl were added to both the plasma and tissue supernatants. The samples were incubated at 80°C for 20 minutes. Keto acids were then extracted using ethyl acetate, following a previously described protocol.^156,157^ The extracts were dried under nitrogen, reconstituted in 200 mM ammonium acetate, and subjected to analysis on a Xevo TQ-S triple quadrupole mass spectrometer coupled to an Acquity UPLC (Waters), controlled by the MassLynx 4.1 operating system.

The analytical column used was a Waters Acquity UPLC BEH C18 Column (1.7 μm, 2.1 × 50 mm), maintained at 30°C. 10 μL of the sample were injected onto the column and eluted at a flow rate of 0.4 ml/min. The gradient consisted of 45% mobile phase A (5 mM ammonium acetate in water) and 55% mobile phase B (methanol) for 2 minutes. This was succeeded by a linear gradient to 95% B from 2 to 2.5 minutes, holding at 95% B for 0.7 minutes, returning to 45% A, and finally re-equilibrating the column at initial conditions for 1 minute. The total run time was 4.7 minutes.

In positive ion mode, mass transitions of m/z 203 → 161 (KIC), 206 → 161 (KIC-d3), 189 → 174 (KIV), 194 → 178 (KIV-^5^C_13_), 203 → 174 (KMV), and 211 → 177 (KMV-d8) were monitored. Endogenous keto acids were quantified using calibrators prepared by spiking dialyzed fetal bovine serum with authentic keto acids (Sigma-Aldrich).

##### Flow injection MS/MS analysis of acyl CoAs

Duke University conducted targeted profiling of acyl CoAs. Here, 500 μL of tissue homogenate, prepared at a concentration of 50 mg/ml in isopropanol/0.1 M KH2PO4 (1:1), underwent extraction with an equal volume of acetonitrile. The resulting mixture was then centrifuged at 14,000 x g for 10 minutes, following a previously established procedure.^158,159^

The supernatants were acidified with 0.25 ml of glacial acetic acid, and the acyl CoAs were subjected to additional purification through solid-phase extraction (SPE) using 2-(2-pyridyl) ethyl functionalized silica gel (Sigma-Aldrich).^160^ Prior to use, the SPE columns were conditioned with 1 ml of acetonitrile/isopropanol/water/glacial acetic acid (9/3/4/4 : v/v/v/v). After application and flow-through of the supernatant, the SPE columns underwent a washing step with 2 ml of acetonitrile/isopropanol/water/glacial acetic acid (9/3/4/4 : v/v/v/v). Acyl CoAs elution was achieved with 2 ml of methanol/250 mM ammonium formate (4/1 : v/v), followed by analysis using flow injection MS/MS in positive ion mode on a Xevo TQ-S triple quadrupole mass spectrometer (Waters). The mobile phase used was methanol/water (80/20, v/v) with 30 mM ammonium hydroxide. Spectra were acquired in the multichannel acquisition mode, monitoring the neutral loss of 507 amu (phosphoadenosine diphosphate) and scanning from m/z 750 to 1100. As an internal standard, heptadecanoyl CoA was employed.

Quantification of endogenous acyl CoAs was carried out using calibrators created by spiking tissue homogenates with authentic Acyl CoAs (Sigma-Aldrich) that covered saturated acyl chain lengths from C0 to C18. Empirical corrections for heavy isotope effects, particularly ^13^C, to the adjacent m+2 spectral peaks in a specific chain length cluster were made by referring to the observed spectra for the analytical standards.

##### LC-MS/MS analysis of nucleotides

Duke University conducted targeted profiling of nucleotide metabolites. 300 μL of tissue homogenates, prepared at a concentration of 50 mg/ml in 70% methanol, underwent spiking with nine internal standards: ^13^C^10,15^N^5^-adenosine monophosphate, ^13^C^10,15^N^5^-guanosine monophosphate, ^13^C^10,15^N^2^-uridine monophosphate, ^13^C^9,15^N^3^-cytidine monophosphate, ^13^C^10^-guanosine triphosphate, ^13^C^10^-uridine triphosphate, ^13^C^9^-cytidine triphosphate, ^13^C^10^-adenosine triphosphate, and nicotinamide-1, N^6^-ethenoadenine dinucleotide (eNAD) (all from Sigma-Aldrich).

Nucleotides were extracted using an equal volume of hexane, following the procedures previously outlined.^161,162^ After vortexing and centrifugation at 14,000 x g for 5 minutes, the bottom layer was subjected to another centrifugation. Chromatographic separation and MS analysis of the supernatants were performed using an Acquity UPLC system (Waters) coupled to a Xevo TQ-XS quadrupole mass spectrometer (Waters).

The analytical column employed was a Chromolith FastGradient RP-18e 50-2mm column (EMD Millipore, Billerica, MA, USA), that was maintained at 40°C. The injection volume was 2 μL. Nucleotides were separated using a mobile phase A consisting of 95% water, 5% methanol, and 5 mM dimethylhexylamine adjusted to pH 7.5 with acetic acid. Mobile phase B comprised 20% water, 80% methanol, and 10 mM dimethylhexylamine. The flow rate was set to 0.3 ml/min. The 22-minute gradient (t=0, %B=0; t=1.2, %B=0; t=22, %B=40) was followed by a 3-minute wash and 7-minute equilibration. Nucleotides were detected in the negative ion multiple reaction monitoring (MRM) mode based on characteristic fragmentation reactions. Endogenous nucleotides were quantified using calibrators prepared by spiked-in authentic nucleotides obtained from Sigma-Aldrich.

##### LC-MS/MS analysis of amino metabolites

The Mayo Clinic conducted LC-MS-based targeted profiling of amino acids and amino metabolites, as previously outlined.^163,164^ In summary, either 20 ml of plasma samples or 5 mg of tissue homogenates were supplemented with an internal standard solution comprising isotopically labeled amino acids (U-^13^C^4^ L-aspartic acid, U-^13^C_3_ alanine, U-^13^C_4_ L-threonine, U-^13^C L-proline, U-^13^C_6_ tyrosine, U-^13^C_5_ valine, U-^13^C_6_ leucine, U-^13^C_6_ phenylalanine, U-^13^C_3_ serine, U-^13^C_5_ glutamine, U-^13^C_2_ glycine, U-^13^C_5_ glutamate, U-^13^C_6_,^15^N_2_ lysine, U-^13^C_5_,^15^N methionine, 1,1U-^13^C_2_ homocysteine, U-^13^C_6_ arginine, U-^13^C_5_ ornithine, ^13^C_4_ asparagine, ^13^C_2_ ethanolamine, D_3_ sarcosine, D_6_ 4-aminobutyric acid).

The supernatant was promptly derivatized using 6-aminoquinolyl-N-hydroxysuccinimidyl carbamate with a MassTrak kit (Waters). A 10-point calibration standard curve underwent a similar derivatization procedure after the addition of internal standards. Both the derivatized standards and samples were subjected to analysis using a Quantum Ultra triple quadrupole mass spectrometer (ThermoFischer) coupled with an Acquity liquid chromatography system (Waters). Data acquisition utilized selected ion monitoring (SRM) in positive ion mode. The concentrations of 42 analytes in each unknown sample were calculated against their respective calibration curves.

##### GC-MS analysis of TCA metabolites

The Mayo Clinic conducted GC-MS-targeted profiling of tricarboxylic acid (TCA) metabolites, as previously detailed,^165,166^ with some modifications. In summary, 5 mg of tissue were homogenized in 1X PBS using an Omni bead ruptor (Omni International, Kennesaw, GA), followed by the addition of 20 μL of an internal solution containing U-^13^C labeled analytes (^13^C_3_ sodium lactate, ^13^C_4_ succinic acid, ^13^C_4_ fumaric acid, ^13^C_4_ alpha-ketoglutaric acid, ^13^C_4_ malic acid, ^13^C_4_ aspartic acid, ^13^C_5_ 2-hydroxyglutaric acid, ^13^C_5_ glutamic acid, ^13^C_6_ citric acid, ^13^C_2_,^15^N glycine, ^13^C_2_ sodium pyruvate). For plasma, 50 μL were used.

The proteins were precipitated out using 300 μL of a mixture of chilled methanol and acetonitrile solution. After drying the supernatant in the speedvac, the sample was derivatized first with ethoxime and then with MtBSTFA + 1% tBDMCS (N-Methyl-N-(t-Butyldimethylsilyl)-Trifluoroacetamide + 1% t-Butyldimethylchlorosilane) and then analyzed on an Agilent 5977B GC/MS (Santa Clara, California) under single ion monitoring conditions using electron ionization. Concentrations of lactic acid (m/z 261.2), fumaric acid (m/z 287.1), succinic acid (m/z 289.1), ketoglutaric acid (m/z 360.2), malic acid (m/z 419.3), aspartic acid (m/z 418.2), 2-hydroxyglutaratic acid (m/z 433.2), cis-aconitic acid (m/z 459.3), citric acid (m/z 591.4), isocitric acid (m/z 591.4), and glutamic acid (m/z 432.4) were measured against 7-point calibration curves that underwent the same derivatization procedure.

##### Targeted lipidomics of low-level lipids

Lipid targeted profiling was conducted at Emory University following established methodologies as previously described.^167,168^ In summary, 20 mg of powdered tissue samples were homogenized in 100 μL PBS using Bead Ruptor (Omni International, Kennesaw, GA). Homogenized tissue samples (or plasma samples) were diluted with 300 μL 20% methanol and spiked with a 1% BHT solution to a final BHT concentration of 0.1% and pH of 3.0 by acetic acid addition. After centrifugation (10 minutes, 14000 rpm), the supernatants were transferred to 96-well plates for further extraction.

The supernatants were loaded onto Isolute C18 solid-phase extraction (SPE) columns that had been conditioned with 1000 μL ethyl acetate and 1000 μL 5% methanol. The SPE columns were washed with 800 μL water and 800 μL hexane, and the oxylipins were then eluted with 400 μL methyl formate. The SPE process was automated using a Biotage Extrahera (Uppsala, Sweden). The eluate was dried with nitrogen and reconstituted with 200 μL acetonitrile before LC-MS analysis. Sample blanks, pooled extract samples used as quality controls (QC), and consortium reference samples were prepared for analysis using the same methods. All external standards were purchased from Cayman Chemical (Ann Arbor, Michigan) at a final concentration in the range 0.01-20 μg/ml and consisted of: prostaglandin E2 ethanolamide (Catalog No. 100007212), oleoyl ethanolamide (Catalog No. 90265), palmitoyl ethanolamide (Catalog No. 10965), arachidonoyl ethanolamide (Catalog No. 1007270), docosahexaenoyl ethanolamide (Catalog No. 10007534), linoleoyl ethanolamide (Catalog No. 90155), stearoyl ethanolamide (Catalog No. 90245), oxy-arachidonoyl ethanolamide (Catalog No. 10008642), 2-arachidonyl glycerol (Catalog No. 62160), docosatetraenoyl ethanolamide (Catalog No. 90215), α-linolenoyl ethanolamide (Catalog No. 902150), oleamide (Catalog No. 90375), dihomo-γ-linolenoyl ethanolamide (Catalog No. 09235), decosanoyl ethanolamide (Catalog No. 10005823), 9,10 DiHOME (Catalog No. 53400), prostaglandin E2-1-glyceryl ester (Catalog No. 14010), 20-HETE (Catalog No. 10007269), 9-HETE (Catalog No. 34400), 14,15 DiHET (Catalog No. 10007267), 5(S)-HETE (Catalog No. 34210), 12(R)-HETE (Catalog No. 10007247), 11(12)-DiHET (Catalog No. 10007266), 5,6-DiHET (Catalog No. 10007264), thromboxane B2 (Catalog No. 10007237), 12(13)-EpOME (Catalog No. 52450), 13 HODE (Catalog No. 38600), prostaglandin F2α (Catalog No. 10007221), 14(15)-EET (Catalog No. 10007263), 8(9)-EET (Catalog No. 10007261), 11(12)-EET (Catalog No. 10007262), leukotriene B4 (Catalog No. 20110), 8(9)-DiHET (Catalog No. 10007265), 13-OxoODE (Catalog No. 38620), 13(S)-HpODE (Catalog No. 48610), 9(S)-HpODE (Catalog No. 48410), 9(S)-HODE (Catalog No. 38410), resolvin D3 (Catalog No. 13834), resolvin E1 (Catalog No. 10007848), resolvin D1 (Catalog No. 10012554), resolvin D2 (Catalog No. 10007279), 9(S)HOTrE (Catalog No. 39420), 13(S)HOTrE (Catalog No. 39620), 8-iso Progstaglandin F2α (Catalog No. 25903), maresin 1 (Catalog No. 10878), maresin 2 (Catalog No. 16369).

LC-MS data were acquired using an Agilent 1290 Infinity II chromatograph from Agilent (Santa Clara, CA), equipped with a ThermoFisher Scientific AccucoreTM C18 column (100 mm × 4.6, 2.6 µm particle size), and coupled to Agilent 6495 mass spectrometer for polarity switch multiple reaction monitoring (MRM) scan. The mobile phases consisted of water with 10 mM ammonium acetate (mobile phase A) and acetonitrile with 10 mM ammonium acetate (mobile phase B). The chromatographic gradient program was: 0.5 minutes with 95% A; 1 minute to 2 minutes with 65% A; 2.1 minutes to 5.0 minutes with 45% A; 7 minutes to 20 minutes with 25% A; and 21.1 minutes until 25 minutes with 95% A. The flow rate was set at 0.40 ml/min. The column temperature was maintained at 35°C, and the injection volume was 6 μL. For MS analysis, the MRM analysis is operated at gas temperature of 290 °C, gas flow of 14 L/min, Nebulizer of 20 psi, sheath gas temperature of 300 °C, sheath gas flow of 11 L /min, capillary of 3000 V for both positive and negative ion mode, nozzle voltage of 1500 V for both positive and negative ion mode, high pressure RF of iFunnel parameters of 150V for both positive and negative ion mode, and low pressure of RF of iFunnel parameters of 60 V for both positive and negative ion mode. Skyline (version 25.1.0.142)^169^ was utilized to process raw LC-MS data. Standard curves were constructed for each oxylipin/ethanolamide and scrutinized to ensure that all concentration points fell within the linear portion of the curve with an R-squared value not less than 0.9. Additionally, features exhibiting a high coefficient of variation (CV) among the quality control (QC) samples were eliminated from the dataset. Pearson correlation among the QCs for each tissue type was computed using the Hmisc R library, and the figures documented in the QC report were generated and visualized with the corrplot R library.^170,171^

##### Hydrophilic interaction LC-MS metabolomics

The untargeted analysis of polar metabolites in the positive ionization mode was conducted at the Broad Institute of MIT and Harvard. The LC-MS system consisted of a Shimadzu Nexera X2 UHPLC (Shimadzu Corp., Kyoto, Japan) coupled to a Q-Exactive hybrid quadrupole Orbitrap mass spectrometer (Thermo Fisher Scientific). Plasma samples (10 μL) were extracted using 90 μL of 74.9/24.9/0.2 v/v/v acetonitrile/methanol/formic acid containing valine-d8 and phenylalanine-d8 internal standards. Following centrifugation, the supernatants were directly injected onto a 150 x 2 mm, 3 µm Atlantis HILIC column (Waters). Tissue (10 mg) homogenization was performed at 4°C using a TissueLyser II (QAIGEN) bead mill set to two 2 minute intervals at 20 Hz in 300 μL of 10/67.4/22.4/0.018 v/v/v/v water/acetonitrile/methanol/formic acid valine-d8 and phenylalanine-d8 internal standards. The column was eluted isocratically at a flow rate of 250 μL/minute with 5% mobile phase A (10 mM ammonium formate and 0.1% formic acid in water) for 0.5 minute, followed by a linear gradient to 40% mobile phase B (acetonitrile with 0.1% formic acid) over 10 minutes, then held at 40% B for 4.5 minutes. MS analyses utilized electrospray ionization in the positive ion mode, employing full scan analysis over 70-800 m/z at 70,000 resolution and 3 Hz data acquisition rate. Various MS settings, including sheath gas, auxiliary gas, spray voltage, and others, were specified for optimal performance. Data quality assurance was performed by confirming LC-MS system performance with a mixture of >140 well-characterized synthetic reference compounds, daily evaluation of internal standard signals, and the analysis of four pairs of pooled extract samples per sample type inserted in the analysis queue at regular intervals. One sample from each pair was used to correct for instrument drift using “nearest neighbor” scaling while the second reference sample served as a passive QC for determination of the analytical coefficient of variation of every identified metabolite and unknown. Raw data processing involved the use of TraceFinder software (v3.3, Thermo Fisher Scientific) for targeted peak integration and manual review, as well as Progenesis QI (v3.0, Nonlinear Dynamics, Waters) for peak detection and integration of both identified and unknown metabolites. Metabolite identities were confirmed using authentic reference standards.

##### Reversed Phase-High Performance & Ion-pairing LC metabolomics

Reversed-phase and ion pairing LC-MS profiling of polar metabolites was conducted at the University of Michigan. LC-MS grade solvents and mobile phase additives were procured from Sigma-Aldrich, while chemical standards were obtained from either Sigma-Aldrich or Cambridge Isotope Labs. Plasma (50 μL aliquot) was extracted by addition of 200 μL of 1:1:1 v:v methanol:acetonitrile:acetone containing the following internal standards diluted from stock solutions to yield the specified concentrations: D_4_-succinic acid, 12.5 µM; :D_3_-malic acid,12.5 µM; D_5_-glutamic acid,12.5 µM; D_10_-leucine,12.5 µM; D_5_-tryptophan,12.5 µM;D_5_-phenylalanine,12.5 µM; D_3_-caffeine,12.5 µM; D_8_-lysine,12.5 µM; D_4_-chenodeoxycholic acid,12.5 µM; phosphatidylcholine(17:0/17:0),2.5 µM; phosphatidylcholine(19:0/19:0),2.5 µM; D_31_-palmitic acid,12.5 µM; D_35_-stearic acid,12.5 µM; gibberellic acid, 2.5 µM; epibrassinolide, 2.5 µM. The extraction solvent also contained a 1:400 dilution of Cambridge Isotope carnitine/acylcarnitine mix NSK-B, resulting in the following concentrations of carnitine/acylcarnitine internal standards: D_9_-L-carnitine, 380 nM; D_3_-L-acetylcarnitine, 95 nM; D_3_-L-propionylcarnitine, 19 nM; D_3_-L-butyrylcarnitine, 19 nM; D_9_-L-isovalerylcarnitine, 19 nM; D_3_-L-octanoylcarnitine, 19 nM; D_9_-L-myristoylcarnitine, 19 nM; D_3_-L-palmitoylcarnitine, 38 nM. After addition of solvent, samples were vortexed, incubated on ice for 10 minutes, and then centrifuged at 15,000 x g for 5 minutes. 150 μL supernatant were retrieved from the pellet, transferred to a glass autosampler vial with a 250 uL footed insert, and dried under a constant stream of nitrogen gas at ambient temperature. A QC sample was created by pooling residual supernatant from multiple samples. This QC sample underwent the same drying and reconstitution process as described for individual samples. Dried supernatants were stored at −80 C until ready for instrumental analysis. On the day of analysis, dried samples were reconstituted in 37.5 μL of 8:2 v:v water:methanol, vortexed thoroughly, and submitted for LC-MS.

Frozen tissue samples (skeletal muscle and adipose) were rapidly weighed into pre-tared, pre-chilled Eppendorf tubes and extracted in 1:1:1:1 v:v methanol:acetonitrile:acetone:water containing the following internal standards diluted from stock solutions to yield the specified concentrations: ^13^C_3_-lactic acid, 12.5 µM; ^13^C_5_-oxoglutaric acid, 125 nM;^13^C_5_-citric acid,1.25 µM; ^13^C_4_-succinic acid,125 nM; ^13^C_4_-malic acid, 125 nM; U-^13^C amino acid mix (Cambridge Isotope CLM-1548-1), 2.5 µg/mL; ^13^C_5_-glutamine, 6.25 µM;^15^N_2_-asparagine, 1.25 µM;^15^N_2_-tryptophan, 1.25 µM; ^13^C_6_-glucose, 62.5 µM; D_4_-thymine, 1 µM; ^15^N-anthranillic acid, 1 µM; gibberellic acid,1 µM; epibrassinolide, 1 µM. The extraction solvent also contained a 1:400 dilution of Cambridge Isotope carnitine/acylcarnitine mix NSK-B, resulting in the following concentrations of carnitine/acylcarnitine internal standards: D_9_-L-carnitine, 380 nM; D_3_-L-acetylcarnitine, 95 nM; D_3_-L-propionylcarnitine, 19 nM; D_3_-L-butyrylcarnitine, 19 nM; D_9_-L-isovalerylcarnitine, 19 nM; D_3_-L-octanoylcarnitine, 19 nM; D_9_-L-myristoylcarnitine, 19 nM; D_3_-L-palmitoylcarnitine, 38 nM. Sample extraction was performed by adding chilled extraction solvent to tissue sample at a ratio of 1 ml solvent to 50 mg wet tissue mass. Immediately after solvent addition, the sample was homogenized using a Branson 450 probe sonicator. Subsequently, the tubes were mixed several times by inversion and then incubated on ice for 10 minutes. Following incubation, the samples were centrifuged and 300 μL of the supernatant was carefully transferred to two autosampler vials with flat-bottom inserts and dried under a constant stream of nitrogen gas. A QC sample was created by pooling residual supernatants from multiple samples. This QC sample underwent the same drying and reconstitution process as described for individual samples. Dried supernatants were stored at −80 C until ready for instrumental analysis. On the day of analysis, samples were reconstituted in 60 μL of 8:2 v:v water:methanol and then submitted to LC-MS.

Reversed phase LC-MS samples were analyzed on an Agilent 1290 Infinity II / 6545 qTOF MS system with a JetStream electrospray ionization (ESI) source (Agilent Technologies, Santa Clara, California) using a Waters Acquity HSS T3 column, 1.8 µm 2.1 x 100 mm equipped with a matched Vanguard precolumn (Waters Corporation). Mobile phase A was 100% water with 0.1% formic acid and mobile phase B was 100% methanol with 0.025% formic acid. The gradient was as follows: Linear ramp from 0% to 100% B from 0-10 minutes, hold 100% B until 17 minutes, linear return to 0% B from 17 to 17.1 minutes, hold 0% B until 20 minutes. The flow rate was 0.45 ml/min, the column temperature was 55°C, and the injection volume was 5 μL. Each sample was analyzed twice, once in positive and once in negative ion mode MS, scan rate 2 spectra/sec, mass range 50-1200 m/z. Source parameters were: drying gas temperature 350°C, drying gas flow rate 10 L/min, nebulizer pressure 30 psig, sheath gas temperature 350°C and flow 11 L/minute, capillary voltage 3500 V, internal reference mass correction enabled. A QC sample run was performed at minimum every tenth injection.

Ion-pair LC-MS samples were analyzed on an identically-configured LC-MS system using an Agilent Zorbax Extend C18 1.8 µm RRHD column, 2.1 x 150 mm ID, equipped with a matched guard column. Mobile phase A was 97% water, 3% methanol. Mobile phase B was 100% methanol. Both mobile phases contained 15 mM tributylamine and 10 mM acetic acid. Mobile phase C was 100% acetonitrile. Elution was carried out using a linear gradient followed by a multi-step column wash including automated (valve-controlled) backflushing, detailed as follows: 0-2min, 0%B; 2-11 min, linear ramp from 0-99%B; 12-16 min, 99%B, 16-17.5min, 99-0%B. At 17.55 minutes, the 10-port valve was switched to reverse flow (back-flush) through the column. From 17.55-20.45 minutes the solvent was ramped from 99%B to 99%C. From 20.45-20.95 minutes the flow rate was ramped up to 0.8 mL/min, which was held until 22.45 min, then ramped down to 0.6mL/minute by 22.65 minutes. From 22.65-23.45 minutes the solvent was ramped from 99% to 0% C while flow was simultaneously ramped down from 0.6-0.4mL/min. From 23.45 to 29.35 minutes the flow was ramped from 0.4 to 0.25mL/min; the 10-port valve was returned to restore forward flow through the column at 25 min.. Column temperature was 35°C and the injection volume was 5 μL. MS acquisition was performed in negative ion mode, scan rate 2 spectra/sec, mass range 50-1200 m/z. Source parameters were: drying gas temperature 250°C, drying gas flow rate 13 L/min, nebulizer pressure 35 psig, sheath gas temp 325°C and flow 12 L/min, capillary voltage 3500V, internal reference mass correction enabled. A QC sample run was performed at minimum every tenth injection.

Iterative MS/MS data was acquired for both reverse phase and ion pairing methods using the pooled sample material to enable compound identification. Eight repeated LC-MS/MS runs of the QC sample were performed at three different collision energies (10, 20, and 40) with iterative MS/MS acquisition enabled. The software excluded precursor ions from MS/MS acquisition within 0.5 minute of their MS/MS acquisition time in prior runs, resulting in deeper MS/MS coverage of lower-abundance precursor ions.

Feature detection and alignment was performed utilizing a hybrid targeted/untargeted approach. Targeted compound detection and relative quantitation was performed by automatic integration followed by manual inspection and correction using Profinder v8.0 (Agilent Technologies, Santa Clara, CA.) Non-targeted feature detection was performed using custom scripts that automate operation of the “find by molecular feature” workflow of the Agilent Masshunter Qualitative Analysis (v7) software package. Feature alignment and recursive feature detection were performed using Agilent Mass Profiler Pro (v8.0) and Masshunter Qualitative Analysis (“find by formula” workflow), yielding an aligned table including m/z, RT, and peak areas for all features.

<u>Data Cleaning and Degeneracy Removal:</u> The untargeted features and named metabolites were merged to generate a combined feature table. Features missing from over 50% of all samples in a batch or over 30% of QC samples were then removed prior to downstream normalization procedure. Next, the software package Binner was utilized to remove redundancy and degeneracy in the data.^172^ Briefly, Binner first performs RT-based binning, followed by clustering of features by Pearson’s correlation coefficient, and then assigns annotations for isotopes, adducts or in-source fragments by searching for known mass differences between highly correlated features.

<u>Normalization and Quality Control:</u> Data were then normalized using a Systematic Error Removal Using Random Forest (SERRF) approach,^173^ which helps correct for drift in peak intensity over the batch using data from the QC sample runs. When necessary to correct for residual drift, peak area normalization to closest-matching internal standard was also applied to selected compounds. Both SERRF correction and internal standard normalization were implemented in R. Parameters were set to minimize batch effects and other observable drifts, as visualized using principal component analysis score plots of the full dataset. Normalization performance was also validated by examining relative standard deviation values for additional QC samples not included in the drift correction calculations.

<u>Compound identification:</u> Metabolites from the targeted analysis workflow were identified with high confidence (MSI level 1)^174^ by matching retention time (+/- 0.1 minute), mass (+/- 10 ppm) and isotope profile (peak height and spacing) to authentic standards. MS/MS data corresponding to unidentified features of interest from the untargeted analysis were searched against a spectral library (NIST 2020 MS/MS spectral database or other public spectral databases) to generate putative identifications (MSI level 2) or compound-class level annotations (MSI level 3) as described previously.^175^

##### LC-MS/MS untargeted lipidomics

<u>Sample preparation:</u> Non-targeted lipid analysis was conducted at the Georgia Institute of Technology. Powdered tissue samples (10 mg) were extracted in 400 μL isopropanol containing stable isotope-labeled internal standards (IS) with bead homogenization with 2mm zirconium oxide beads (Next Advance) using a TissueLyser II (10min, 30 Hz). Samples were then centrifuged (5 min, 21,100xg), and supernatants were transferred to autosampler vials. Plasma samples (25 μL) were extracted by mixing with 75 μL isopropanol containing the IS mix followed by centrifugation. Sample blanks, pooled extract samples used as quality controls (QC), and consortium reference samples, were prepared for analysis using the same methods. The IS mix consisted of PC (15:0-18:1(d7)), Catalog No. 791637; PE (15:0-18:1(d7)), Catalog No. 791638; PS (15:0-18:1(d7)), Catalog No. 791639; PG(15:0-18:1(d7)), Catalog No. 791640; PI(15:0-18:1(d7)), Catalog No. 791641; LPC(18:1(d7)), Catalog No. 791643; LPE(18:1(d7)); Catalog No. 791644; Chol Ester (18:1(d7)), Catalog No. 700185; DG(15:0-18:1(d7)), Catalog No. 791647; TG(15:0-18:1(d7)-15:0), Catalog No. 791648; SM(18:1(d9)), Catalog No. 791649; Cholesterol (d7), Catalog No. 700041. All internal standards were purchased from Avanti Polar Lipids (Alabaster, Alabama) and added to the extraction solvent at a final concentration in the 0.1-8 μg/ml range.

<u>Data collection:</u> Lipid LC-MS data were acquired using a Vanquish (ThermoFisher Scientific) chromatograph fitted with a ThermoFisher Scientific AccucoreTM C30 column (2.1 × 150 mm, 2.6 µm particle size), coupled to a high-resolution accurate mass Q-Exactive HF Orbitrap mass spectrometer (ThermoFisher Scientific) for both positive and negative ionization modes. For positive mode analysis, the mobile phases were 40:60 water:acetonitrile with 10 mM ammonium formate and 0.1% formic acid (mobile phase A), and 10:90 acetonitrile:isopropyl alcohol, with 10 mM ammonium formate and 0.1% formic acid (mobile phase B). For negative mode analysis, the mobile phases were 40:60 water:acetonitrile with 10 mM ammonium acetate (mobile phase A), and 10:90 acetonitrile:isopropyl alcohol, with 10 mM ammonium acetate (mobile phase B). The chromatographic method used for both ionization modes was the following gradient program: 0 minutes 80% A; 1 minute 40% A; 5 minutes 30% A; 5.5 minutes 15% A; 8 minutes 10% A; held 8.2 minutes to 10.5 minutes 0% A; 10.7 minutes 80% A; and held until 12 minutes. The flow rate was set at 0.40 ml/min. The column temperature was set to 50°C, and the injection volume was 2 μL.

For analysis of the organic phase the electrospray ionization source was operated at a vaporizer temperature of 425°C, a spray voltage of 3.0 kV for positive ionization mode and 2.8 kV for negative ionization mode, sheath, auxiliary, and sweep gas flows of 60, 18, and 4 (arbitrary units), respectively, and capillary temperature of 275°C. The instrument acquired full MS data with 240,000 resolution over the 150-2000 m/z range. LC-MS/MS experiments were acquired using a DDA strategy. MS2 spectra were collected with a resolution of 120,000 and the dd-MS2 were collected at a resolution of 30,000 and an isolation window of 0.4 m/z with a loop count of top 7. Stepped normalized collision energies of 10%, 30%, and 50% fragmented selected precursors in the collision cell. Dynamic exclusion was set at 7 seconds and ions with charges greater than 2 were omitted.

<u>Data processing:</u> Data processing steps included peak detection, spectral alignment, grouping of isotopic peaks and adduct ions, drift correction, and gap filling. Compound Discoverer V3.3 (ThermoFisher Scientific) was used to process the raw LC-MS data. Drift correction was performed on each individual feature, where a Systematic Error Removal using Random Forest (SERRF) method builds a model using the pooled QC sample peak areas across the batch and was then used to correct the peak area for that specific feature in the samples. Detected features were filtered with background and QC filters. Features with abundance lower than 5x the background signal in the sample blanks and that were not present in at least 50% of the QC pooled injections with a coefficient of variance (CV) lower than 80% (not drift corrected) and 50% (drift corrected) were removed from the dataset. Lipid annotations were accomplished based on accurate mass and relative isotopic abundances (to assign elemental formula), retention time (to assign lipid class), and MS2 fragmentation pattern matching to local spectral databases built from curated experimental data. Lipid nomenclature followed that described previously.^176^

<u>Quality control procedures:</u> System suitability was assessed prior to the analysis of each batch. A performance baseline for a clean instrument was established before any experiments were conducted. The mass spectrometers were mass calibrated, mass accuracy and mass resolution were checked to be within manufacturer specifications, and signal-to-noise ratios for the suite of IS checked to be at least 75% of the clean baseline values. For LC-MS assays, an IS mix consisting of 12 standards was injected to establish baseline separation parameters for each new column. The performance of the LC gradient was assessed by inspection of the column back pressure trace, which had to be stable within an acceptable range (less than 30% change). Each IS mix component was visually evaluated for chromatographic peak shape, retention time (lower than 0.2 minute drift from baseline values) and FWHM lower than 125% of the baseline measurements. The CV of the average signal intensity and CV of the IS (<=15%) in pooled samples were also checked. These pooled QC samples were used to correct for instrument sensitivity drift over the various batches using a procedure similar to that described by the Human Serum Metabolome (HUSERMET) Consortium.^177^ To evaluate the quality of the data for the samples themselves, the IS signals across the batch were monitored, PCA modeling for all samples and features before and after drift correction was conducted, and Pearson correlations calculated between each sample and the median of the QC samples.

##### Metabolomics/Lipidomics Data filtering and normalization

The untargeted metabolomics datasets were categorized as either “named”, for chemical compounds confidently identified, or “unnamed”, for compounds with specific chemical properties but without a standard chemical name. While the preprocessing steps were performed on the named and unnamed portions together, only the named portions were utilized for differential analysis. For each dataset i.e. each assay for each tissue type, the following steps are performed:

- Average rows that have the same metabolite ID.
- Merge the “named” and “unnamed” subparts of the untargeted datasets.
- Convert negative and zero values to NAs.
- Remove features with > 20% missing values.
- Features with < 20% missing values are imputed either using K-Nearest Neighbor imputation (for datasets with > 12 features) or half-minimum imputation (for datasets with < 12 features).
- All data are log2-transformed. Note that for all log2 calculations, 1 is added to each value before log-normalization. This allows metabolite values that fall between 0 and 1 to have a positive log2 value.
- The untargeted data are median-MAD normalized if neither sample medians nor upper quartiles were significantly associated with sex or sex-stratified training group (Kruskal-Wallis p-value < 0.01).

Outlier detection was performed by examining the boxplot of each Principal Component and extending its whiskers to the predefined multiplier above and below the interquartile range (5x the IQR). Samples outside this range are flagged. All outliers were reviewed by Metabolomics CAS, and only confirmed technical outliers were removed.

##### Redundant Metabolite/Lipid Management

To address metabolites measured on multiple platforms (e.g. a metabolite measured on the HILIC positive and RP positive platforms), and metabolites with the same corresponding RefMet ID (e.g. alpha-Aminoadipic-acid, Aminoadipic acid both correspond to RefMet name ‘Aminoadipic acid’), we utilize the set of internal standards described above to make a decision on which platform’s measurement of a given feature to include in further analysis. For each tissue, based on whichever platform has the lowest coefficient of variation across all included reference standards for a given refmet id, that metabolite was chosen. The other platforms or metabolites, for this tissue, corresponding to this refmet id were removed from further downstream analysis. Data for all metabolites removed, including information about the coefficient of variation in the standards, normalized data, or differential analysis results, is available in the R Package(see below), but is not loaded by default.

### Statistical analysis

#### Differential analysis

To model the effects of both exercise modality and time, relative to non exercising control, each measured molecular feature was treated as an outcome in a linear mixed effects model accounting for fixed effects of exercise group (RE, EE, or CON), timepoint, as well as demographic and technical covariates (see covariate selection below for more info). Participant identification was treated as a random effect. For each molecular feature, a cell-means model is fit to estimate the mean of each exercise group-timepoint combination, and all hypothesis tests are done comparing the means of the fixed effect group-timepoint combinations. In order to model the effects of exercise against non-exercising control, a difference-in-changes model was used which compared the change from pre-exercise to a during or post-exercise timepoint in one of the two exercise groups to the same change in control. This effect is sometimes referred to as a “delta-delta” or “difference-in-differences” model.

##### Model selection via simulation

In order to determine the optimal statistical package/model for these data – given MoTrPAC’s unique sampling design (see ‘acute exercise intervention’ above) – a simulation study was conducted to determine the impact of various approaches to account for correlation in repeated measurements. The simulation evaluated type 1 error, power, and bias for multiple plausible modeling approaches, to identify those that would obtain higher power while maintaining nominal type 1 error rates. For a subset of molecular features in the datasets, mean, covariance, skew, and kurtosis were summarized over the relevant timepoints in the acute exercise bout within subgroups defined by sex and exercise type. Then, these summary values were used in combination with the “covsim” R package to generate non-normally distributed simulated data.^178^ Additionally, sample size and missingness patterns were aligned in the simulated data to match those of the observed data. Using a total of 5 million simulated instances where data were generated under the null hypothesis (i.e., no change in the mean values of the outcome over the exercise timepoints) or alternative hypothesis (i.e., the mean value of the outcome differs between at least two timepoints), type 1 error rate and power, respectively, were evaluated for seven analytic strategies: (1) pairwise t-tests between timepoints of interest, (2) ordinary least squares regression, (3) differential expression for repeated measures (i.e., dream) with random intercepts,^140^(4) linear mixed models with random intercepts, (5) mixed models for repeated measures using an unstructured covariance matrix, and (6) generalized linear models using generalized estimating equations (GEEs), and (7) a first degree autoregressive correlation structure. These analyses were replicated for adipose, blood, and muscle samples. The dream function obtained type 1 error rates 5.2, 5.4, and 5.2% for adipose, blood, and muscle analyses, respectively, with higher power than all other methods apart from generalized linear models with GEEs. Although generalized linear models with GEEs had higher power than dream, they also had higher type 1 error with 7.7%, 6.0%, and 6.5% type 1 error rates for adipose, blood, and muscle analyses, respectively. Therefore, due to stability of type 1 error and relatively high power compared to other approaches, the *dream* function from the variancePartition R package was selected for the primary analyses for every omic platform except Methyl-Cap (see methylation capture sequencing methods above for more details).

##### Covariate selection

Covariates were selected through a combination of a priori knowledge about factors influencing molecular levels as well through empiric screening. Factors were considered for model inclusion by correlating the principal components of each tissue-ome feature set to demographic and technical factors using variancePartition::canCorPairs.^140^ Visualizations and computational analysis that describe this process for the decisions for covariate selection can be found at: https://github.com/MoTrPAC/motrpac-human-presuspension-acute/tree/main/QC. Ultimately, the following fixed effect covariates were included in the models for every omic platform, except Methyl Cap (see methods above): group and timepoint in a cell means model, clinical site, age, sex, and BMI. ‘Participant id’ was included as a random effect in every model. Each ome then had ome-specific covariates selected as described in prior sections.

A targeted analysis was conducted to assess whether and how race, ethnicity, and genetic ancestry could be incorporated, given prior evidence that these factors can influence omic data.^179,180^ We evaluated multiple models that included patient-reported race and/or ethnicity, as well as principal components derived from whole-genome sequencing (WGS), to determine their impact on model fit using the BIC. However, no combinations of individual or joint inclusion of race, ethnicity, or WGS-derived principal components led to an improvement in average model BIC across all features for a platform.

The final set of covariates included in each model for each tissue and ome can be found in the R Package *MotrpacHumanPreSuspensionAnalysis*, and in Supplementary Table 2.

##### Sex-specific sensitivity analysis

To test whether the differential model was driven by the predominantly female cohort (72%), a sensitivity analysis estimated exercise effects for male and female participants within each group-timepoint combination. The analysis was limited to transcriptomics across all tissues and metabolomics in blood and muscle, the datasets with the largest sample sizes. The group-timepoint term from the overall analysis was replaced with a sex-group-timepoint term to estimate sex-specific effects. Because the control groups had smaller sample sizes, this comparison contrasted each exercise timepoint against group pre-exercise values rather than using a difference-in-differences model. The clinical site covariate was also omitted to limit the number of estimated parameters. Enrichment analysis was performed for the transcriptomic features that exhibited differential responses in the male vs female participants via ORA using the background of all transcripts measured in the tissue.

##### Cell type sensitivity analysis

To evaluate whether shifts in cellular composition confounded the differential expression results, a sensitivity analysis was performed using deconvolution-derived cell-type proportions in blood. Cellular composition was first estimated from whole-blood transcriptomes by deconvolving the bulk RNA-seq profiles with CIBERSORTx (cibersortx.stanford.edu) using the LM22 leukocyte signature matrix in ‘Batch Correct’ mode disabled. Because LM22 resolves 22 closely related leukocyte subsets whose marker profiles are highly correlated, the subsets were collapsed by summing their fractions into broader populations: T cells, B cells, NK cells, monocytes, macrophages, dendritic cells, neutrophils, and mast cells. The reliability of these estimates was then assessed by correlating CIBERSORTx proportions at pre-exercise with matched complete blood count values for the same participants, where available. Finally, the blood RNA-seq differential analysis was refit with the coarse cell-type fractions (T cells, B cells, NK cells, monocytes, and neutrophils) added as numeric covariates. Proportions were z-scored before inclusion, and the model otherwise matched the original differential analysis pipeline. Comparing the refit results against the original model indicated whether accounting for cellular composition altered the estimated exercise effects.

##### Missingness

Data missingness varied for multiple reasons including experimental design (i.e. temporal randomization which randomized some participants to have samples obtained at a subset of timepoints, see Supplementary Table 1 and assay limitations (metabolomics and proteomics can have missingness as described in their individual sections). For some omic sets, imputation to alleviate missingness could be performed (metabolomics) but for others was not, including proteomics. Analysis demonstrated proteomics missingness was approximately at random (data not shown).

Experimental samples were designed and processed with a minimum of 6 samples for each exercise group at every timepoint. After outlier removal, and due to expected and standard proteomics processing, some individual molecular features in proteomic platforms may have fewer observations due to missingness. Given the above issues affecting the ultimate sample size for effect estimation, for a feature to be included in differential analysis, a minimum number of 3 participants were required to have a paired pre-exercise sample for all groups and all during/post-exercise timepoints. Thus, all group comparisons (e.g. RE vs. CON at 3.5 hours post-exercise) required 3 participants with matched pre- and post-exercise samples in each group. Unpaired samples (e.g. a participant with only one time point measured) were kept in the analysis to inform group-timepoint means but do not count toward the minimum 3 paired n samples. This requirement was satisfied in all omes and tissues except a subset of MS-acquired proteomics and phosphoproteomics features in SKM and AT. Features not meeting this criterion in any group-timepoint set were excluded from differential analysis.

##### Specific omic-level statistical considerations

Models for all omes other than methylation were fit using ‘variancePartition::dream’.^140^ For all other omes the normalized values were used directly as input to the statistical model.

The full implementation of the statistical models for each of non-methylation datasets can be found in the R Package *MotrpacHumanPreSuspensionAnalysis*. Methylation data was processed separately, and the methods can be found in the methylation methods section.

##### Contrast types

The specific contrasts made in the statistical analysis include three categories of comparison: difference-in-differences relative to control, group-specific, and resistance vs endurance. All comparisons were structured using ‘variancePartition::makeContrastsDream’.^140^

The difference-in-differences model as described at the opening of this section represents the primary DA analysis, and any individual feature mentioned as statistically significantly changing due to exercise will be referring to the difference-in-differences results unless otherwise specifically stated.

The next category of contrasts is a group-specific comparison, which compares a given post-intervention timepoint to pre-exercise within a given group, without comparing to the control group. This contrast was implemented to more easily quantify effect sizes within each group independently.

Finally, the RE vs EE is very similar to the difference-in-differences model, but sets the RE group as a matched control to the EE group. In this comparison, the change in response to exercise at a given timepoint in one exercise group is directly compared to the corresponding change in the other exercise group. This contrast can directly test the hypothesis that the EE effect is different from the RE effect. There are no samples obtained during RE bouts, so those timepoints in blood were excluded from this comparison.

##### Contrast categorization

To characterize the response of individual molecular features to exercise, we assigned features to one of three mutually exclusive categories based on adjusted p-values from the difference-in-differences and group-specific contrasts at a given timepoint:

<u>Exercise effects</u>, in which a change within the exercise group remains significant after accounting for the control group. These features change from their respective exercise-group baseline, and that change differs from the control group at the same timepoint, ruling out non-exercise stimuli as the sole explanation. This requires the following: Exercise vs control (difference-in-differences) adjusted p < 0.05; Exercise vs pre-exercise (group-specific) adjusted p < 0.05; Control vs pre-exercise (group-specific) not considered

<u>Non-exercise induced changes</u>, in which features change to a similar degree in exercise and control groups. These changes reflect stimuli shared across groups, such as biopsy, fasting, or circadian rhythm. This category quantifies the value of the control arm: in a single-arm design, these features would be misattributed to exercise. Reporting them separately allows readers to assess which apparent exercise responses in uncontrolled studies may instead reflect protocol effects.This requires the following: Exercise vs control adjusted p > 0.05; Exercise vs pre-exercise adjusted p < 0.05; Control vs pre-exercise adjusted p < 0.05

<u>Control-driven exercise effects,</u> in which an exercise effect offsets a change occurring in the control group, such that the exercise response is detectable only relative to the control. This category captures features that would be missed in a single-arm design: the exercise group appears stable from baseline, but the lack of response reflects an exercise effect counteracting a biopsy, fasting, or circadian change seen in controls. This requires the following: Exercise vs control adjusted p < 0.05; Exercise vs pre-exercise adjusted p > 0.05; Control vs pre-exercise not considered

Features meeting none of these criteria were left uncategorized.

Within the exercise effects category, we further classified features by exercise modality. This classification distinguishes responses unique to one training mode from responses common to both, which informs whether a feature reflects a modality-specific stimulus or a general response to exercise:

<u>EE-specific</u>, in which the response is detectable after endurance but not resistance exercise, and the two responses differ from each other. This requires the following: EE vs control adjusted p < 0.05; RE vs control adjusted p > 0.05; EE vs RE adjusted p < 0.05

<u>RE-specific</u>, in which the response is detectable after resistance but not endurance exercise, and the two responses differ from each other. This requires the following: EE vs control adjusted p > 0.05; RE vs control adjusted p < 0.05; EE vs RE adjusted p < 0.05

<u>EE-RE shared</u>, in which the response is detectable after both modalities and does not differ between them. This requires the following: EE vs control adjusted p < 0.05; RE vs control adjusted p < 0.05; EE vs RE adjusted p > 0.05

We note that these categories depend on a fixed significance threshold, and features near the threshold may shift categories under alternative cutoffs; category assignments should therefore be interpreted as descriptive.

##### Significance thresholds

For each of the above contrasts, p-values were adjusted for multiple comparisons for each unique contrast-group-tissue-ome-timepoint combination separately using the Benjamini-Hochberg method to control False Discovery Rate (FDR).^181^ Features were considered significant at a FDR of 0.05 unless otherwise stated.

#### Human feature to gene mapping

The feature-to-gene map links each feature tested in differential analysis to a gene, using Ensembl version 105 (mapped to GENCODE 39)^182^ as the gene identifier source. Proteomics feature IDs (UniProt IDs) were mapped to gene symbols and Entrez IDs using UniProt’s mapping files.^183^ Epigenomics features were mapped to the nearest gene using the ChIPseeker::annotatePeak()^184,185^ function with Homo sapiens Ensembl release 105 gene annotations. Gene symbols, Entrez IDs, and Ensembl IDs were assigned to features using biomaRt version 2.58.2 (Bioconductor 3.18).^186–188^

For ATAC-seq and methyl capture features, relationship to gene and custom annotation provide gene proximity information and custom ChIPseeker annotations, respectively. Relationship to gene values indicate the distance of the feature from the closest gene, with 0 indicating overlap.

Custom annotation categories include Distal Intergenic, Promoter (<=1kb), Exon, Promoter (1-2kb), Downstream (<5kb), Upstream (<5kb), 5’ UTR, Intron, 3’ UTR, and Overlaps Gene.

Metabolite features were mapped to KEGG IDs using KEGGREST, RefMet REST API, or web scraping from the Metabolomics Workbench, built on August 4th, 2026.^189,190^

#### Association analysis between pre-exercise molecular abundance and clinical phenotypes

For core clinical phenotypes, the association between pre-exercise (resting) molecular abundance and phenotype measurements were determined using linear regression. Analyses were conducted separately for each tissue, assay, and clinical variable of interest. Clinical traits included in the analysis were: absolute VO_2_peak (L/min), chest press 1RM (lbs), leg press 1RM (lbs), leg extension 1RM (lbs), HOMA-IR, post-exercise lactate (P10M), and non-esterified fatty acids (NEFA).

One-repetition maximum strength was assessed for the leg press, chest press, and leg extension using the standard MoTrPAC protocol for resistance exercise participants. Following exercise familiarization and warm-up, resistance was progressively increased until the participant completed the heaviest load possible for one repetition. If a participant reached the maximum resistance available on the machine before a true 1RM could be established, the participant rested for 60 seconds and then completed as many repetitions as possible at the machine’s maximum load. For participants who completed more than one repetition, the Data Management, Analysis, and Quality Control Center calculated an estimated 1RM using the Brzycki equation: estimated 1RM = weight lifted ÷ [1.0278 − (0.0278 × number of repetitions)].^191^ Because the Brzycki equation was developed primarily for sets of fewer than 10 repetitions, records exceeding the 10 repetitions were considered invalid and not calculated. Because strength metrics are limited to only the resistance exercise cohort, the statistical power to detect associated pre-exercise molecular measurements is limited compared to non-strength metrics (n = 55-70 resistance participants vs up to 172 for metrics measured on all participants).

HOMA-IR was calculated from fasting insulin and glucose as:

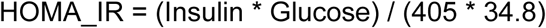

For NEFA, pre-exercise measurements were used for participants in any randomized group (Control, Resistance, Endurance). However, for lactate, lactate levels at 10 minutes post-exercise were used only for Resistance and Endurance participants, and controls were excluded from this clinical endpoint. Since lactate is the only endpoint measured post-exercise, and is bimodal depending on exercise group, the exercise modality was included as an additional covariate for this endpoint only.

For each feature, a linear regression model was fit (using the lm() function in the stats package in R)^192^ to test the association between pre-exercise molecular abundance and the clinical variable of interest. Models required a minimum of 5 residual degrees of freedom for stable estimation (i.e., n > p + 5, where n is the number of samples and p is the number of predictors). Because of inherent missingness in the global mass-spectrometry proteomics data in adipose and muscle, this criterion could not be met for all proteins, and some were therefore excluded from model fitting. The outcome variable was feature abundance, and the primary independent variable was the clinical trait, adjusted for age, sex, and BMI. Subsequently, the estimated regression coefficient, standard error, t-statistic, nominal p-value, and number of observations for each feature were extracted for each clinical trait. Multiple testing correction was performed within each tissue-ome-trait combination using the Benjamini-Hochberg false discovery rate method. For the outcomes “Lactate” and “NEFA,” these metabolites were measured using the blood metabolomics platform but were excluded as predictors when modeling their respective outcomes to avoid predicting each metabolite from itself.

Pre-exercise clinical trait association results were integrated with DA results with acute exercise by first restricting to features with an adjusted p-value < 0.05 for clinical association, and then classifying exercise-responsive features as described in ‘Contrast Categorizations’ above.

All analyses were performed in R version 4.4.1.

#### Enrichment analysis

##### Preparation of differential analysis results

The differential analysis results tables for each combination of tissue and ome were converted to matrices of z-scores with either gene symbols, RefMet metabolite/lipid IDs, or phosphorylation flanking sequences as rows and contrasts as columns. These matrices serve as input for the enrichment analyses. For the transcriptomics, TMT proteomics, and Olink proteomics results, transcripts and proteins were first mapped to gene symbols. To resolve cases where multiple features mapped to a single gene, only the most extreme z-score for each combination of tissue, contrast, and gene was retained. Metabolite and lipid identifiers were standardized using the Metabolomic Workbench Reference List of Metabolite Names (RefMet) database.^193^ For the CAMERA-PR enrichment of phosphoproteomics results, since peptides could be phosphorylated at multiple positions, phosphorylation sites were separated into single sites with identical row information (e.g., protein_S1;S2 becomes protein_S1 and protein_S2, both with the same data). For any combinations of contrast and phosphosite that were not uniquely defined, only the most extreme z-score (maximum absolute value) was selected for inclusion in the matrix. Singly phosphorylated sites were required to use the curated kinase–substrate relationship information available from PhosphoSitePlus (PSP),^194^ described below.

##### Gene set selection

Gene sets were obtained from the MitoCarta3.0 database, CellMarker 2.0 database,^126^ and the C2-CP (excluding KEGG_LEGACY) and C5 collections of the human Molecular Signatures Database (MSigDB; v2023.2.Hs)^126,195,196^ Metabolites and lipids were grouped according to chemical subclasses from the RefMet database.^193^ This includes subclasses such as “Acyl carnitines” and “Saturated fatty acids”.Flanking sequences for protein phosphorylation sites were grouped according to their known human protein kinases provided in the *Kinase_Substrate_Dataset* file from PhosphoSitePlus (PSP; v6.7.1.1; https://www.phosphosite.org/staticDownloads.action; last modified 2023-11-17).^194^ In addition to these kinase sets, sets from the directional Post-Translational Modification Signatures Database (PTMsigDB) were included for analysis.^197^

For each combination of tissue and ome, molecular signatures were filtered to only those genes, metabolites/lipids, or flanking sequences that appeared in the differential analysis results. After filtering, all molecular signatures were required to contain at least 5 features, with no restriction on the maximum size of sets. Additionally, gene sets were only kept if they retained at least 70% of their original genes to increase the likelihood that the genes that remain in a given set are accurately described by the set label.

##### Analysis of Molecular Signatures

Analysis of molecular signatures was carried out with the pre-ranked Correlation Adjusted MEan RAnk (CAMERA-PR) gene set test using the z-score matrices described in the “Preparation of differential analysis results” section to summarize the differential analysis results for each contrast at the level of molecular signatures.^197,198^ Z-scores were selected as the input statistics primarily to satisfy the normality assumption of CAMERA-PR, and the analysis was carried out with the *cameraPR.matrix* function from the TMSig R/Bioconductor package.^192,199^

##### Over-representation analysis (ORA)

Over-representation analysis (ORA) was conducted using the run_ORA function from the *MotrpacHumanPreSuspensionAnalysis* R package which uses the hypergeometric test for statistical significance. The input consisted of differentially expressed genes (adjusted p-value < 0.05) identified from one or more omic layers in a particular tissue. The background gene set included all genes detected in each of the ome(s) and tissue(s) under consideration.

##### Statistical significance thresholds

For both CAMERA-PR and ORA results, p-values were adjusted within each combination of tissue, ome, contrast, and broad molecular signature collection (MitoCarta3.0, CellMarker 2.0, C2, C5, RefMet, PSP, and PTMsigDB) using the Bejamini-Hochberg method. Gene sets and RefMet subclasses were declared significant if their adjusted p-values were less than 0.05. This threshold was raised to 0.1 for kinase sets.

##### Visualization methods

Bubble heatmaps were generated from the enrichment analysis results using the *enrichmap* function from the TMSig R/Bioconductor package.^199^

#### Clustering techniques

##### Fuzzy C-Means Clustering

The z-score matrices described in the Enrichment Analysis Methods were used as input for fuzzy c-means (FCM) clustering.^200,201^ For each tissue, the matrices for the different omes, excluding global proteomics and phosphoproteomics in adipose (as these were only measured at two timepoints) were stacked to form a single matrix with tissue-specific timepoints as columns and features being a row. For blood, the “during exercise” contrasts were also excluded. The rows of this matrix were then divided by their sample standard deviations, which were calculated after two columns of zeros were temporarily included to represent the pre-exercise timepoints; these zero columns were discarded before clustering. The Mfuzz *R* package was used to perform FCM clustering with the parameter *m* set to the value of *Mfuzz::mestimate* for each tissue.^202,203^ The minimum distance between centroids was used as the cluster validity index to determine the optimal number of clusters for each tissue. A scree/elbow plot of these distances for a range of cluster numbers (3-14) was generated for each tissue, and the final cluster numbers were determined through manual inspection: 13 for both adipose, and 12 for blood and muscle. For visualization purposes, hard clustering was performed by assigning each feature to the cluster with the highest membership probability; features with probabilities below 0.3 are not clustered or displayed. Cluster numbers are assigned arbitrarily.

##### Analysis of clusters

In order to uncover the biological relevance of each cluster, non-parametric CAMERA-PR as well as ORA was applied to the matrix of cluster membership probabilities separately for each tissue and ome, though CAMERA-PR results are available only though the R package.^198^ The same molecular signatures described in the “Enrichment Analysis” Methods were tested. The non-parametric version of CAMERA-PR is a modification of the two-sample Wilcoxon–Mann–Whitney rank sum test. As with the parametric version, it accounts for inter-molecular correlation to control the false positive rate. Upper-tailed tests were used to determine which molecular signatures were highly ranked according to their membership probabilities (i.e., which molecular signatures closely followed the cluster centroids).

#### PLIER methods

To delve into cross-tissue temporal trends in acute exercise response, we applied PLIER,^204^ which takes an expression matrix of features vs samples as input and outputs patterns of activity, in the form of latent variables (LVs), and connects these patterns with user-provided prior information, such as reactome or KEGG pathways, or structural classifications in the case of metabolomics data. We use the PLIER package R function *num.pc* to identify the number of significant principle components for the singular value decomposition (SVD) of the data and specify double this number as the desired LV count. If *num.pc* is computationally infeasible or produces a value > 50, we use a default of 100 LVs for a PLIER run, with rigorous testing of alternative LV depth to ensure coverage of the significant discoverable exercise patterns. Both the cross-tissue RNA analysis and the cross-tissue metabolomics analysis specified the default 100 LVs. PLIER works to identify a set of LVs that best account for all patterns identified in the dataset, optimizing individual LV trajectories over the samples and feature associations to minimize the difference between normalized input data and PLIER LV output, while also accounting for feature associations with prior knowledge in the form of pathway enrichments or metabolite classifications.

We ran PLIER separately on the cross-tissue RNA seq data and on the cross-tissue metabolomics data. For each ome separately studied, we generated a cross-tissue matrix of exercise response, selecting the subset of features that were present in all three tissues, then z-scoring the normalized data matrix for the given ome for each tissue. Following by-tissue z-scoring, the z-scored tissue matrices are concatenated into one large matrix with rows corresponding to the consensus features across all tissues, and columns the cumulative samples from each tissue. Significance of an LV to differentiate resistance or endurance exercise from control at a given timepoint relative to pre-exercise is determined by comparing the differences in LV value post-exercise vs pre-exercise for each subject at that timepoint separately between resistance and control and endurance and control within each tissue and adjusting for multiple hypotheses using the *compare_means* R function.^205^

#### Transcription Factor Motif Enrichment

Transcription factor motif enrichment analysis was performed on sets of DEGs for each tissue, timepoint and exercise modality combination. DEGs for motif enrichment analysis were selected for each tissue by satisfying an unadjusted p-value threshold of 0.05. Motif enrichment analysis was carried out by findMotifs.pl (HOMER v4.11.1)^67^ on a set of 472 TF motifs provided by the HOMER package. Enrichment was performed on a subset of the proximal promoter region of each DEG, ranging from 300 bp upstream to 50 bp downstream of the transcription start site (TSS), and normalized for GC%. All HOMER runs used the same background set of all gene promoters. The search lengths of the motifs were 8, 10, and 12 bp. Downstream analysis of HOMER results were conducted using the knownResults.txt output file of enrichments of the 472 inputted TF motifs using Benjamini-Hochberg FDR-corrected p-values for significance of TF enrichment.

#### Spatiotemporal Clustering and Inference of Omics Networks

Spatiotemporal Clustering and Inference of Omics Networks (SC-ION, https://github.com/nmclark2/SCION)^77^ was applied to the dataset to infer unsupervised multi-omics networks of exercise response. The exact parameters used to generate results for the figures presented in the manuscript are found in the figures folder in the motrpac-human-presuspension-acute repository.

##### Selecting regulator and target data

SC-ION takes as input two data matrices where one matrix is assigned as the “regulator” matrix and the other is assigned as the “target” matrix. Two different schemes for regulator/target matrices were considered. In the first scheme (Muscle-only network), phosphosites on transcription factors (TFs) in muscle were used as the regulators, and DA transcripts in the muscle were used as the targets. In the second scheme (Blood-to-muscle network), metabolites in blood were considered regulators, and DA transcripts in muscle were again the targets. In both schemes, EE and RE samples were separated, and one network was inferred for each modality. Control samples were excluded from the SC-ION analysis. The final output is therefore four separate networks: Muscle-only EE, Muscle-only RE, Blood-to-muscle EE, and Blood-to-muscle RE.

##### Preparation of input data

For the muscle-only network, the normalized values for the muscle phosphoproteome were used as the regulator matrix. Since SC-ION cannot incorporate missing values, the muscle phosphoproteome was filtered for phosphosites quantified in at least n = 3 samples per group timepoint combination with no more than 40% missing values, and then missing values were imputed with Multiple Imputation by Chained Equations (MICE)^206^ by using the average imputed value from 15 multiple imputations. This imputed matrix was only used for SC-ION inference and no other analyses in this manuscript. This imputed matrix was filtered for phosphosites located on TFs using “The Human Transcription Factors” database (https://humantfs.ccbr.utoronto.ca/). For the blood-to-muscle network, the normalized values for the blood metabolites were used as the regulator matrix. For both networks, the normalized values for the muscle transcriptome were used as the target matrix. The target matrix was filtered for DA transcripts (adjusted p-value < 0.05 in any exercise-with-control contrast). All matrices were split into EE and RE samples prior to running SC-ION.

##### Clustering

SC-ION incorporates an optional clustering step to group together features with similar expression profiles prior to inferring the network. For this implementation, c-means clustering (see C-means clustering) was used to cluster only the regulator and target features used as inputs to each network.

##### Edge trimming

All of the edges in the inferred networks are assigned a weight which indicates the relative confidence in the prediction. To trim the networks based on edge weights, a permutation analysis was performed by randomly shuffling the regulator and target matrices and running SC-ION 100 times. The edge weights were then ranked and assigned a *p*-value based on a permutation test. The *p*-values were then adjusted using Benjamini-Hochberg (BH). The networks were originally trimmed to edge weights with a BH-adjusted *p*-value < 0.05 as determined by the permutation test. However, this resulted in overly dense networks, so a more conservative edge weight cutoff (edge weight > 0.1) was chosen based on the edge weight distribution.

##### Importance score calculation

Following network inference, all edges from the two EE networks were merged using a union (keeping all edges) to form a combined EE network (same for the combined RE network). A Network Motif Score (NMS) was calculated for each feature in the combined EE and combined RE network.^77^ Feed-forward loop (FFL), diamond, and 3-chain motifs were used as input for the NMS (Figure 6a). The total number of motifs, as well as the features within each motif, were determined using the NetMatchStar application in Cytoscape.^207^ This resulted in two NMS, one for EE and one for RE. Further, an average NMS was calculated to determine the overall importance of each feature in the combined EE+RE network.

##### Visualization

The combined EE and RE networks were merged using a union to form the final, combined EE+RE network, which is reported along with the NMS in Supplementary Table 6. All network visualization on the combined EE+RE network was performed in Cytoscape.

##### Validation of TFEB targets

The EE+RE network was filtered to the subset of TFEB plus its first-degree (directly upstream or directly downstream) neighbors. Then, three published ChIP-Seq datasets were used to subset this network to only the downstream targets with demonstrated TFEB binding (237/719 predicted targets, 33%)^114,115,208,209^.

#### Secreted factor analysis

Differential abundance results were aligned according to relative timepoint and tissue. First, DA features from RNAseq, global proteomics, and phosphoproteomics were identified after filtering for features that had an FDR adjusted p-value of < 0.05. No limitations were placed on direction of effect. DA features were mapped to their respective gene symbol using the human feature-to-gene map as described above. Respective gene symbols were then used to connect DA features in muscle, adipose, or blood cells to corresponding circulating plasma proteins as measured by Olink. If tissue DA features matched DA protein features in the plasma, they were considered as putative exercise-induced, tissue-secreted factors. These features were then annotated with their likelihood to be localized to the extracellular domain using subcellular localization database “COMPARTMENTS”,^101^ which provides location scores ranging from 0 to 5, with higher scores indicating greater confidence in localization. The “All channels integrated” file for human proteins was obtained from (https://compartments.jensenlab.org/Search). To calculate the extracellular score, we selected the highest score among the following compartments associated with extracellular localization: *Extracellular region*, *Extracellular space*, *Extracellular exosome*, and *Extracellular vesicle*.

For CCN1-receptor correlations within each tissue, biweight midcorrelation was conducted as previously described,^210^ given its robustness to outlier-driven associations across multi-gene correlations. Pearson correlation was used for inter-tissue CCN1 correlation (i.g., SKM CCN1 vs. AT CCN1). Temporal concordance was assessed at the gene level and was defined as tissue differential abundance during exercise or at an early post-exercise timepoint (during exercise, 10 min post, or 15-45 min post) with a corresponding significant plasma change at the same or a later post-exercise timepoint (15-45 min, 3.5-4 hr, or 24 hr), or tissue differential abundance at 3.5-4 hr post-exercise with a corresponding plasma change at 3.5-4 hr or 24 hr. Directionality concordance was assessed at the level of each individual tissue-omics-timepoint feature entry and was defined as agreement between the sign of the tissue log fold-change and that of the corresponding plasma protein (the plasma timepoint with the largest absolute log fold-change was used to represent each protein’s overall direction).

### QUANTIFICATION AND STATISTICAL ANALYSIS

#### Statistical parameter reporting

All relevant statistical parameters for the analyses performed in this study such as sample size (*N*), center and spread (e.g. mean/median, standard deviation/error), statistical methodology (e.g. mixed-effects linear model) and significance cutoffs (e.g. adjusted *p*-value < 0.05) are reported in the main text, figure legends, Methods, and/or supplementary information. Where appropriate, methods used to determine the validity of certain statistical assumptions are discussed.

#### Statistical limitations

This report presents analyses and results for 175 participants randomized before the suspension of the MoTrPAC study due to the COVID-19 pandemic with at least one biospecimen at the baseline acute bout. The main post-suspension MoTrPAC study will include 1,541 participants randomized under a slightly modified protocol.^37^ The current paper focuses on evaluating feasibility and generating hypotheses for the main study. Results should be interpreted with caution for several reasons: 1) small sample sizes reduce the power to detect even moderate effects;^211^ 2) simple randomization of small groups can create imbalances in both known and unknown confounders;^212,213^ and 3) NIH initiatives have long emphasized caution in interpreting small studies to enhance reproducibility.^214^

Although blocked randomization with site-based stratification was employed, discrepancies in regulatory approvals, start-up times, and pandemic-related interruptions resulted in imbalanced and small sample sizes. Specific limitations in the baseline (pre-intervention) data include: 1) subgroup sample sizes based on intervention group, sex, and timepoint as small as 4 participants; 2) limited generalizability, as 50% of controls with biosamples were randomized at two of the ten sites, and 74% of participants at four sites; and 3) “sex differences” may reflect “body composition differences” due to insufficient data to disentangle sex from body composition, and variations in DXA machine operators and brands across sites.

Testing for heterogeneity of response in small subgroups was largely avoided. As Brookes et al. show,^215^ interaction effects must be at least double the main effect to achieve 80% power. While the EE and RE groups each had around 70 participants with biosamples, providing 80% power to detect a 0.5 effect size (difference in means/SD) using a two-sample t-test (alpha = 0.05, two-sided), for tests of interaction effects to have 80% power an interaction effect size > 1 would be required, which is considered large.^216^ Consequently, heterogeneity of response was explored only descriptively within the larger randomized groups (EE/RE), with some inferential statistics (e.g., confidence intervals, p-values) emphasizing interval estimation. Our aim was to present the results from a hypothesis-generating perspective, following Ioannidis’s cautionary guidance,^211^ and to lay a solid foundation for future analyses in the main MoTrPAC study.

### Additional resources

#### Package assembly and distribution

The *MotrpacHumanPreSuspensionAnalysis* R package (https://github.com/MoTrPAC/MotrpacHumanPreSuspensionAnalysis) contains functions to generate visualizations, as well as data objects for differential abundance analysis, summary statistics for normalized expression data, feature to gene mapping, and molecular signature datasets as described in the methods.

The *MotrpacHumanPreSuspensionData* R package (https://github.com/MoTrPAC/MotrpacHumanPreSuspensionData) is a package containing individual level data, which cannot be made openly available and can be accessed with approval from a data access committee. See motrpac-data.org for details.

Data used in the preparation of this article were obtained from the Molecular Transducers of Physical Activity Consortium (MoTrPAC) database, which is available for public access at motrpac-data.org. The specific collection versions are: quant-id/human-precovid/c1.0, analysis/human-precovid-sed-adu/c2.0, phenotype/humanprecovid-sed-adu/c3.0.

Lastly, the motrpac-human-presuspension-acute GitHub repository (https://github.com/MoTrPAC/motrpac-human-presuspension-acute) contains code and individual parameters for each figure panel, utilizing the MotrpacHumanPreSuspensionAnalysis and MotrpacHumanPreSuspensionData packages.

## Acknowledgements

The MoTrPAC Study is supported by NIH grants U24OD026629 (Bioinformatics Center), U24DK112349, U24DK112342, U24DK112340, U24DK112341, U24DK112326, U24DK112331, U24DK112348 (Chemical Analysis Sites), U01AR071133, U01AR071130, U01AR071124, U01AR071128, U01AR071150, U01AR071160, U01AR071158 (Clinical Centers), U24AR071113 (Consortium Coordinating Center), U01AG055133, U01AG055137, U01AG055135, U01AG070959, U01AG070960, and U01AG070928 (Pre-Clinical Animal Sites).

Additional grant funding: J.M.R: K23 HL150327, R03OD038387; P.R: K23HL177335-01; D.H.K: K23HL164980, 23CDA1040581; B.R: 5U01AR071150, 2U01AR071130; E.R: P30DK072476; E. K.:R01AG066474; K.C.B:1K01HL177266-01A1; K.M.:R01AG089069; M.R:NIH 5 T32 GM135066; N.M and S.E: P30AG094848; Z.C:NIH R00 HL159241; R.G: R01NR019628, R01DK081572, 21CVD01 (Leducq Foundation), R01HL133870; M.E.L, S.M, and M.T.W: The Wu Tsai Human Performance Alliance at Stanford and the Joe and Clara Tsai Foundation. T.C. and E.V: P30AG044271; R.S.R: I01BX003271-05A1, 1IK6BX007133-01; G.M.M.: R03 AG102042.

## Authorship Information

**Lead Contact:** Daniel H. Katz

**Lead Authors:** Daniel H. Katz, Christopher A. Jin, Gina M. Many, Gregory R. Smith, Hasmik Keshishian, Natalie M. Clark, Gayatri Iyer

**Second Authors:** Cheehoon Ahn, Malene E. Lindholm, Tyler J. Sagendorf, Nikolai G. Vetr

**Writing Group Members:** David Amar, Jacob L. Barber, Anna R. Brandt, Paul M. Coen, Yongchao Ge, Patrick Hart, Fang-Chi Hsu, Byron C. Jaeger, Shruti Marwaha, David JimenezMorales, Damon T. Leach, D. R. Mani, Samuel Montalvo, Hanna Pincas, Prashant Rao, James A. Sanford, Kevin S. Smith, Gaurav Tiwari, Anne M. Weitzel

**Senior Leadership:** Joshua N. Adkins, Euan A. Ashley, Charles F. Burant, Steven A. Carr, Robert E. Gerszten, Bret H. Goodpaster, Michael E. Miller, Stephen B. Montgomery, Venugopalan D. Nair, Jeremy M. Robbins, Stuart C. Sealfon, Michael P. Snyder, Lauren M. Sparks, Russell Tracy, Scott Trappe, Martin J. Walsh, Matthew T. Wheeler, Ashley Y. Xia

**Co-Corresponding:** Stuart C. Sealfon, Robert E. Gerszten, Scott Trappe, Charles F. Burant, Bret H. Goodpaster

**Lead Analyst:** Christopher A. Jin

**Analysis/Figure Team:** Cheehoon Ahn, David Amar, Jacob L. Barber, Natalie M. Clark, Yongchao Ge, Patrick Hart, Fang-Chi Hsu, Gayatri Iyer, Byron C. Jaeger, David Jimenez-Morales, Christopher A. Jin, Daniel H. Katz, Hasmik Keshishian, Damon T. Leach, Malene E. Lindholm, D. R. Mani, Gina M. Many, Samuel Montalvo, Prashant Rao, Tyler J. Sagendorf, Gregory R. Smith, Nikolai G. Vetr

### MoTrPAC Study Group

**Bioinformatics Center:** David Amar, Trevor Hastie, David Jimenez-Morales, Daniel H. Katz, Malene E. Lindholm, Shruti Marwaha,Samuel Montalvo, Robert Tibshirani, Jay Yu, Jimmy Zhen, Euan A. Ashley, Matthew T. Wheeler

**Biospecimens Repository:** Sandra T. May, Jessica L. Rooney, Russell Tracy

**Data Management, Analysis, and Quality Control Center:** Catherine Gervais, Fang-Chi Hsu, Byron C. Jaeger, David Popoli, Joseph Rigdon, Courtney G. Simmons, Cynthia L. Stowe, Michael E. Miller

Exercise Intervention Core: W. Jack Rejeski

**NIH:** Ashley Y. Xia

Preclinical Animal Study Sites: Sue C. Bodine, R. Scott Rector

**Chemical Analysis Sites:** Hiba Abou Assi, Mary Anne S. Amper, Brian J. Andonian, Isaac K. Attah, Jacob L. Barber, Kevin Bonanno, Clarisa Chavez Martinez, Natalie M. Clark, Johanna Y. Fleischman, David A. Gaul, Yongchao Ge, Marina A. Gritsenko, Joshua R. Hansen, Patrick Hart, Zhenxin Hou, Chelsea M. Hutchinson-Bunch, Olga Ilkayeva, Gayatri Iyer, Pierre M. Jean-Beltran, Christopher A. Jin, Maureen T. Kachman, Hasmik Keshishian, Damon T. Leach, Minghui Lu, D. R. Mani, Gina M. Many, Nada Marjanovic, Nikhil Milind, Matthew E. Monroe, Ronald J. Moore, Venugopalan D. Nair, German Nudelman, Nora-Lovette Okwara, Vladislav A. Petyuk, Paul D. Piehowski, Hanna Pincas, Wei-Jun Qian, Prashant Rao, Abraham Raskind, Alexander Raskind, Stas Rirak, Jeremy M. Robbins, Margaret Robinson, Tyler J. Sagendorf, James A. Sanford, Gregory R. Smith, Kevin S. Smith, Yifei Sun, Gaurav Tiwari, Mital Vasoya, Nikolai G. Vetr, Alexandria Vornholt, Anne M. Weitzel, Yilin Xie, Xuechen Yu, Elena Zaslavsky, Zidong Zhang, Bingqing Zhao, Joshua N. Adkins, Charles F. Burant, Steven A. Carr, Clary B. Clish, Facundo M. Fernandez, Robert E. Gerszten, Stephen B. Montgomery, Christopher B. Newgard, Eric A. Ortlund, Stuart C. Sealfon, Michael P. Snyder, Martin J. Walsh

**Clinical Sites:** Cheehoon Ahn, Alicia Belangee, Bryan C. Bergman, Daniel H. Bessesen, Gerard A. Boyd, Anna R. Brandt, Nicholas T. Broskey, Toby L. Chambers, Clarisa Chavez Martinez, Maria Chikina, Alex Claiborne, Zachary S. Clayton, Paul M. Coen, Katherine A. Collins-Bennett, Tiffany M. Cortes, Gary R. Cutter, Matthew Douglass, Daniel E. Forman, Will A. Fountain, Aaron H. Gouw, Kevin J. Gries, Fadia Haddad, Joseph A. Houmard, Kim M. Huffman, Ryan P. Hughes, John M. Jakicic, Catherine M. Jankowski, Neil M. Johannsen, Johanna L. Johnson, Erin E. Kershaw, Dillon J. Kuszmaul, Bridget Lester, Colleen E. Lynch, Edward L. Melanson, Cristhian Montenegro, Kerrie L. Moreau, Masatoshi Naruse, Bradley C Nindl, Tuomo Rankinen, Ulrika Raue, Ethan Robbins, Kaitlyn R. Rogers, Renee J. Rogers, Irene E. Schauer, Robert S. Schwartz, Chad M. Skiles, Lauren M. Sparks, Maja Stefanovic-Racic, Andrew M. Stroh, Kristen J. Sutton, Anna Thalacker-Mercer, Todd A. Trappe, Caroline S. Vincenty, Elena Volpi, Katie L. Whytock, Gilhyeon Yoon, Thomas W. Buford, Dan M. Cooper, Sara E. Espinoza, Bret H. Goodpaster, Wendy M. Kohrt, William E. Kraus, Nicolas Musi, Shlomit Radom-Aizik, Blake B. Rasmussen, Eric Ravussin, Scott Trappe

### MoTrPAC Study Group Acknowledgements

Nicole Adams, Abdalla Ahmed, Andrea Anderson, Carter Asef, Arianne Aslamy, Marcas M. Bamman, Jerry Barnes, Susan Barr, Kelsey Belski, Will Bennett, Amanda Boyce, Brandon Bukas, Emily Carifi, Chih-Yu Chen, Haiying Chen, Shyh-Huei Chen, Samuel Cohen, Audrey Collins, Gavin Connolly, Elaine Cornell, Julia Dauberger, Carola Ekelund, Shannon S. Emilson, Jerome Fleg, Nicole Gagne, Mary-Catherine George, Ellie Gibbons, Jillian Gillespie, Aditi Goyal, Bruce Graham, Xueyun Gulbin, Jere Hamilton, Leora Henkin, Andrew Hepler, Lidija Ivic, Ronald Jackson, Andrew Jones, Lyndon Joseph, Leslie Kelly, Gary Lee, Adrian Loubriel, Ching-ju Lu, Kristal M. Maner-Smith, Ryan Martin, Padma Maruvada, Alyssa Mathews, Curtis McGinity, Lucas Medsker, Kiril Minchev, Samuel G. Moore, Michael Muehlbauer, Anne Newman, John Nichols, Concepcion R. Nierras, George Papanicolaou, Lorrie Penry, June Pierce, Megan Reaves, Eric W. Reynolds, Teresa N. Richardson, Jeremy Rogers, Scott Rushing, Santiago Saldana, Rohan Shah, Samiya M. Shimly, Cris Slentz, Deanna Spaw, Debbie Steinberg, Suchitra Sudarshan, Alyssa Sudnick, Jennifer W. Talton, Christy Tebsherani, Nevyana Todorova, Mark Viggars, Jennifer Walker, Michael P. Walkup, Anthony Weakland, Gary Weaver, Christopher Webb, Sawyer Welden, John P. Williams, Marilyn Williams, Leslie Willis, Yi Zhang, Frank Booth, Karyn A. Esser, Laurie Goodyear, Andrea Hevener, Ian Lanza, Jun Li, K Sreekumaran Nair

## Disclosures

Disclaimer: The content of this manuscript is solely the responsibility of the authors and does not necessarily represent the views of the National Institutes of Health, or the United States Department of Health and Human Services.

## Declaration of Interests

B.H.G has served as a member of scientific advisory boards; J.M.R is a consultant for Edwards Lifesciences; Abbott Laboratories; Janssen Pharmaceuticals; M.P.S is a cofounder and scientific advisor of Xthera, Exposomics, Filtricine, Fodsel, iollo, InVu Health, January AI, Marble Therapeutics, Mirvie, Next Thought AI, Orange Street Ventures, Personalis, Qbio, RTHM, SensOmics. M.P.S is a scientific advisor of Abbratech, Applied Cognition, Enovone, Jupiter Therapeutics, M3 Helium, Mitrix, Neuvivo, Onza, Sigil Biosciences, Captify Inc, WndrHLTH, Yuvan Research, Ovul, Erudio and Lyten; S.B.M is a member of the scientific advisory board for PhiTech, MyOme and Valinor Therapeutics; S.A.C is on the scientific advisory boards of PrognomIQ, MOBILion Systems, Kymera, and Stand Up2 Cancer; S.C.S is a founder of GNOMX Corp, leads its scientific advisory board and serves as its temporary Chief Scientific Officer; A.V is a consultant for GNOMX Corp; B.C.N is a member of the Science Advisory Council, Institute of Human and Machine Cognition, Pensacola, FL; E.E.K is a consultant for NodThera and Sparrow Pharmaceuticals and a site PI for clinical trials for Arrowhead Pharmaceuticals; E.A.A is: Founder: Personalis, Deepcell, Svexa, Saturnus Bio, Swift Bio. Founder Advisor: Candela, Parameter Health. Advisor: Pacific Biosciences. Non-executive director: AstraZeneca, Dexcom. Publicly traded stock: Personalis, Pacific Biosciences, AstraZeneca. Collaborative support in kind: Illumina, Pacific Biosciences, Oxford Nanopore, Cache, Cellsonics; G.R.C is a part of: Data and Safety Monitoring Boards: Applied Therapeutics, AI therapeutics, Amgen-NMO peds, AMO Pharma, Argenx, Astra-Zeneca, Bristol Meyers Squibb, CSL Behring, DiamedicaTherapeutics, Horizon Pharmaceuticals, Immunic, Inhrbx-sanfofi, Karuna Therapeutics, Kezar Life Sciences, Medtronic, Merck, Meiji Seika Pharma, Mitsubishi Tanabe Pharma Holdings, Prothena Biosciences, Novartis, Pipeline Therapeutics (Contineum), Regeneron, Sanofi-Aventis, Teva Pharmaceuticals, United BioSource LLC, University of Texas Southwestern, Zenas Biopharmaceuticals. Consulting or Advisory Boards: Alexion, Antisense Therapeutics/Percheron, Avotres, Biogen, Clene Nanomedicine, Clinical Trial Solutions LLC, Endra Life Sciences, Genzyme, Genentech, Immunic, Klein-Buendel Incorporated, Kyverna Therapeutics, Inc., Linical, Merck/Serono, Noema, Neurogenesis, Perception Neurosciences, Protalix Biotherapeutics, Regeneron, Revelstone Consulting, Roche, Sapience Therapeutics, Tenmile. G.R.C is employed by the University of Alabama at Birmingham and President of Pythagoras, Inc. a private consulting company located in Birmingham AL. J.M.J is on the Scientific Advisory Board for Wondr Health, Inc.; P.M.J.B is currently an employee at Pfizer, Inc., unrelated to this project; R.R is a scientific advisor to AstraZeneca, Neurocrine Biosciences, and the American Council on Exercise, and a consultant to Wonder Health, Inc. and seca.; B.J receives consulting fees from Perisphere Real World Evidence, LLC, unrelated to this project. M.J.W. is a minority partner of Arch Venture Partners, Inc. and co-founder and scientific advisory board member of EnGenome Biosciences, Inc.

## Notes

https://motrpac-data.org/

### Summary of Updates

1. This revision incorporates updates from the MoTrPAC collection c2.0 data release (analysis/human-precovid-sed-adu/c2.0). Key changes include: Data Updates: Corrected minor data and coding errors in muscle transcriptomics, muscle ATAC-seq, muscle proteomics/phosphoproteomics, blood proteomics, and updated metabolite name mappings across metabolomics tables. Manuscript Impact: Propagated changes across data files, figures, tables, and text. Core conclusions remain largely unchanged, but exact effect sizes, p-values, and counts of significant molecular changes have been updated for affected tissue-ome combinations. Full release notes: https://motrpac-data.org/knowledge-center/data-collections/analysis-results/analysis-human-precovid-sed-adu 2. New Content: Expanded comparisons between molecular data and clinical phenotypes, and added a Supplemental Results section. 3. Display Items: Reordered and revised several figures for clarity. 4. Author List: Added authors Anne Marie Weitzel, Shruti Marwaha, and Gaurav Tiwari, and updated author position for Nik Vetr.

