## Supplementary Results for "Multi-Omic, Multi-Tissue Responses to Acute Exercise in Sedentary Adults: Findings from the Molecular Transducers of Physical Activity Consortium"

**Study design and participant characteristics** **1**
**Exercise induced multi-omic changes** **1**
**Cell type deconvolution analysis** **2**
**Exercise modality-specific effects across tissues and omes** **3**
Trait associations 4
**Temporal molecular responses with acute exercise** **4**
Timepoint-Specific Omic and Pathway Enrichments 4
Multi-Omic Trajectory Characterization via Fuzzy C-Means Clustering 4
Cross-Tissue Overlap and Novel Transcriptional Trajectories 5
Latent Variable Modeling of Cross-Tissue Behavior (PLIER) 6
**Muscle ATAC-seq QC Analysis** **7**
Muscle ATAC-seq QC Methods 8
Supplementary Figure Legends 8

### Study design and participant characteristics

Across features highlighted in the manuscript, each tissue-ome-group-timepoint combination included a median of N = 31 participants (interquartile range [IQR] 18-36). In proteomic and phosphoproteomic assays, which have feature-level missingness inherent to the measurement platform, the number of sample measurements for each feature is therefore variable. Per ome-specific standards, any features with greater than 70% missingness in each tissue were removed from further analysis and median of N = 17 samples went into proteomic and phosphoproteomic measurements per tissue-ome-group-timepoint combination. The distribution of the number of samples for all features measured in mass-spectrometry based proteomic platforms is highlighted in Extended data Figure 1c.

While exercise group, timepoint, and covariates like age and sex accounted for a large proportion of the variation in molecular features, for most tissue-ome combinations, effects attributable to individual differences as captured by participant ID (PID) accounted for a larger percentage of variation (Extended Data Fig. 1d).

### Exercise induced multi-omic changes

While most DA molecules were tissue-specific (i.e. DA in only one tissue), we sought to explore in greater detail those features which were overlapping (Fig. 2b). A total of 3,620 features were DA in both blood and SKM, driven by shared transcripts, while SKM and AT shared 88 unique DA features (Fig. 2b, Extended Data Fig. 2c, Supplementary Table 1, Supplementary Results). Overlap in DA features depended on ome, which is also impacted by sample availability (Fig. 1b-c, Extended Data Fig. 1c, Supplementary Table 1, Methods). In omes with greater sample representation across timepoints (i.e. transcriptomics and metabolomics), agreement in SKM and blood was higher than SKM and AT. Of the 8,897 DA transcripts in SKM, ~40% were also DA in blood while less than 1% were DA in AT. Of 110 DA metabolites in SKM, ~64% were also DA in blood, but just 11% in AT. Phosphoproteomic agreement between SKM and AT was low, with less than 1% of 4253 DA phosphosites in SKM being DA in AT (Supplementary Table 2). However, caution must be exerted in interpretation as proteomic and phosphoproteomic assessment was limited to one timepoint in AT, versus 3 in SKM. Differentially accessible peaks could only be detected in SKM (n = 1209 via ATAC-seq), while the greatest number of differentially methylated regions (n = 580) were observed in blood (Extended Data Fig. 2c). Across tissues, there was a single methylated region observed in both SKM and AT (Extended Data Fig. 2c). This peak was annotated to a long non-coding RNA (lncRNA) gene (ENSG00000245482) that is an antisense RNA to *ALG10* (Alpha-1,2-Glucosyltransferase), a protein involved in N-linked glycosylation.[1]

Corresponding with ome coverage across tissues, the majority of shared features across all three tissues were transcripts (N=47) (Fig. 2c, Extended Data Fig. 2c). Among shared DA features, the direction of exercise effect varied across tissues and modalities (Fig. 2c). Those that shared directionality across tissues included the nuclear receptors *NR4A1* (aka *NUR77*) and *NR4A2*, the scaffold protein *NEDD9* (Enhancer of filamentation 1), the EGFR ligand, *AREG*, and the NFkB inhibitor, *NFKBIZ*, which was acutely downregulated during EE in the blood, but then displayed elevated expression following both EE and RE in all tissues (Fig. 2d, Extended Data Fig. 2d), with *NFKBIZ* suggesting temporally resolved anti-inflammatory effects of exercise.

Of the 39 shared DA phosphosites, assessed in the AT and SKM, two-thirds shared directionality across tissues (Fig. 2e). Shared DA phosphoproteins included pS920 of CTNND1, a protein involved in c-MYC and CDK1 signaling,[2] decreased with EE in AT and SKM, suggesting a multi-tissue role for this phosphosite in acute exercise. To further investigate phosphospecific signatures that were distinct or shared between SKM and AT, we performed ORA on DA phosphosites using PTMSigDB, which is a site-level pathway database designed for use with phosphoproteomics data (Extended Data Fig. 2e, Supplementary Table 2).[3] The SKM DA phosphosites were significantly enriched in pathways including kinase pathways known to be affected by exercise such as MAPK/ERK, MTOR, CDK1/2/5, and GSK3B. AT and SKM shared 3 enriched pathway signatures that included perturbations related to BYL719, a PI3K inhibitor, and RO4929097, a gamma secretase inhibitor that targets NOTCH signaling[4], and MAP2K2 kinase enrichment.

We then looked at shared pathway enrichment using DA features observed in each individual tissue as well as on features which overlapped in blood/SKM and AT/SKM (i.e. tissue pairs where overlap was sufficient for analysis). Enriched pathways shared across all three tissues included several with known responses to exercise (e.g. cellular response to stress, mitochondrion, IL-6) as well as less well-characterized pathways involved in physiological homeostasis such as cytoskeletal organization and neurogenesis across all three tissues (Fig. 2f, Supplementary Table 2).

In contrast to cross-tissue agreement in DA molecules, we further examined cross-omic agreement within tissues (Extended Data Fig. 2f). While most DA features were DA in only one omic-layer, there were notable numbers of genes that were DA across 2 or more. In particular, 856 genes in SKM were DA by transcriptomics and by phosphoproteomics. In blood, 75 genes had changes in transcription and plasma proteomic levels (Extended Data Fig. 2f).

### Cell type deconvolution analysis

After collapsing the LM22 subsets, blood transcriptomes were deconvolved into seven leukocyte populations across 807 samples from 173 participants spanning the EE, RE, and CON arms. Neutrophils were the dominant estimated population (mean 61.1% of cells), followed by T cells

(11.0%), monocytes (8.8%), NK cells (7.9%), macrophages (5.2%), B cells (4.5%), and mast cells (1.5%).[5]

Estimated neutrophil proportions at pre-exercise were moderately correlated with matched complete blood count neutrophil values (Pearson R = 0.22 across all participants; R = 0.37 after excluding two samples with implausibly low clinical neutrophil counts (3.6% and 4.1%) (Extended Data Fig. 3d). In sensitivity analyses, adding the coarse cell-type fractions as covariates to the primary model had little effect on the blood RNA-seq differential analysis. Across exercise modalities and timepoints, log fold-changes from the covariate-adjusted model were nearly identical to the original estimates (Pearson R = 0.91-0.99, Extended Data Fig. 3e), suggesting cell type did not have substantial impact on the reported DA transcripts from the primary model.

### Exercise modality-specific effects across tissues and omes

To further illustrate pathway-level differences between EE and RE, we applied pathway enrichment via Pre-ranked Correlation-Adjusted MEan-RAnk (CAMERA-PR), which accounts for intergene correlations, to all EE vs. RE DA model comparisons (Extended Data Fig. 4b). At the transcript level in SKM, RE showed stronger enrichment in terms related to muscle remodeling including hypertrophy, HSF1 activation, and RNA splicing (Extended Data Figure 4b and 4c). The HSF1 pattern highlights a temporal offset in activity between EE and RE, with RE upregulation persisting to 24 hours (Extended Data Fig. 4b). The pattern of HSF1 modality differences was similar in blood at 24 hrs (Extended Data Fig. 4c), suggesting more persistent systemic stress-induced response following RE.[6] At the feature level, this is exemplified by *HSPH1* which is strongly induced by EE at 30 minutes post exercise in SKM, but not until 3.5 hours in RE (Extended Data Fig. 4c,d).

EE displayed enhanced mitochondrial enrichments to include fatty acid beta oxidation, mitochondrial translation, and ATP biosynthetic processes in the SKM (Extended Data Fig. 4b). In the AT, EE-specific increases in enrichment for mitochondria-related pathways were observed earlier than in the SKM, at the 30 min post-exercise timepoint. In the blood, RE induced a more pronounced immune response indicated by enrichment in terms related to neutrophil degranulation and T-cell receptor signaling. RE enrichment was also present for the glycoprotein VI mediated activation cascade, a response to blood vessel damage (Supplementary Table 3). Divergent SKM transcriptional l feature responses to EE and RE included known transcriptional regulators of SKM regeneration such as Myogenin (*MYOG)* (Extended Data Fig. 4e).

Divergent phosphoproteomic signatures included pT114 of the anti-apoptotic factor NOL3 (Extended Data Fig. 4e), which increased in RE, with implications for muscle injury prevention.[7] RE also increased phosphorylation of two proteins involved in structural integrity of the Z-disc: FHOD1 (Formin Homology Domain Protein 1, Figure 5F) and Myotilin (Extended Data Fig. 4e), which were decreased in EE and indicative of sarcomeric protection mechanisms following RE. The same pattern was observed for a novel lncRNA intronic to the cytoskeleton gene *LIMA1* (Extended Data Fig. 4e).

#### Trait associations

As described in the section related to Fig. 3, baseline clinical and fitness traits were correlated to pre-exercise molecular features in all participants.

In adipose tissue, large numbers of genes correlated with Homeostatic Model Assessment for Insulin Resistance (HOMA-IR), while relatively few genes were associated with $VO_2$peak (2293 and 38 respectively) (Extended Data Fig. 4g; Supplementary Table 3). Among transcripts associated with both VO2peak and HOMA-IR (n = 24), all showed opposing directions of association. Genes positively associated with HOMA-IR (n=1292) were enriched for similar immune and inflammatory processes, whereas genes negatively associated with HOMA-IR (n=1001) were enriched for fatty-acid oxidation and mitochondrial metabolism (Supplementary Table 3). These findings are consistent with prior studies linking adipose inflammatory tone to insulin resistance.[8] These results support the idea that higher baseline $VO_2$peak is anti-inflammatory,[9] and this anti-inflammatory effect may mediate some of the association with improved cardiometabolic health.

In blood, 141 metabolites were significantly associated with HOMA-IR (Extended Data Fig. 4g; Supplementary Table 3). These metabolites were largely related to lipid metabolism, particularly glycerolipid and phospholipid metabolism, membrane-lipid remodeling, and fatty-acid storage and transport. In skeletal muscle, 41 metabolites were associated with NEFA, while both proteins (493) and transcripts (946) were associated with $VO_2$peak (Extended Data Fig. 4g).

Selected features of interest included *FUT1*, a novel exercise-responsive gene that was positively associated with $VO_2$peak and upregulated in response to both EE and RE in muscle. Muscle expression of *PDK4* was positively associated with NEFA, pairing with its role of promoting lipid oxidation, and was induced in response to both EE and RE. Another NEFA-associated gene was *ANGPTL4*, which showed the highest positive association and per unit increase (beta) of all correlated feature-traits, and increased in muscle with both EE and RE (Supplementary Table 3). ANGPTL4 is a secreted glycoprotein with central metabolic functions including regulation of glucose and lipid metabolism, and angiogenesis.[10]

### Temporal molecular responses with acute exercise

#### Timepoint-Specific Omic and Pathway Enrichments

To expand on the global timepoint-specific enrichment analyses, we examined detailed metabolomic shifts across tissues following acute endurance exercise (EE) and resistance exercise (RE). As a central fuel source for OXPHOS, triglycerides decreased at various timepoints across all tissues in response to EE and RE. Unsaturated fatty acids decreased during (EE) and early post exercise (RE) in blood, which later rebounded (Extended Data Fig 5a).

#### Multi-Omic Trajectory Characterization via Fuzzy C-Means Clustering

To better understand multi-omic trajectories, we analyzed temporal molecular signatures across the post-exercise timepoints using fuzzy c-means clustering, identifying 13 distinct temporal clusters in adipose tissue (AT), and 12 in the blood and SKM (see Methods). Omic contributions to each cluster varied widely (Fig. 4c and expanded in Extended Data Fig. 5b); in SKM, phosphoproteomic changes were more prevalent in trajectories with early maximal responses (muscle clusters 4, 6, 10, 8, and 9). Transcriptomic features were more likely to show a maximal response at 3.5 hours (muscle clusters 12, 1, and 3). Proteomic changes were relatively evenly distributed, though slightly more prevalent in clusters with changes in muscle at 24 hours (muscle clusters 7, 4, and 11). In blood, metabolic features appeared in trajectories with early (blood clusters 4 and 12) and delayed (blood clusters 8 and 11) maximal responses.

When ORA was applied to features assigned to each cluster, multiple pathways emerged across time and tissues (Extended Data Fig. 5b, Supplementary Table 4). For example, oxidative phosphorylation pathways were enriched in transcriptomic and proteomic layers across mid-peaking and late-peaking trajectories, both up and downregulated. Alternatively, autophagy transcriptomic features were significantly enriched in temporal clusters with mid-peaking, upregulated signatures across all three tissues. Triglycerides, as a fuel source, were enriched in temporal trajectories that were downtrending, but each tissue demonstrated different time signatures; triglycerides in SKM and AT favored patterns with later depletion, while blood triglycerides were enriched within clusters with early to mid response patterns.

### Cross-Tissue Overlap and Novel Transcriptional Trajectories

While cluster membership within each tissue was strictly non-overlapping, cross-tissue comparisons revealed modest commonalities (Extended Data Fig. 5c). For example, shared between clusters adipose-8 and muscle-6/10 were transcription factors (TFs) including NR4A1, *NR4A2*, *EGR1*, and *JUN*, as well as key angiogenic regulators like *APOLD1* and *ADAMTS1* (Extended Data Fig. 5d). Importantly, ADAMTS1 plays an anti-angiogenic role,[12] highlighting that molecular brakes are stimulated by exercise temporally along with molecular activators. In AT, *CCN2* displayed the most significant modality-consistent upregulation at 45 minutes post-exercise; the pattern in SKM was similar, although *CCN2* activity persisted to 3.5 hours with RE. Also known as connective tissue growth factor (CTGF), *CCN2* expression is induced in response to VEGF in endothelial cells.[13]

There was also overlap between features in blood and SKM whose trajectories were highest at 3.5 hours post-exercise (Extended Data Fig. 5e). These transcripts include key regulators for a variety of adaptive processes relevant to both tissues induced by EE and/or RE. For example, growth and differentiation factors like *CRK* and *EIF4G2,* are upregulated along with factors to balance these growth signals, such as upregulation of *GRB10 and PPP2CA*[14–17]. This group also contains a handful of novel findings. *SDCBP,* which encodes syntenin-1, plays a role in exosome formation,[18] positioning it as a central regulator for general exerkine signaling. *HLX* is a hematopoietic TF linked to leukemia; in the setting of exercise it may be resetting blood and SKM for differentiation and regeneration.[19]

Our trajectory analysis also identified a trajectory enriched for VEGF pathway transcripts that persist at their highest levels in SKM to 24 hours post exercise (cluster 7, Fig. 4e). These features are candidates for driving long term adaptation, once the initial stress of the acute bout has passed. Of note, some muscle-7 features have increased transcription at 3.5 hours as well (Fig. 4f). These features appear to have no clear role in AT, but in blood seem to roughly split into an up- and downregulated group. For example, *INPP4B* is upregulated in SKM, but downregulated in blood during exercise (EE only). This fits with the known function of *INPP4B* as both tumor suppressor and oncogene, depending on context.[20] *INPP4B* was previously shown to increase with exercise in SKM, but here we show the opposing behavior in blood.[21] As a suppressor of the PI3K-Akt pathway, it may act as counter-regulatory in SKM, but activating in blood.[22] Angiomotin, encoded by the *AMOT* gene, is a key factor in angiogenic patterning,[23] though is not previously described in human exercise adaptation; in rats, exercise is known to affect isoform balance.[24]

Taken together, these analyses not only reveal novel exercise response molecules, but also a nuanced and often tissue-specific temporal regulation of key exercise response pathways. These data can be explored across a range of other enriched pathways, such as autophagy, where blood and SKM exhibited temporally similar pathway induction, but with largely divergent transcriptional programs (see R package).

### Latent Variable Modeling of Cross-Tissue Behavior (PLIER)

To capture coordinated molecular responses across tissues, we applied the pathway-level information extractor (PLIER) method to both the transcriptomic and metabolomic datasets to track latent variables (LVs) of molecules exhibiting consistent cross-tissue behavior (Extended Data Fig. 6a). The PLIER analysis identified many transcriptomic LVs, with several biological pathways highlighted by each LV (Fig. 4g-h). The cross-tissue correlation and total transcripts per LV are shown in Extended Data Fig. 6b-c).

One specific latent variable, LV13, was strongly correlated with ribosomal biogenesis, a process that has been linked to SKM hypertrophy[25], and displayed contrasting patterns of expression in AT, SKM and blood, with decreased expression in AT post RE and SKM post EE and RE at the early time point (15-45m) followed by increased expression SKM at the late timepoint (24h) after both RE and EE, while decreased expression in blood was observed at multiple post-exercise timepoints peaking at 3.5 h across both exercise modalities (Fig. 4g-h). Four response patterns (LV6, LV33, LV73, LV79) highlighted transcription regulatory activity. Key splicing factor genes including *HNRNPU* and *DYRK1A* are associated with LV6, representing strong increases in SKM at 3.5 hours in both EE and RE, but also weaker increases post exercise in AT and blood. LV6 highlights the cross-tissue splicing regulatory relationships with acute exercise, see also Keshishian et al.[26] and Zhang et al.[27] LV79 exercise response overlapped with LV6 at a number of timepoints in blood, but also exhibited strong decreases in expression in muscle post EE and RE, unlike LV6. LV73 was also enriched for the same pathways as LVs 6 and 79, but was also enriched for heat shock protein activity, and represented decreases post exercise at the early timepoint in AT but increases during EE and post exercise in both EE and RE in blood and increases post EE in SKM. This diversity of significant exercise responses by distinct subsets of overlapping pathways reflects the complex nature of exercise response and how transcription regulation encompasses multiple goals and trajectories.

Transporter activity regulation-associated LV54 (Fig. 4g) reflected an RE-specific increase in expression at 3.5 and 24-hour timepoints in SKM and weaker increases following both EE and RE at the earlier post-exercise timepoints (10 and 30m) in blood. Pathway and LV-associated genes include *FHL1* and *MEF2C*, the latter a well-studied muscle TF whose motif is enriched for exercise response genes in this study (Fig. 5h), as well as a previous MoTrPAC exercise training study in rats.[28]

In the companion metabolomic analysis, we highlight 7 LVs (Extended Data Fig. 6e-f; Supplementary Table 4). Five of the 7 LVs were enriched for at least one known metabolite classification (Extended Data Fig. 6f). The cross-tissue correlation and total metabolites per LV are shown in Extended Data Fig. 6g-h). Between 10 and 30 metabolites were associated with each highlighted LV (Extended Data Fig. 6h).

Unsaturated fatty acid-associated LV4 reflects similar patterns of exercise response in blood as acyl carnitine-associated LV17, which is discussed in the main results section, but does not show significant responses in muscle. Aside from LV17, two other LVs emerged emphasizing blood-SKM interaction (Extended Data Fig. 6g). LV11 included a range of metabolites such as amino acids, lactate, pyruvate, and hypoxanthine (Supplementary Table 4). This group collectively increased in blood during EE and following both exercise interventions, but in SKM, values decreased 24 hours post EE and RE. Many of these metabolites are byproducts of energy production (TCA intermediates, glycolysis products) showing up in the plasma.[29] The decrease in SKM at 24 hours may point to an overcompensation in clearance. Conversely, LV65 represented an alternative blood-SKM interaction. This metabolite set included citrate, isocitrate, malate, 3-hydroxybutyrate and short chain acylcarnitines (Supplementary Table 4). In contrast to LV11, these features peaked in blood *and* SKM early after both exercise modalities, suggesting weaker clearance from SKM as compared to LV11. As a whole, these groups of features may represent a useful marker of exercise demand and adaptation.

### Muscle ATAC-seq QC Analysis

We present an extended exploration of the SKM ATAC-seq data and point out potential limitations of the difference-in-changes model arising from the technical structure specific to the SKM ATAC-seq data (Fig. S1a). Two sources of technical variation are directly accounted for already in the model: 1) Due to practical limitations, samples from each participant were generally grouped together within their own nuclei isolation batch, leading to correlation between nuclei isolation batch and participant. These effects are attenuated by the fact that each participant serves as their own pre-exercise control within the mixed effects model with a random effect of participant (see Methods). 2) FRiP, a standard covariate in ATAC-seq analysis, was also included in the ~M0 model (see Supplemental Methods, below) due to low values in control libraries at 24hr and for resistance libraries overall (Fig. S1b). These adjustments were effective: In PCA analysis of unadjusted and adjusted voom-transformed counts, PC1 variation reduced from 49.67% to 21.27%, with correlations between the adjusted PCs and sex, age, BMI, site, and FRiP reduced or eliminated, demonstrating an even distribution of these variables across groups and timepoints (Fig. S1c). However, visually, clustering between group:timepoint (Fig. S1d) was attenuated, suggesting a smaller acute exercise signal relative to residual variance in these data.

In post-hoc analyses, three additional factors were identified, and our analysis shows that this variation may have a greater impact on the control group.

Nuclei isolation batch. The order of nuclei isolation was non-random (first controls, followed by EE, then RE). PCA analysis form ~M0 revealed an association between PC1 and nuclei isolation batch driven by group effect, with residual effect of isolation order within the control group (Fig. S1e).

Uneven distribution of sequencing batch. Owing to barcode compatibility, while at least one sequencing batch (4_571) sequenced samples from all three randomized groups, three sequencing batches contained samples from only one of the three groups (Fig. S1a).

Freeze/thaw samples. The control group had a higher number of samples requiring thawing and re-tagmentation owing to QC failures of original samples. The control group had 15 such samples, compared to 7 and 4 in EE and RE respectively. In PCA analysis, ~M0-PC2 visually separates libraries generated from fresh nuclei from those generated from previously isolated nuclei within all three groups (Fig. S1f-g).

Taken together, these factors may limit the separation of technical variation from biological signal when comparing exercise groups to control. This may explain why the number of peaks reported by the difference-in-changes analysis may be limited (Fig. 2a). However, due to the correlation between these factors and group, they were not included in the main model.

Within-group analyses, without comparison to controls, are released for all omes, including ATAC-seq, within the accompanying data package and provide an alternative approach that returns substantially more significant peaks (Fig. S1h). The within-group approach does not take into account potential circadian, feeding, or study effects that could be identified by comparisons against controls. That said, this analysis in the ATAC-seq control group returns zero significant peaks across all timepoints in the within-group analysis, suggesting these control effects may be attenuated in ATAC-seq, contrary to other omes. Further, of the significant peaks identified by the difference-in-differences model, 95% are corroborated by the within-group approach (Fig. S1i). The differences-in-differences model compounds variance from both the exercised and control groups, so biological changes may fail to reach significance when applying the same threshold (FDR < 0.05). Future efforts using alternative models to manage the above variability, while retaining time and group effects, could provide a more comprehensive understanding of chromatin accessibility changes resulting from an acute exercise bout.

### Muscle ATAC-seq QC Methods

Six libraries were dropped due to sample swaps (two libraries) and instances of one collection timepoint per PID (four libraries), resulting in a final 111 libraries generated for 33 participants. Voom-transformed counts were assessed through PCA before and after isolating group:timepoint signal by regressing out covariates included in the standard model (~M0, Formula 1) aside from randomGroupCode and Timepoint. PCA was carried out on all samples together, and within randomGroupCode. PC1-PC5 were correlated (Pearson’s) with known metadata and 15 representative QC metrics that were hand picked from highly correlated clusters of all 66 numeric QC variables in the metadata. Discrete variables were modeled using dummy variables (randomGroupCode, Timepoint, Sex, codedsiteid, Seq_batch_short, nuclei2atac_lag). Nuclei isolation batch, which reflects the order of nuclei isolation, remained continuous for this purpose.

**Formula 1 / ~M0**: ~ BMI + Sex + calculatedAge + codedsiteid + FRiP + Timepoint + randomGroupCode

### Supplementary Figure Legends

#### **Fig. S1**

a) **Metadata by PID**. Skeletal muscle ATAC-seq experimental design per PID (row). For each timepoint (column), the order of nuclei isolation (left section), nuclei usage (center section), and sequencing batch (right section) are detailed. Sample batch numbers reflect the order in which nuclei were isolated. Use of previously isolated nuclei is indicated by a number printed in the cell. The reuse of nuclei was more concentrated in the controls (15 of 40 controls, 7 of 33 endurance, 4 of 38 resistance) and not consistent between individuals. Importantly, 20 of 33 individuals had libraries sequenced in different batches.

b) **Distribution of FRiP**. Boxplots show distribution by group and timepoints, with individual points plotted for each library.

c) **Correlation of PCs with metadata and QC Metrics**. Pearson's correlation coefficients from comparisons including all samples are plotted. Correlation coefficients are printed for significant correlations ($p < 0.05$).

d) **~M0 PCA plot**. Scatterplot of PC1 and PC2 when ~M0 covariates are adjusted for while protecting for group and timepoint (n = 111, color = group, shape = timepoint, linetype = timepoint).

e) **Nuclei isolation order relationship with ~M0 PC1**. Regression lines and correlation coefficients (Spearman's) provided by group and for all samples (color = group, shape = timepoint).

f-g) **Distribution of ~M0 PC2 by use of frozen nuclei**. (f) distributions shown by group, (g) further separates samples by (color = use of frozen nuclei, shape = timepoint).

h) **Count of significant features by group and timepoint**. The significance threshold of FDR $< 0.05$ was applied. Within-group control results returned zero significant peaks across all timepoints so they are not included.

i) **Overlapping and unique significant features between modeling approaches**. Count of significant features (FDR $< 0.05$) unique to the difference-in-changes approach (pink), shared between difference-in-changes and within-group modeling (green), and unique to the within-group analysis (yellow) for each exercise modality and timepoint.

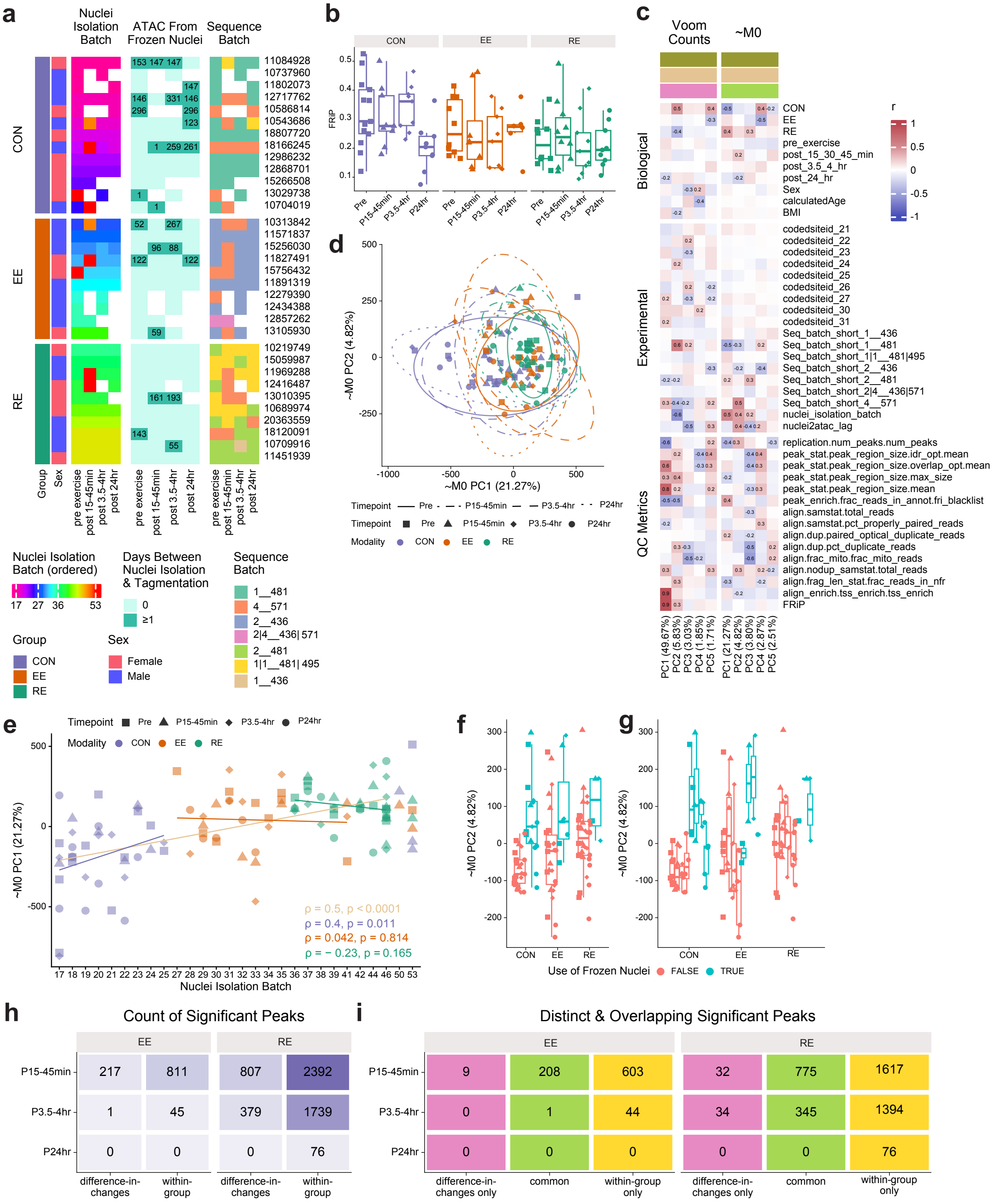

a
Nuclei Isolation Batch
ATAC From Frozen Nuclei
Sequence Batch
CON
EE
RE
Group
Sex
pre exercise
post 15-45min
post 3.5-4hr
post 24hr
Nuclei Isolation Batch (ordered)
17 27 36 53
Days Between Nuclei Isolation & Tagmentation
0
≥1
Sequence Batch
1__481
4__571
2__436
2|4__436| 571
2__481
1|1__481| 495
1__436
Group
CON
EE
RE
Sex
Female
Male
b
CON
EE
RE
FRiP
Pre
P15-45min
P3.5-4hr
P24hr
c
Voom Counts
~M0
Biological
Experimental
QC Metrics
CON
EE
RE
pre_exercise
post_15_30_45_min
post_3.5_4_hr
post_24_hr
Sex
calculatedAge
BMI
codedsiteid_21
codedsiteid_22
codedsiteid_23
codedsiteid_24
codedsiteid_25
codedsiteid_26
codedsiteid_27
codedsiteid_30
codedsiteid_31
Seq_batch_short_1__436
Seq_batch_short_1__481
Seq_batch_short_1|1__481|495
Seq_batch_short_2__436
Seq_batch_short_2__481
Seq_batch_short_2|4__436|571
Seq_batch_short_4__571
nuclei_isolation_batch
nuclei2atac_lag
replication.num_peaks.num_peaks
peak_stat.peak_region_size.idr_opt.mean
peak_stat.peak_region_size.overlap_opt.mean
peak_stat.peak_region_size.max_size
peak_stat.peak_region_size.mean
peak_enrich.frac_reads_in_annot.fri_blacklist
align.samstat.total_reads
align.samstat.pct_properly_paired_reads
align.dup.paired_optical_duplicate_reads
align.dup.pct_duplicate_reads
align.frac_mito.frac_mito_reads
align.nodup_samstat.total_reads
align.frag_len_stat.frac_reads_in_nfr
align_enrich.tss_enrich.tss_enrich
FRiP
PC1 (49.67%)
PC2 (5.83%)
PC3 (3.03%)
PC4 (1.85%)
PC5 (1.71%)
PC1 (21.27%)
PC2 (4.82%)
PC3 (3.80%)
PC4 (2.87%)
PC5 (2.51%)
r
d
~M0 PC2 (4.82%)
~M0 PC1 (21.27%)
Timepoint Pre P15-45min P3.5-4hr P24hr
Modality CON EE RE
e
~M0 PC1 (21.27%)
Nuclei Isolation Batch
ρ = 0.5, p < 0.0001
ρ = 0.4, p = 0.011
ρ = 0.042, p = 0.814
ρ = − 0.23, p = 0.165
f
g
~M0 PC2 (4.82%)
Use of Frozen Nuclei FALSE TRUE
h
Count of Significant Peaks
EE RE
P15-45min 217 811 807 2392
P3.5-4hr 1 45 379 1739
P24hr 0 0 0 76
difference-in-changes
within-group
i
Distinct & Overlapping Significant Peaks
EE RE
P15-45min 9 208 603 32 775 1617
P3.5-4hr 0 1 44 34 345 1394
P24hr 0 0 0 0 0 76
difference-in-changes only
common
within-group only
